# Onco-fetal protein Nogo-A restricts human and mouse glioma vascularization and growth via VEGF-Notch-hippo-metabolic signaling

**DOI:** 10.1101/2024.10.02.616320

**Authors:** Marc Schwab, Moheb Ghobrial, Jorge Luis Jimenez-Macia, Michal Beffinger, Ji Yanrong, Andrin Wacker, Hidekiyo Harada, Jau-Ye Shiu, Oskar Nowitzki, Ignazio de Trizio, Francesco Girolamo, Mariella Errede, Tobias Weiss, Dorothée Gramatzki, Miguel Maurer, Oliver Weinman, Viola Vogel, Ivan Radovanovic, Martin E. Schwab, Nadia Dahmane, Jeffrey Greenfield, Antonio E. Chiocca, Daniela Virgintino, Michael Weller, Johannes Vom Berg, Johannes Vogel, Ramana V. Davuluri, Karl Frei, Ralph Schalbach, Katrien De Bock, Philippe P. Monnier, Sean E. Lawler, Thomas Wälchli

## Abstract

Glioblastoma is one of the most deadly human cancers characterized by high degrees of vascularization, but targeting its vasculature has resulted in very limited success so far. Angiogenesis, the growth of new blood vessels, is highly dynamic during brain development, enters a mostly quiescent state in the adult homeostatic brain, and is reactivated in vascular-dependent CNS diseases including brain tumors. In consequence, a better understanding of the relevance of the onco-fetal axis – describing the reactivation of fetal signaling programs in tumors – in endothelial– and perivascular cells of the human brain tumor vasculature harbors great translational potential, yet remains poorly defined. In development, neurovascular link (NVL) molecules guide both neuronal growth cones as well as capillary endothelial tip cells. Nogo-A is an NVL molecule known to inhibit axonal growth in the developing and adult CNS and to restrict angiogenesis during brain development, but its role in the mouse and human brain tumor vasculature along the onco-fetal axis remains unknown. Here, we characterize Nogo-A as an onco-fetal protein expressed in the neurovascular unit (NVU) in human fetal brains and human gliomas in vivo that negatively regulates sprouting angiogenesis and endothelial tip cells in glioma vascularization. The Nogo-A-specific Delta 20 domain restricts angiogenic sprouting and branching and promotes vascular normalization while inhibiting glioma growth in experimental gliomas. Moreover, Nogo-A expression in tumor cells negatively correlate/s with glioma malignancy in vivo. In vitro, Nogo-A Delta 20 reduced human brain– and brain tumor endothelial cell (HBMVEC, HBTMVEC) and human umbilical vein endothelial cell (HUVEC) spreading, migration, and sprouting, in a dose-dependent manner and inhibited filopodia extension glucose metabolism. Mechanistically, RNA sequencing of Nogo-A Delta 20-treated HBMVECs and HBTMVECs revealed Nogo-A Delta 20-induced positive regulation of the angiogenesis-inhibiting Dll4-Notch-pathway and inhibition of the angiogenesis-promoting VEGF-VEGFR and Hippo-YAP-TAZ pathways, whereas metabolomics and functional metabolic assays revealed Nogo-A Delta 20-induced negative regulation of endothelial glycolysis in HBMVECs and HBTMVECs. These findings characterize Nogo-A as an onco-fetal protein in the human glial brain tumor vasculature and identify Nogo-A Delta 20 signaling as an important negative regulator of human glioma vascularization and growth. Enhancing Nogo-A signaling may be an attractive alternative or combinatorial anti-angiogenic therapy to restrict human glioma/glioblastoma vascularization and growth.

## INTRODUCTION

Glial brain tumours are classified into lower-grade gliomas (WHO I-II) and higher-grade gliomas (WHO III-IV) and malignant transformation of lower-grade to higher-grade gliomas involves angiogenesis, the growth of new blood vessels^1–6^. It is well known that patients with glial brain tumors that harbor mutations in the isocitrate dehydrogenase 1 and 2 (IDH1 or IDH2) genes have better outcomes than those with wild-type IDH genes^7^. Interestingly, IDH mutations affect the tumor vasculature of low-and high-grade gliomas at both the morphological and molecular levels. For instance, IDH WT gliomas reveal an increased vessel density, a higher number of abnormal tumor blood vessels, hyper-proliferation of the vasculature, and the formation of larger glomeruloid vascular structures when compared to IDH mutant gliomas^8^. The molecular mechanisms underlying these observations remain, however, obscure. Glioblastoma (GBM) – the WHO grade IV glioma – represents the most prevalent and lethal malignant primary brain tumor^9^ and is among the most devastating human malignancies, with a median survival of less than 15-months and a 5-year survival of only 5%^10–12^. Despite standard treatment of care consisting in maximal safe surgical resection followed by chemo– and radiotherapy^12,13^, there is currently no efficient therapy prolonging survival after tumor recurrence. Typical features of GBMs include their high grade of vascularization established by angiogenesis^1–3^, as well as their extraordinary vessel abnormality^1412^. GBM vascularization and in consequence growth are driven by mutual interactions among the cellular components of the neurovascular unit (NVU)/perivascular niche (PVN) including endothelial– and perivascular cells (ECs and PVCs) such as pericytes, astrocytes, neurons, macrophages, microglia, and neuronal stem cells^15^. Therefore, therapies that target GBM angiogenesis and the GBM NVU have been tested^10,16–18^, but GBM tumors exhibit high resistance to both anti-angiogenic therapies and cytotoxic treatments. Accordingly, despite promising data in preclinical studies, anti-angiogenic therapies have not shown survival benefits in GBM patients in randomized controlled trials^10,16^, mainly owing to our limited knowledge regarding the cellular and molecular mechanisms regulating/driving angiogenesis/vascularization and the NVU in brain tumors^1,2,10,15,19,20^, as well as to an insufficient understanding of the differences between the healthy (fetal/developing and adult) and tumor vasculature.

Tumor blood vessels are abnormal, tortuous, and leaky and their architecture differs between brain and peripheral tumors^4,21^. During (glial) brain tumor progression, dysfunction of the tumor vasculature – which is mediated by dysregulated pro– and anti-angiogenic factors including vascular endothelial growth factor (VEGF) – results in hypoxia and an acidic microenvironment that, in turn, promotes tumor progression via hypoxia-inducible factor 1α (HIF1α)-induced transcriptional programs. Blocking VEGF signaling partially counteracts vascular dysfunction by transient pruning and by normalizing the immature and leaky brain tumor vessels for them to resemble the normal brain vasculature. Vascular normalization results in survival benefits in patients with newly diagnosed as well as with recurrent glioblastoma treated with antiangiogenic agents^4,25,26^. Importantly, however, adverse effects of high dose of anti-angiogenic therapy include resulting hypoxia, which increases migration and invasiveness of cancer cells^27^, as well as potential decrease of the blood-brain-barrier (BBB)/blood-tumor-barrier (BTB) permeability in glial brain tumors/glioblastoma^27^. Therefore, anti-angiogenic therapies that simultaneously inhibit brain tumor angiogenesis/vascularization while normalizing the brain tumor vasculature (thereby preventing hypoxia and tumor invasion), and that at the same time do not alter (but reconstitute) the blood-brain-barrier properties are/represent ideal therapeutic strategies. Notably, molecular cues affecting angiogenesis and barriergenesis are at play during brain development^19,21,28–30^ and can benefit our understanding of the brain tumor vasculature. For instance, as we have previously described, CNS-specific cues for angiogenesis regulate both angiogenesis and the formation and differentiation of the BBB, whereas general cues for angiogenesis do not^19,21,31^, highlighting the tight link between CNS-specific angiogenesis and barriergenesis across development and disease.

Interestingly, whereas brain vascularization/angiogenesis is highly dynamic during brain development, it is mostly quiescent in the adult brain with very little proliferating endothelial cells (ECs), thereby ensuring a stable blood-brain barrier (BBB)^19,21,32–34^. Brain vascularization/angiogenesis and the NVU are reactivated in various angiogenesis-dependent central nervous system (CNS) pathologies including brain tumors, brain vascular malformations, and stroke^1,4,15,19,21,33^. However, whether there is molecular similarity between developmental– and (brain) tumor angiogenesis (termed onco-fetal axis^35–42^, see below), and how neurodevelopmental pathways regulate brain tumor (vessel) growth remains poorly defined. Accordingly, a better understanding of pathological brain tumor vasculature requires a molecular understanding of normal vascular brain development^4,19,32^.

During development, the brain vascular network is established in a two-step process: i) during embryogenesis, the perineural vascular plexus (PNVP) surrounding the CNS forms via vasculogenesis (= de novo formation of blood vessels from angioblasts); ii) at the postnatal stage^19,21,43,44^, the intraneural vascular plexus (INVP) within the brain parenchyma is formed by sprouting angiogenesis (= formation of new blood vessels from pre-existing ones)^19,21,43,45^. Sprouting angiogenesis refers to endothelial tip cells (ETCs) and their filopodia at the forefront of vascular sprouts that guide the growing vessels^19,46,47^, to endothelial stalk cells that proliferate thereby elongating the vascular sprout^46,48^, and to quiescent endothelial phalanx cells that line the parental vessel^33^. Once neighboring sprouts have fused, pericytes and perivascular astrocytes stabilize the vessel sprout and facilitate the formation of functional BBB microvessels^15,16,19,20^. ETCs drive sprouting angiogenesis during both development and in tumors^1,2,46,49,50^ and interact with tumor cells within the perivascular niche/NVU in non-CNS/peripheral tumors^51,52^ Peri-/neurovascular crosstalk and metabolism are typical features of brain tumors^1,2,13,15,16,50,53–57^, but little is known about how developmental signaling axes are reactivated in brain tumors to regulate angiogenesis/vascularization/ and vascular endothelial metabolism^1,16,53,54^.

The onco-fetal axis is defined as the reactivation of fetal signaling programs in tumor tissues and has been described in cancer cells in brain and peripheral tumors^35–41^ as well as in endothelial cells in liver cancer^42^ but its relevance in endothelial– and perivascular cells of the brain vasculature in human (glial) brain tumors remains poorly understood. Because onco-fetal programs represent interesting therapeutic targets given their upregulation within the brain tumor tissue as compared to the surrounding healthy brain (thereby minimizing the likelihood of detrimental side-effects^28^), a better understanding of the onco-fetal axis in the (glial) brain tumor vasculature harbors great scientific and translational potential.

The onco-fetal axis includes molecules of the neurovascular link (NVL), which are attractive and repulsive molecular guidance cues that are shared between growing blood vessel ETCs and growing axonal growth cones, and can be CNS-specific or general regulators of vessel growth^1921^.

Nogo-A is a high molecular weight membrane protein expressed on the surface of oligodendrocytes and neurons with important inhibitory functions for/on neurite growth, axonal regeneration and synaptic plasticity in the adult CNS and acting as a repulsive axonal guidance molecule during development of the CNS as well as of the peripheral nervous system. We have identified an additional function of Nogo-A as a negative regulator of angiogenesis, endothelial tip cell formation and 3D vascular network architecture in the postnatal CNS in vivo and in vitro^30,58^ thereby characterizing Nogo-A as a NVL molecule and a general/non-CNS specific cue for angiogenesis^19^.

Nogo-A exerts its inhibitory effects via two functional domains: the Nogo-A specific fragment Nogo-A Delta 20 (rat amino acids 544 – 725), and the Nogo-66 sequence (rat amino acids 1019-1083), which is present in in all three Nogo isoforms Nogo-A, –B and –C^59,60^. The Nogo-A fragment Delta 21 serves as control peptide for Nogo-A Delta 20^30,59,60^, Both Nogo-A Delta 20 and Nogo-66 bind to several domain specific, signal transducing multimeric receptor complexes in neurons^61,62^. The known functional receptors for the Nogo-66 domain are Nogo receptor 1 (NgR1), the co-receptor paired immunoglobin-like receptor B (PirB) the adaptor element low density lipoprotein receptor (LRP), and the effector part p75/TROY^63^. Only recently, sphingosine-1-phosphate receptor 2 (S1PR2), co-receptor tetraspanin 3 (TSPAN3) and syndecan 3/4 (SDC3/4) have been identified as receptors for the Nogo-A Delta 20 domain^61–64^. Nogo-A Delta 20 but not Nogo-66 exert inhibitory effects on developmental CNS angiogenesis^30^. Although no function has been ascribed to S1PR2, TSPAN3, and SDC4 during developmental angiogenesis so far, there is evidence that S1PR2, TSPAN3 and SDC4 play a role in neo-angiogenesis^65–67^. S1PR2, TSPAN3 and SDC4 are expressed on normal and tumor blood vessels in many organs of adult mice^65^.

Given the negative regulatory roles of Nogo-A and S1PR2, TSPAN3, and SDC4 on angiogenesis^30,65,66^ and their mutual interaction in neurons^61^, it is tempting to speculate that Nogo-A Delta 20 and S1PR2/TSPAN3/SDC4 constitute a novel ligand-receptors complex for developmental and tumor angiogenesis in the CNS.

In brain tumors, Nogo-A has been proposed as a marker allowing to differentiate between the various types of human gliomas, with a higher Nogo-A expression in oligodendrogliomas than in the more densely vascularized glioblastomas^68^. Furthermore, it was reported that Nogo-A expression negatively correlated with the malignancy grade of oligodendrogliomas^69^, where Nogo-A might restrict oligodendroglioma vascularization and thereby tumor growth. Another study has shown that the splicing factor polypyrimidine tract binding protein 1 (PTBP1) repressed Nogo-A protein translation and increased the proliferation of human glioma cell lines, but Nogo-A overexpression slowed glioma cell proliferation, suggesting that PTBP1 may – in addition to exerting a cell-autonomous effect on tumor cell division – control glioblastoma vascularization by regulating Nogo-A expression in glioma cells. These findings suggest inhibitory roles for Nogo-A on glioma/glioblastoma cell proliferation and migration. Glioblastomas invade the brain by infiltrating into the CNS white matter along myelinated nerve fiber tracts that express Nogo-A which prevents tumor cell migration via activation of inhibitory RhoA signaling. Glioblastoma cells are capable of secreting Secreted protein acidic and rich in cysteine (SPARC) that inhibits Nogo-A-RhoA mediated tumor cell invasion of white matter structures resulting in improved response to cytostatic therapy and prolonged survival^70^. Moreover, the Nogo-66 receptor NgR1 also inhibits myelin-associated infiltration of glioblastoma cells into the CNS white matter and NgR1 downregulation is associated with highly infiltrative characteristics and poor survival of glioblastoma patients^71^. Despite these observations, however, expression of oligodendroglial markers including Nogo-A and Olig2 did not reveal independent prognostic effect in IDH-wildtype glioblastoma in a tissue micro array (TMA) dataset^72^, whereas another study indicated the potential usefulness of myelin-associated proteins, notably cerebrospinal fluid (CSF) Nogo-A and serum MAG as circulating biomarkers of primary (glial) brain tumors^73^.

However, whether Nogo-A acts as an onco-fetal protein as well as its role on angiogenesis and endothelial cell function in human gliomas remains unknown. Here, using a variety of in vivo and in vitro assays, we characterize Nogo-A as an onco-fetal protein in human glioma vascularization that negatively regulates/restricts sprouting angiogenesis and endothelial metabolism via its S1PR2/TSPAN3/SDC4 receptor complex in human and mouse glioma and molecular crosstalk with the key angiogenic pathways VEGF-VEGFR, Dll4-Jagged-NOTCH, and Hippo-YAP-TAZ, and find that its endogenous expression negatively correlates with human glial brain tumor vascular density and positively associates/correlates with IDH-WT low-grade glioma patient survival.

## RESULTS

### Nogo-A is an onco-fetal protein expressed in vicinity of blood vessel endothelial (tip) cells in the human fetal brain and in the NVU/PVN of human glial brain tumors

In both humans and mice, the brain is predominantly vascularized by sprouting angiogenesis guided by endothelial tip cell filopodia during fetal/embryonic and postnatal brain development^19,21,29^. Nogo-A expression and neuronal function has been described during mouse embryogenesis^60^ but whether it is expressed in the NVU/PVN of the human fetal brain and of human glial brain tumors has not been addressed so far. To investigate whether Nogo-A constitutes an onco-fetal protein that is reactivated in brain tumors, we performed immunofluorescence microscopy of the main NVU/PVN cellular components of human fetal brain and of human glial brain tumors. Nogo-A was expressed during fetal forebrain neocortex development, at gestational week 22 (GW22) and upregulated in brain tumors, as revealed by immunofluorescence staining against Nogo-A and the nuclear marker TO-PRO-3^74^ (Supplementary Figure 1a-f). Nogo-A was expressed in vicinity of. angiogenic brain endothelial (tip) cells (ETCs) labeled with the endothelial marker cluster of differentiation 105 (CD105, (Supplementary Figure 1a-c) and in perivascular cells including Glial fibrillary acidic protein (GFAP)^+^ astrocytes during brain development, (Supplementary Figure 1d-f), reminiscent of its expression pattern in the postnatal mouse brain cortex^30^, and suggesting that migrating endothelial (tip) cells of the developing human CNS vasculature contact the Nogo-A containing parenchyma. Sphingosine-1-Phosphate Receptor 2 (S1PR2), a component of the putative Nogo-A receptor complex S1PR2/TSPAN3/SDC4 ^61,62,64^ was also highly expressed in CD105^+^ angiogenic ECs during fetal brain development (Figure 1g,h).

**Figure 1:**
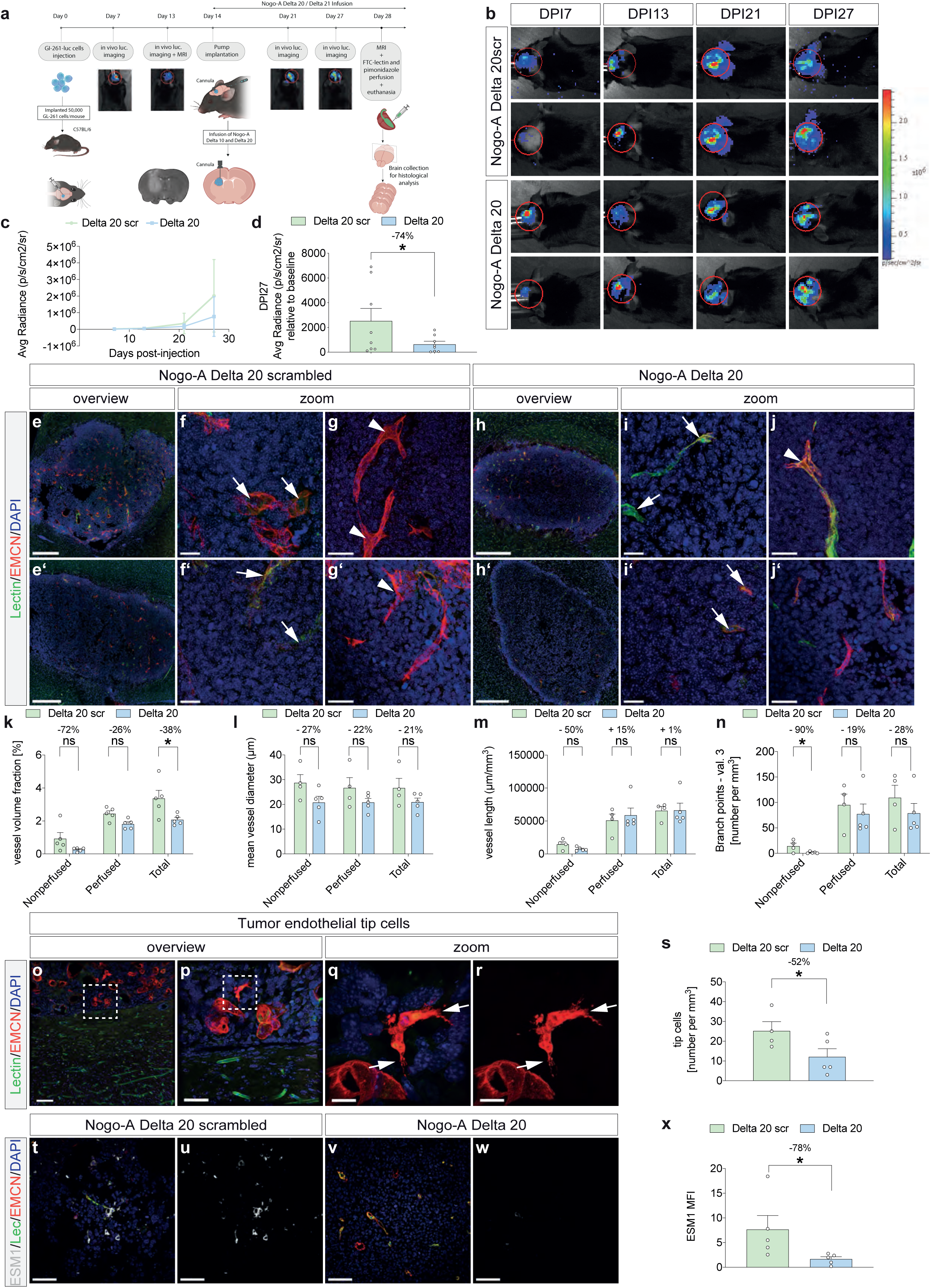
Local infusion of Nogo-A Delta 20 inhibits tumor growth and decreases the vascular volume fraction as well as the number of endothelial tip cells of growing blood vessels in the orthotopic GL-261 glioma model. (**a**) In an orthotopic GL-261 mouse GBM model, we injected luciferase expressing GL-261, performed in vivo bioluminescence measurement and MRI in order to; i) randomize the mice before the beginning of the treatment to have the same mean tumor size in both the treatment and the control groups and ii) monitor tumor growth. Nogo-A Delta 20 and Nogo-A Delta 20 scrambled (inactive form of Nogo-A Delta 20) were infused from days post injection (DPIs) 14 to 28 into the tumor by osmotic minipumps. Luciferase bioluminescence was measured at DPIs 7, 13 (day of randomization of mice), 21, and 27, while T2 weighted (T2w) MRI was performed on DPIs 14 and 28. Before euthanasia, mice were perfused with FITC-lectin in order to visualize blood vessel perfusion in the glial brain tumors. (**b**-**d**), Intracranial astrocytoma/glioma growth is reduced in Nogo-A Delta 20 treated mice compared with Delta 20 scrambled treated control littermates (n=8 mice per group).(**e-j’)** Coronal brain sections (40 µm) of glioblastoma-bearing mice (28 days post-inoculation, DPI28) which were perfused with FITC Lectin (green) to visualize perfused blood vessels and stained for endomucin to visualize blood vessels including endothelial tip cells (red) and the nuclear marker DAPI (blue). FITC Lectin_+_ established blood vessels (green) are shown within the glioblastoma and in the tumor-surrounding healthy brain parenchyme whereas endomucin^+^ endothelial (tip) cells (red) are present within the tumor but almost completely absent in the tumor-surrounding healthy brain tissue. (**k)** Vessel volume fraction of perfused, nonperfused and total vessels. In Delta 20 treated tumors, ∼2.1% of the tissue is occupied by vessel structures, of which ∼87.5% are perfused as compared to control treatment where ∼3.4% of the volume is occupied by vessels, of which ∼72.5% are perfused. Nogo-A Delta 20 infusion significantly decreases the vascular volume fraction of perfused and total vessel structures (n = 4-5 mice per group). (**l**) Vessel diameter of perfused, nonperfused and total vessels. Tumor blood vessels tend to show smaller diameters in the Nogo-A Delta 20 treated mice (∼20.8μm for non-perfused, perfused, and total vessels) as compared to the control group (∼28.7μm for non-perfused vessels, and ∼26.7μm for perfused and total vessels) (n = 4-5 mice per group). (**m**) Vessel length of perfused and total vessels remain unchanged upon Nogo-A Delta 20 treatment. However, nonperfused vessels tend to be shorter in Nogo-A Delta 20-treated mice (∼7450μm/mm^3^ of tissue for Nogo-A Delta 20 treated mice, and ∼14700μm/mm^3^ of tissue for Nogo-A Delta 20 scrambled treated mice) (n = 4-5 mice per group). (**n**) Vessel branching was significantly reduced in nonperfused vessels following Nogo-A Delta 20 treatment as compared to Nogo-A Delta 20 scrambled treatment. Perfused and total vessels tend to have less branch points of valence 3 (three vessel branches emanating from the vessel branch point, n = 4-5 mice per group). (**o**-**r**) Endomucin^+^ endothelial tip cells (red, arrows) and FITC Lectin^+^ established blood vessels (green) are shown within the glioblastoma. (**a,f-n**) Endomucin seems to be specific for tumor blood vessels (red) (as compared to endomucin^−^ blood vessels in the surrounding normal brain). (**o**-**r**) Note the nicely visualized endomucin^+^ endothelial tip cells including filopodial protrusions (red, arrows) (**q**,**r**) within the glioma. The boxed areas are enlarged in **o**-**p** and in **q**-**r**. (**s**) Infusion of Nogo-A Delta 20 significantly decreases the mean number of endothelial tip cells within the tumor tissue, showing its inhibitory effect on mouse glioma blood vessels (n = 4-5 mice per group). (**t-w**) Coronal brain sections were stained for the endothelial tip cell marker ESM-1. (**x**) Nogo-A Delta 20 infusion significantly decreases ESM-1 expression in mouse gliomas as revealed by mean fluorescence intensity (MFI) analysis (n = 4-5 mice per group). Data represent mean ± SEM. For statistical analysis, two-tailed unpaired Student’s t –test (**d**, **s**,**x**,**ci**) and two-way ANOVA with Sidak’s multiple comparison test comparing treatment columns (**k**-**n**) were performed. \**P* < 0.05. Scale bars: 500 µm in **e**,**e’**,**h**,**h’**; 20 µm in **f**,**f’**,**i**,**i’**; 40 µm in **g**,**g’**,**j**,**j’**,**p**; 10 µm in **q**-**r**, 70 µm in **t**-**w**.

Within the physiological and pathological NVU, blood vessel endothelial cells are in contact with perivascular supportive cells such as pericytes, astrocytes, and neuronal stem cells^19,21,29,75–78^. We therefore assessed Nogo-A expression in perivascular astrocytes in human GBM. Glial fibrillary acidic protein (GFAP)^+^ astrocytes formed typical patterns by contacting ECs and revealed Nogo-A expression in glioblastoma with GFAP^+^/Nogo-A^+^ astrocytes (Figure 1e,f). However, Nogo-A appeared to be upregulated in human glioblastoma astrocytes (Figure 1e,f).

Together, these data suggest that Nogo-A is expressed in close proximity of endothelial (tip) cells during fetal brain development, and in astrocytes in GBM, thereby characterizing Nogo-A as an onco-fetal protein in the human brain vasculature.

### Nogo-A ablation increases the number of endothelial tip cells and their filopodia but does not affect the formation of the blood-brain-barrier in the postnatal brain

In postnatal day 8 (P8) mouse forebrain cortices, S1PR2 was expressed in the brain tissue and on Isolectin B4 (IB4)^+^ blood vessels (Supplementary Figure 2a-c), indicating that, similarly to the situation in the human fetal brain, vascular endothelial cells and their growing tips express the Nogo-A receptor S1PR2 in the postnatal mouse brain where they have a high chance to encounter Nogo-A expressing neurons^30^. Next, we analyzed whether genetic ablation of Nogo-A and S1PR2 affected endothelial tip cells at the leading edge of growing vessel sprouts^19,21,29^. As previously observed^30^, IB4^+^ ETCs with their typical protruding filopodia were recognized in the P8 brain cortex. In P8 Nogo-A^−/−^ and S1PR2^−/−^ cortices^30,61^ (Supplementary Figure 2d-f), stereological analysis revealed that the number of ETCs/mm3 was significantly increased (by 49% and 42%, respectively) as compared to the P8 WT control mice (Supplementary Figure 2g). Further, whereas the number of tip cell filopodia/ETC was not changed between the groups, filopodia length was significantly increased in P8 Nogo-A^−/−^ and S1PR2^−/−^ mice, filopodia straightness/ETC was decreased in Nogo-A^−/−^ and S1PR2^−/−^mice as compared to their WT littermates (Supplementary Figure 2h-m).

Next, to determine whether Nogo-A influenced the formation of the blood-brain-barrier^19,21^, we investigated the effect of Nogo-A^−/−^ on the immature, leaky blood-brain barrier in the first postnatal week^79^, by intracardial injection of Evans blue^80^. Evans blue extravasation was not significantly changed between the P8 Nogo-A-/– and WT mice. (Extended Data Figure 2b,d), indicating that Nogo-A does not affect blood-brain-barrier formation and leakiness at this postnatal stage of mouse brain development.

Together, these results indicate negative regulatory effects of Nogo-A on postnatal CNS angiogenic ETCs and their filopodia (in agreement with Wälchli et al., PNAS, 2013), while not affecting barriergenesis, thereby confirming the previously suggested role of Nogo-A as a general mechanism of angiogenesis (Wälchli et al., Neuron, 2015).

### Nogo-A Delta 20 infusion restricts tumor vascularization and tumor growth in experimental mouse-in-mouse glioblastoma

Based on our previous observations showing that Nogo-A gene deletion leads to hypervascularization and additional functional blood vessels in the postnatal mouse brain in vivo^30,58^ and the observed strong inhibitory effects of Nogo-A Delta 20 on brain developmental angiogenesis and endothelial cells in vitro^30^, we tested whether local delivery/topical application of Nogo-A Delta 20 might affect a growing vascular bed in vivo. Therefore, we applied Nogo-A Delta 20, Nogo-A Delta 21 and Nogo-66 peptides on the ex vivo Chicken Chorioallantoic Membrane (CAM) angiogenesis assay^81,82^ (Supplementary Figure 3a-k). Qualitatively, CAMs at Incubation Day 12 (ID12) upon topical application of Nogo-A Delta 20 and Nogo-66 revealed a decreased density of CAM vessels as compared to the Nogo-A Delta 21 and PBS control groups (Supplementary Figure 3d-g). Notably, we observed that Nogo-A Delta 20 and Nogo-66 mainly affected the small caliber vessels, presumably the capillary bed (Supplementary Figure 3d-g), in agreement with our previous observations in Nogo-A^−/−^ animals in the postnatal mouse brain^30,58^. To quantitatively assess these differences, we next determined vessel density and angiogenesis (inhibition) by measuring blood flux on the CAMs at ID12.5 (Supplementary Figure 3a,h-k). The blood flux in Nogo-A Delta 20 and Nogo-66 treated CAMs was significantly decreased as compared with the control groups, with the strongest inhibitory effect again observed at the level of capillaries (Supplementary Figure 3b,d-g).

As VEGFA-VEGFR is a major pro-angiogenic and permeability signaling axis which is highly expressed during brain development and in brain tumors and thought to mainly drive angiogenesis/neovascularization of brain tumors and particularly of gliomas^3–5^, we next determined whether Nogo-A inhibitory peptides were capable of counteracting VEGF-A driven angiogenesis.

While VEGF-A significantly increased CAM vessel density/blood flux (Supplementary Figure 3c,h), Nogo-A Delta 20 and Nogo-66 significantly counteracted the pro-angiogenic effects of VEGF-A at ID12.5 (Supplementary Figure 3c,j,k), indicating that Nogo-A peptides are capable of restricting VEGF-A-VEGFR driven angiogenesis.

Next, to test whether S1PR2 is involved in the negative regulatory effect of Nogo-A Delta 20 on the growing CAM vasculature, we applied Nogo-A Delta 20 on CAM membranes incubated with JTE-013. Blockade of S1PR2 using JTE-013 significantly counteracted the inhibitory effects of Nogo-A Delta 20 on CAM angiogenesis (Supplementary Figure 4b,g).

Given that the highest decrease of blood flux (as a surrogate for vessel density and angiogenesis) in response to Nogo-A inhibitory peptides was found for Nogo-A Delta 20, we used the Nogo-A Delta 20 for all subsequent experiments.

Next, we addressed whether intratumoral delivery of Nogo-A Delta 20/Delta 21/Delta 20 scrambled affects the growth and vasculature of glial brain tumours. We referred to the GL261 glioma mouse model which possesses numerous features reminiscent of human gliomas, including abnormal vasculature, high VEGFA production and oedema^4,31,83–85^. Orthotopic intracerebral injection of Luciferase-expressing syngeneic GL261 glioblastoma (GBM) cells^83,86,87^ into the right medial striatum of adult (6-10 weeks old) C57BL/6 wild-type (WT) mice was performed (Figure 1a).

We examined how intratumoral delivery of Nogo-A Delta 20 and the control Delta 20 scrambled impacted the GL-261 glioma growth and angiogenesis/vasculature. Therefore, we used osmotic minipumps filled with 100uM Nogo-A Delta 20/ Delta 20 scrambled implanted on the back of animals (Figure 1a and see paragraph “methods”). We first assessed intracranial tumor growth in Nogo-A Delta 20 versus Nogo-A Delta 20 scrambled infused mice in the orthotopic glioma tumor model. Intracranial tumor growth in Nogo-A Delta 20 treated mice was significantly reduced, reaching less than 30% of the volume of control tumors in Delta 20 scrambled infused mice at day 13 of treatment (DPT13) (corresponding to day 27 post cell inoculation (DPI27)), as revealed by bioluminescence analysis of Luc^+^ tumor cells (Figure 1b-d). Magnetic resonance imaging (MRI) confirmed the reduction of tumor volumes in Nogo-A Delta 20 treated mice as compared to control mice infused with Nogo-A Delta 21 at DPT14 (corresponding to day 28 post cell inoculation, Supplementary Figure 5a,b), and the decreased tumor volume was likely not influenced by edema, as this parameter was very low and very similar in the two groups (Supplementary Figure 5a).

To determine the in vivo role of Nogo-A in/on tumor vascularization, we investigated the effects of intratumoral Nogo-A Delta 20 on vascular morphology and sprouting angiogenesis in mouse GBMs. The reduced tumor growth was associated with a decreased tumor vascularization, as reflected by stereological quantification of the vascular network in mouse brain tumors using an adapted version of our previously published method in the developing postnatal mouse brain^29^ (Figure 1e-x). The total, perfused and non-perfused vascular network in the mouse brain tumors was analyzed using a combination of Endomucin (EMCN) immunofluorescence to visualize the total vessel tree and intravascular Fluorescein-5-isothiocyanat (FITC) Lectin injection to label perfused vessels^88^, thereby allowing simultaneous assessment of the total Endomucin^+^ vascular network including newly formed vessel sprouts with endothelial tip cells (vascular sprouts are FITC-Lectin^+^/Endomucin^+^ or FITC-Lectin^−^/Endomucin^+^) as well as FITC-Lectin^+^/Endomucin^+^ perfused functional blood vessels (Figure 1e-x). Morphologically, the vasculature of control Nogo-A Delta 21/Delta 20 scrambled infused tumors consisted of irregular, tortuous and highly branched blood vessels (Figure 1e,f,g), whereas, in contrast, the vasculature of the tumors grown in Nogo-A Delta 20 infused mice was more regular, less tortuous and less branched, resulting in a well-organized structure of the tumor vasculature (Figure 1h-j). Using design-based stereological methods, we provide absolute parameters characterizing the mouse brain tumor vessel network. The vascular volume fraction of Endomucin^+^ total-, FITC-Lectin^+^/Endomucin^+^ perfused– and FITC-Lectin^−^/Endomucin^+^ non-perfused tumor blood vessels was significantly reduced in the Nogo-A-Delta 20 treated animals, by 38%, 26%, and 72%, respectively (Figure 1k), indicating that the decreased tumor vascular volume density/fraction is accompanied by a similar decrease in functional vascular volume fraction/vessel perfusion. Stereological analysis revealed a vascular volume fraction of 3.37% (2.45% perfused, 0.93% non-perfused) in the Nogo-A Delta 20 scrambled infused mouse brain tumor tissue and of 2.08% (1.82% perfused, 0.26% non-perfused) in the Nogo-A Delta 20 treated mice (Figure 1k), which is similar/in accordance with previous findings in the postnatal mouse brain^29,58^. Moreover, the significantly decreased tumor vessel density in Nogo-A treated mice was accompanied by changes in tumor vessel morphology, notably a non-significant/slight decrease in the mean vessel diameter (Figure 1l), whereas vessel length did not show significant changes (Figure 1m). The reduction in vascular volume fraction was further reflected by a decrease in the number of vessel branchpoints of valence 3 of total, perfused, and non-perfused tumor vessels (reduction of 28% (not significant), 19% (not significant), and 90%, respectively, Figure 1n). While we observed a reduction in the number of branchpoints of both the non-perfused and perfused tumor vessels, the main effect was at the level of non-perfused vessels, suggesting the regulatory inhibitory effect of Nogo-A Delta 20 on sprouting angiogenesis (resulting in vascular/vessel branchpoints) in the forming tumor vascular bed (Figure 1n). Given the inhibitory effect of Nogo-A Delta 20 on (capillary) tumor vessels and vessel sprouting/branching, we next investigated whether Nogo-A Delta 20 affected ETCs. Therefore, we developed an adapted protocol of our previously described methodology in the postnatal mouse brain^29^ using a combination of FITC-Lectin perfusion and Endomucin staining which enabled the visualization of Endomucin^+^ endothelial tip cells with its typical, finger-like protrusions within the brain tumor, (Figure 1p-r). Indeed, vascular sprouts, endothelial tip cells and filopodial extensions were clearly detectable on the blood vessels of control and Nogo-A Delta 20 treated brain tumors (Figure 1o-r, higher magnifications). Importantly, in accordance with the observed reduction/inhibition in vascular sprouting upon Nogo-A Delta 20 infusion (Figure. 1e–n), tumor blood vessels in Nogo-A Delta 20 treated mice displayed fewer sprouts and filopodia (Figure 1o-r, higher magnifications), indicating inhibitory effects of Nogo-A Delta 20 signaling on tumor blood vessel endothelial tip cells. Quantitatively, Nogo-A Delta 20 significantly decreased the number of Endomucin^+^ ETCs and the mean fluorescence intensity of Endothelial cell specific molecule 1 (ESM-1, a suggested tip cell marker) in the GL-261 mouse tumors (Figure 1s). Stereological analysis revealed 25,21 endothelial tip cells per mm^3^ in the Nogo-A Delta 20 scrambled and 12,12 endothelial tip cells in the Nogo-A Delta 20 conditions (Figure 1s).

Notably, given that the used/utilized design-based stereological method provides absolute parameters characterizing the 3D mouse brain glioma vessel network, the values of non-perfused, perfused and total vascular volume fraction the control groups were very similar to the ones we previously reported in the postnatal mouse brain (Wälchli et al., Nat Protoc, 2015), highlighting the similar active ongoing angiogenesis in developing brains and in brain tumors^21^.

Taken together, these data indicate inhibitory effects of Nogo-A Delta 20 on tumor vascularization and ETCs, tumor vessel architecture, and tumor growth.

### Nogo-A Delta 20 infusion promotes tumor vessel normalization and reduces tumor microenvironment hypoxia in experimental mouse-in-mouse glioblastoma

Because anti-angiogenic therapy can result in hypoxia in the tumor neurovascular unit/microenvironment which increases invasiveness of cancer cells^4,27^, and fuels brain tumour progression and vascular dysfunction (e.g. tortuous tumor vessels with abnormal architecture) via hypoxia-inducible factor 1α (HIF1α)-driven transcriptional programs ^24,89^, and because vessel normalization prunes immature brain tumour vessels in mice, alleviates hypoxia and results in survival benefits in glioblastoma patients (newly diagnosed and recurrent) receiving antiangiogenic agents^4,25,26^, we next tested whether intratumoral delivery of Nogo-A Delta 20 affected vessel normalization and intratumoral hypoxia. The percentage of perfused vessels as revealed by the ratio of FITC-Lectin^+^/Endomucin^+^ blood vessels (labeling perfused vessels) and Endomucin^+^ blood vessels (identifying all tumor vessels) showed an increase, although not significant, in Nogo-A Delta 20 treated mice (Figure 2a-d,i) but qualitative exploration of structural properties of tumor blood vessels in FITC-Lectin/Endomucin stained tumor sections revealed that the vascular architecture in the Nogo-A Delta 20-treated GL-261 tumors was less irregular, less tortuous, and less organized (Figures 2a-h). Overall, the glioma vasculature in the Nogo-A Delta 20 mice was more organized than in the control mice and appeared normalized (Figure 2a-x).

**Figure 2:**
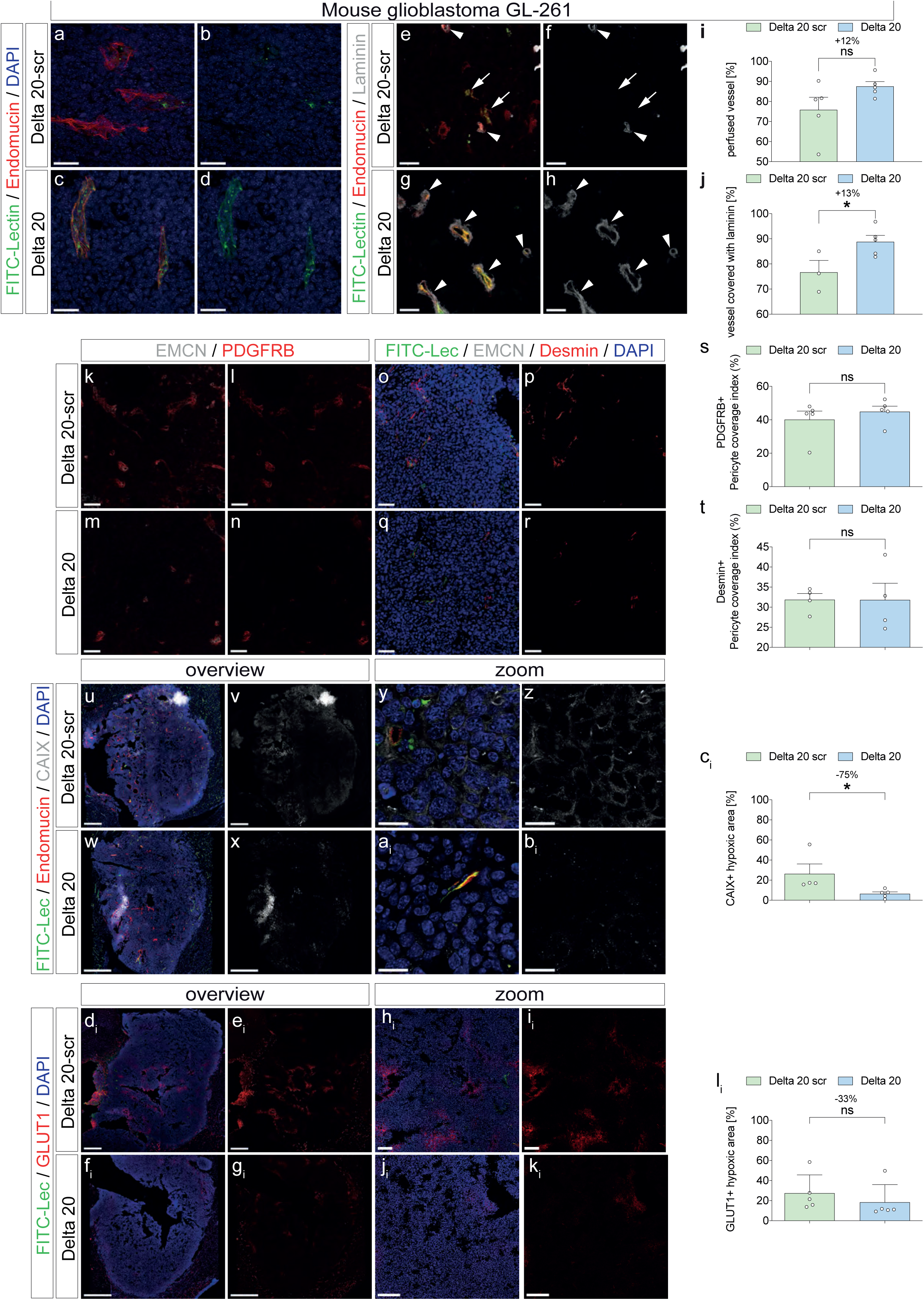
Infusion of Nogo-A Delta 20 induces vessel normalization and decreases hypoxia in the mouse orthotopic GL-261 glioma model. Nogo-A Delta 20 and Nogo-A Delta 20 scrambled (inactive form of Nogo-A Delta 20) were infused into the tumor by osmotic minipumps. Glioblastoma-bearing mice (28 days post-injection, DPI28) were perfused with FITC Lectin (green) to visualize perfused blood vessels on coronal brain sections (40 μm), which were then stained for endomucin to visualize blood vessels (red), laminin (white, **e**-**h**), Pdgfr– and desmin to visualize pericytes (red, **k-r**) the β Carbonic Anhydrase IX and Glucose transporter 1 (CAIX, GLUT1, white, **u**-**k**_i_) to detect the hypoxic area. Cell nuclei were stained using the general nuclear marker DAPI (blue). (**i)** In Delta 20 treated tumors, ∼87.5% of the vasculature is perfused as compared to control treatment where ∼72.5% is perfused. Nogo-A Delta 20 infusion significantly decreases the percentage of perfused vessels. (**j)** Delta 20 infusion significantly increased laminin deposition around vessels as compared to Delta 20 scrambled control. (**s**,**t**) Quantification of pericyte coverage (Pdgfr-+ and desmin+, showed no difference in pericyte coverage between β the two conditions (*n* = 4-5 mice per group). (**u-c**_i_) Nogo-A Delta 20 significantly decreases the CAIX positive area (**c**_i_), as compared to the control group Nogo-A Delta 20 scrambled. (**d**_i_-**l**_i_) As another indication of decreased tumor hypoxia, Glut-1 positive area was decreased upon Nogo-A Delta 20 treatment (*n* = 4-5 mice per group). Data represent mean ± SEM. For statistical analysis, two-tailed unpaired Student’s t –test were performed. \**P* < 0.05. Scale bars: 40 µm in **a**-**d**; 20 µm in **e**-**h**; 400 µm in **k**-**n**; 10 µm in **o**-**r**; 500 µm in **t**-**w**; 100 µm in **x**-**a**_i_; 50 µm in **c**_i_-**j**_i_;.

As tumor vessels are unstable and characterized by an incomplete or absent basement membrane^90^, we next assessed the fraction of tumor vessels with a laminin-positive vascular basement membrane. Tumors in Nogo-A Delta 20 infused mice contained significantly more mature Endomucin^+^/Laminin^+^, FITC-Lectin^+^/Endomucin^+^/Laminin^+^ or FITC-Lectin^+^/Laminin^+^ vessels (Figure 2e-h,j), indicating improved vessel maturation. Interestingly, while the coverage of FITC-Lectin^+^/Endomucin^+^ blood vessels with the vascular basement membrane markers Laminin was significantly increased, vessel coverage with the pericyte markers PDGFRB and Desmin did not reveal significant changes in the Nogo-A Delta 20-treated animals (Figure 2k-t).

Vascular normalization is thought to reduce hypoxia thereby attenuating pro-angiogenic stimuli^1,3,4,53,90,91^. Thus, we next assessed whether the observed vascular normalization alleviates hypoxia (Jain, Science, 2005, Tian et al., Nature, 2015, Viallar and Larrivée, Angiogenesis, 2017). Immunofluorescence staining using carbonic anhydrase IX (CAIX) and GLUT1 for hypoxia confirmed that the Nogo-A Delta 20-induced vascular normalization resulted in decreased hypoxia (Figure 2u-b_i_,d_i_-k_i_). Strikingly, intratumoral hypoxia, as revealed by CAIX (significantly) and GLUT1 (not significantly) stainings, was decreased in these Nogo-A Delta 20 treated mice (Figure2c_i_,l_i_) Taken together, these data show that Nogo-A Delta 20 gain-of-function exerts strong inhibitory effects on glioma vessel– and tumor growth while inducing tumor vessel normalization of the altered morphology of the tortuous and tumor vessels resulting in a more well-organized structure of the tumor vasculature and in consequence to a marked reduction in intratumoral hypoxia, suggesting reconditioning of the brain tumor microenvironment.

### Nogo-A Delta 20 infusion restricts tumor growth in experimental human-in-mouse xenograft glioblastoma

Next, to further test the effects of Nogo-A Delta 20-mediated anti-angiogenic therapy in a setting that more closely recapitulates human GBM, we implanted human patient-derived G30-LRP glioblastoma cells in naïve adult athymic nude mice (Jimenez-Macia et al., 2022), as the G30-LRP glioblastoma tumor model recapitulates growth kinetics of human glioblastoma (Jimenez-Macia et al., 2022) (Figure 3a). Next, we examined how intratumoral delivery of Nogo-A Delta 20/Delta 21 affected the G30-LRP xenogeneic glioma growth. Similarly to the GL-261 model, we implanted osmotic minipumps filled with 150uM Nogo-A Delta 20/Nogo-A Delta 21 on the back of animals (Figure 3a and see paragraph “methods”). We assessed intracranial tumor growth in Nogo-A Delta 20 versus Nogo-A Delta 21 infused mice in the orthotopic human-in-mouse glioma tumor model (Jimenez-Macia et al., 2022) and observed that intracranial tumor growth in Nogo-A Delta 20 treated mice was significantly reduced, reaching 52% of the volume of control tumors in Delta 21 infused mice (Figure 3b-e) at day 9 of treatment (DPT9) (corresponding to day 22 post cell inoculation (DPI22)), as revealed by bioluminescence analysis of Luc^+^ tumor cells (Figure 1b-d). Moreover, tumor volume (as quantified by DAPI^+^ tumor cell volume) in Nogo-A Delta 20 treated mice were/was significantly reduced (by 68%) as compared to those in the control mice infused with Nogo-A Delta 21 (Figure 3f),

**Figure 3:**
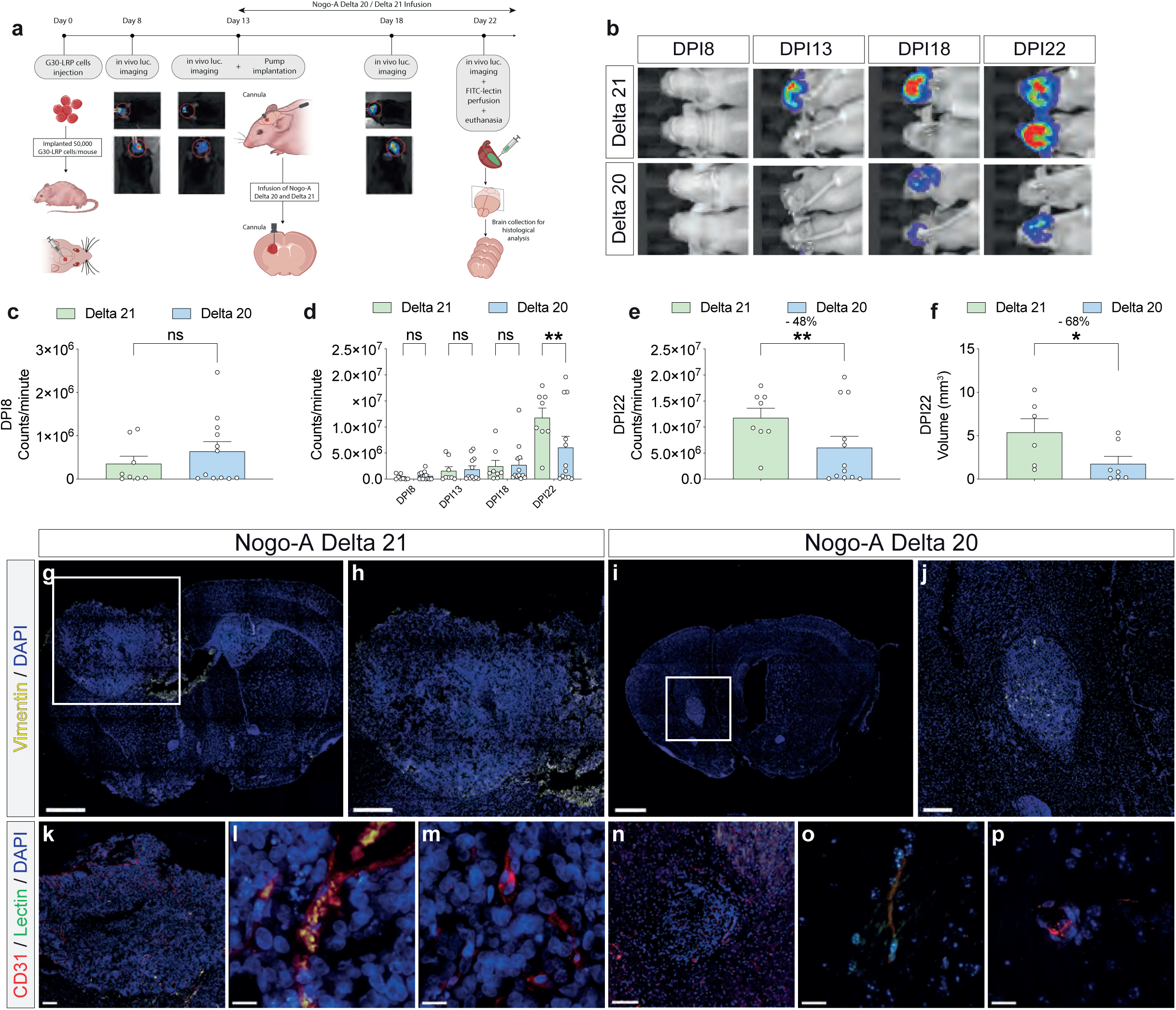
Local infusion of Nogo-A Delta 20 inhibits tumor growth in the orthotopic G30-LRP xenogeneic glioma model. (**a**) In an orthotopic G30-LRP mouse xenograft GBM model, we injected luciferase expressing G30-LRP human tumor cells, performed in vivo bioluminescence measurement in order to; monitor tumor growth. Nogo-A Delta 20 and Nogo-A Delta 21 (inactive domain of Nogo-A) were infused from days post injection (DPIs) 8 to 22 into the tumor by osmotic minipumps. Before euthanasia, mice were perfused with FITC-lectin in order to visualize blood vessel perfusion in the glial brain tumors. (**b**) Luciferase bioluminescence was measured at DPIs 8, 13 (day of randomization of mice), 18, and 22. (**b**-**d**), Intracranial astrocytoma/glioma growth is reduced in Nogo-A Delta 20 treated mice compared with Delta 21 treated control littermates (n=6-12 mice per group). (**g-p**) Coronal brain sections (40 µm) of glioblastoma-bearing mice (22 days post-inoculation, DPI22) which were perfused with FITC Lectin (green) to visualize perfused blood vessels and stained for CD31 to visualize blood vessels (red) and human Vimentin to detect tumor cells and the nuclear marker DAPI (blue). (**g**-**j**) Vimentin^+^ tumor cells allowed clear identification of the tumor mass together with increased nuclei density showed by DAPI staining. Note the clear decrease in tumor size in Nogo-A Delta 20 treated mice (**i**,**j**) as compared to Nogo-A Delta 21 control group (**g**,**h**). Moreover, Delta 21 treated mice tend to show increased tumor migration towards the other ventricle, also called butterfly glioblastoma. (**k**-**n**) Tumors exhibited high grade of vascularization, a typical feature of human glioblastoma. Data represent mean ± SEM. For statistical analysis, two-tailed unpaired Student’s t –test (**f**) and two-way ANOVA with Sidak’s multiple comparison test comparing treatment columns (**c**-**e**) were performed. \**P* < 0.05, \**P* < 0.01.

We next assessed the in vivo role of Nogo-A in tumor vascularization and investigated the effects of intratumoral Nogo-A Delta 20 on vascular morphology in human-in-mouse GBMs. The reduced tumor growth was indeed associated with a decreased tumor vascularization, as reflected by visual decrease of the vascular area (Figure 3g-p). The total, perfused and non-perfused vascular network in the mouse brain tumors was visualized using a combination of Cluster of Differentiation (CD31)^+^ immunofluorescence to visualize the total vessel network and intravascular FITC Lectin injection to label the perfused vessels^88^, thereby allowing simultaneous assessment of the total CD31^+^ vascular network including newly formed vessel sprouts that are FITC-Lectin^+^/CD31^+^ or FITC-Lectin^−^/Endomucin^+^ as well as FITC-Lectin^+^/CD31^+^ perfused functional blood vessels (Figure 3k-p). Morphologically, the vasculature of control Nogo-A Delta 21 infused tumors consisted of irregular, tortuous and highly branched blood vessels, whereas, in contrast, the vasculature of the Nogo-A Delta 20 infused tumors was more regular, less tortuous and less branched (Figure 3k-p).However, due to the very small sizes of tumors in the Nogo-A Delta 20 treated mice, a quantification of the vascular parameters of the vessel network including the vascular volume fraction, vessel length and vessel branchpoints of CD31^+^ total-, FITC-Lectin^+^/CD31^+^ perfused– and FITC-Lectin^−^/CD31^+^ non-perfused blood vessels as well as of vascular normalization was not possible.

In summary, similar to the responses observed in the syngeneic orthotopic GL-261 model, Nogo-A Delta 20 treatment inhibited tumor growth and vascularization in the xenogeneic orthotopic G30-LRP GBM model (Figure 3e,f), indicating its inhibitory effect on both mouse and human experimental glioma growth and vessels formation.

### Nogo-A Delta 20 inhibits human glioblastoma EC spreading, migration, sprouting tip cell filopodia extension

Based on the observed anti-angiogenic roles of Nogo-A in developmental and tumor angiogenesis in vivo, we next investigated the functional role of Nogo-A Delta 20 on human brain (tumor) endothelial cells in vitro. To determine whether Nogo-A Delta 20 exerts direct inhibitory effects on human CNS endothelial cells, we plated CD146+ MACS-sorted isolated/ purified, primary human brain (tumor) microvascular endothelial cells (HBMVECs and HBTMVECs) on substrates coated with the soluble Nogo-A-specific fragment Delta 20, which is known to exert potent inhibitory effects on growing neurites, spreading fibroblasts and postnatal mouse brain endothelial cells^30,59,63^. HB(T)MVEC spreading and adhesion were inhibited by Nogo-A Delta 20 (Figure 4a-j), whereas Nogo-A Delta 21 had minor inhibitory effects (Figure 4a-j), as previously reported^30,59^.

**Figure 4.**
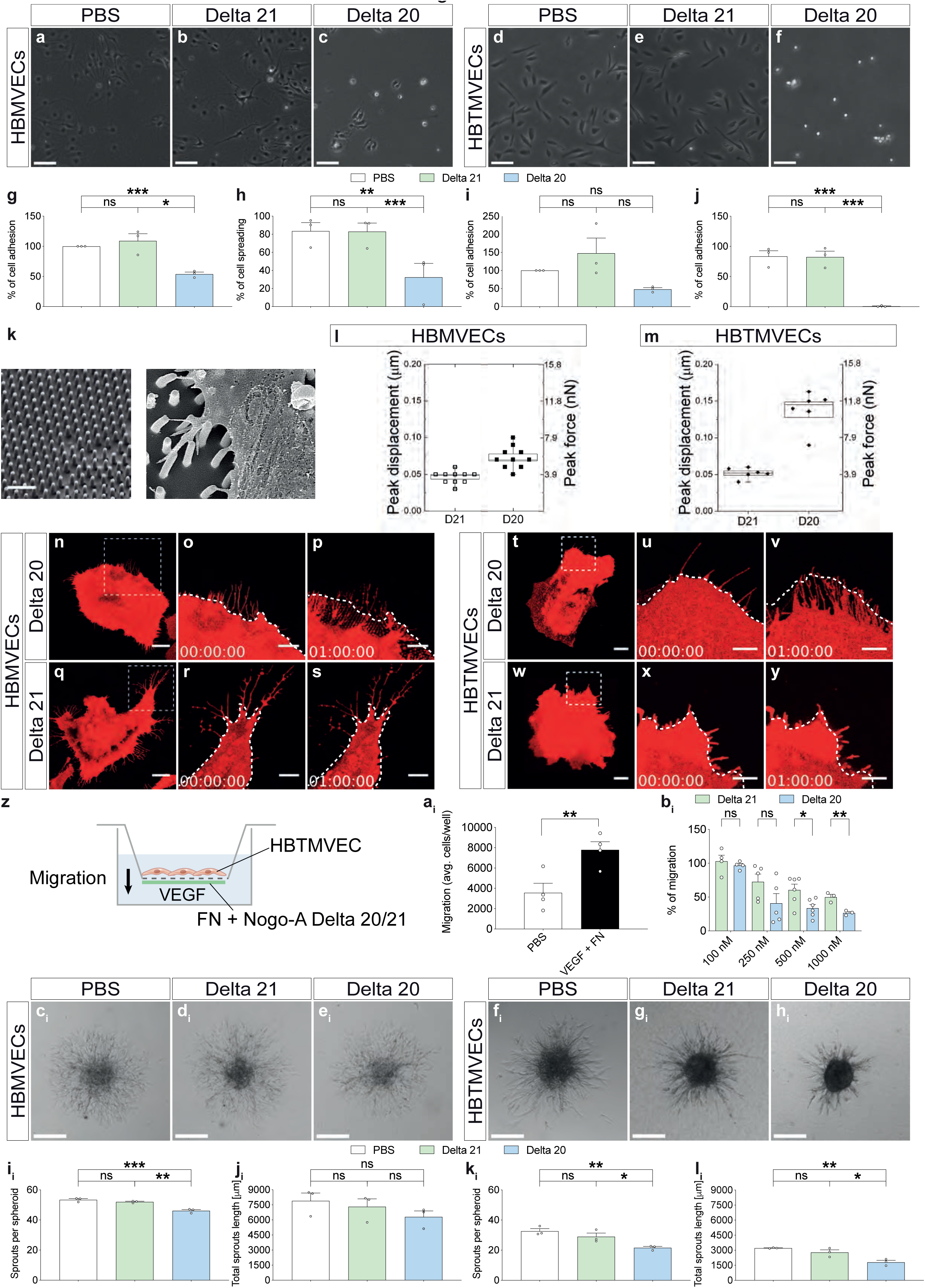
Nogo-A Delta 20 inhibits adhesion, spreading, migration and sprouting of human brain (tumor) EC. **(a**-**j)** HB(T)MVEC spreading (**g**,**i**) and adhesion (**h**,**j**) were decreased on dishes coated with Nogo-A Delta 20. No inhibition of HB(T)MVEC spreading and adhesion could be seen on dishes coated with Nogo-A Delta 21, n= 3 experiments. (**k**) Scanning electron microscopy (SEM) image of a nanopillar array (left) and EC filopodia attaching to nanopillars. Note the nanopillar-deflection caused by retracting EC filopodia allowing to optically measure the displacements of the nanopillar and the induced corresponding traction forces. (**l**,**m**) Explorative movements of HB(T)MECs (and their lamellipodia– and filopodia extensions) were reduced upon Nogo-A Delta 20 treatment as evidenced by the decreased peak nanopillar displacement and peak filopodia force (**U**) as compared to control treatment. **(n**-**y)** Time course of Nogo-A Delta 20-induced retraction of HB(T)MVEC lamellipodia followed by filopodia (**n**-**s**) over 1h. Nogo-A Delta 21-treated HB(T)MVEC showed only minor shape changes (**t**-**y**). The boxed areas in **n,q,t,w** are enlarged on their respective right side. (**z**) Scheme of the trans-well system to investigate HBTMVEC cell migration. (**a**_i_) HBTMVEC migration was significantly increased by applicating a chemotactic (VEGF-A) and a haptotactic (fibronectin, FN) gradient, n= 4 experiments. (**b**_i_) Nogo-A Delta 20 dose-dependently reduced the migration of HBTMVEC with increasing concentrations of Nogo-A Delta 20 (n=4). Note the slight inhibitory activity of Nogo-A Delta 21 **(c**_i_-**h**_i_) on HB(T)MVEC sprout formation using hanging drops composed of a collagen type I-methylcellulose matrix containing VEGF-A, basic FGF (bFGF), and EGF was decreased upon Nogo-A Delta 20 treatment (**e**_i_,**h**_i_) as compared to the PBS and Delta 21 control groups (**c**_i_,**d**_i_,**f**_i_,**g**_i_). **(i**_i_,**l**_i_) HBMVEC sprout formation (number of sprouts per spheroid) but not total sprout length were significantly reduced upon Nogo-A Delta 20 treatment as compared to the control groups (**i**_i_,**j**_i_, n=3). Nogo-A Delta 20 treatment significantly reduced the sprout formation (number of sprouts per spheroid) and total sprout length in HBTMVECs as compared to the control groups (**k**_i_,**l**_i_, n=3). Data represent mean ± SEM. For statistical analysis, one-way ANOVA with Tukey’s multiple comparison test comparing treatment columns were performed. \**P* < 0.05, \*\**P* < 0.01, \*\*\**P* < 0.001. Scale bars: 200 µm in **a**-**f** and **c**_i_-**h**_i_; 1 µm in **n**,**q**,**t**,**w**; and 5 µm in **o**,**p**,**r s**,**u**,**v**.

To mimic the in vivo situation of Nogo-A Delta 20 encountering growing brain (tumor) blood vessel endothelial tip cells we cultured DiI-labeled HB(T)MVECs on a fibronectin-coated nanopillar substrate, allowing assessment of filopodia dynamics and traction/pulling forces exerted by the spreading of HB(T)MVECs on the nanopillars (Figure 4k). HB(T)MVECs developed a flat, well-spread morphology with typical filopodia and lamellipodia (Figure 4n,q,t,w). Nogo-A Delta 20 added at a concentration of 1 µM lead to the retraction of lamellipodia after 5-10 min (Figure 4o,p,r,s and u,v,x,y), followed by a retraction of filopodia starting after 40-50 min (Figure 4o,p and u,v). Addition of soluble Nogo-A Delta 21 did not cause retraction of HB(T)MVEC lamellipodia and filopodia (Figure 4r,s and x,y)

Next, we examined the movement of Nogo-A Delta 20 treated HBMVECs and HBTMVECs on the nanopillar surface. In Nogo-A Delta 20 treated HBMVECs and HBTVECs, the mean displacements of the nanopillars were significantly increased as compared to the control group (0.07 µm and 0.05 µm and 0.145 µm and 0.052 µm, respectively, Figure 4l,m). Nogo-A Delta 20 treated HBMVEC and HBTMVEC exerted significantly increased average traction forces of 5.2 nN and 11.6 nN when compared to the Nogo-A Delta 21 treated HBMVECs and HBTMVECs (3.9 nN and 4.0 nN, respectively, Figure 5l,m). These results indicate an anti-adhesive and anti-explorative effect of Nogo-A on human brain (tumor) endothelial cells and their lamellipodial and filopodial protrusions, structures that are crucial for migration, proliferation, and sprouting of vascular ECs during brain development and in brain tumors in vivo. These data reveal that Nogo-A Delta 20 causes acute direct destabilizing effects on HB(T)MVEC lamellipodia and filopodia, structures that are crucial for pathfinding, migration and sprouting of vascular endothelial (tip) cells during developmental brain and brain tumor vascularization in vivo.

**Figure 5:**
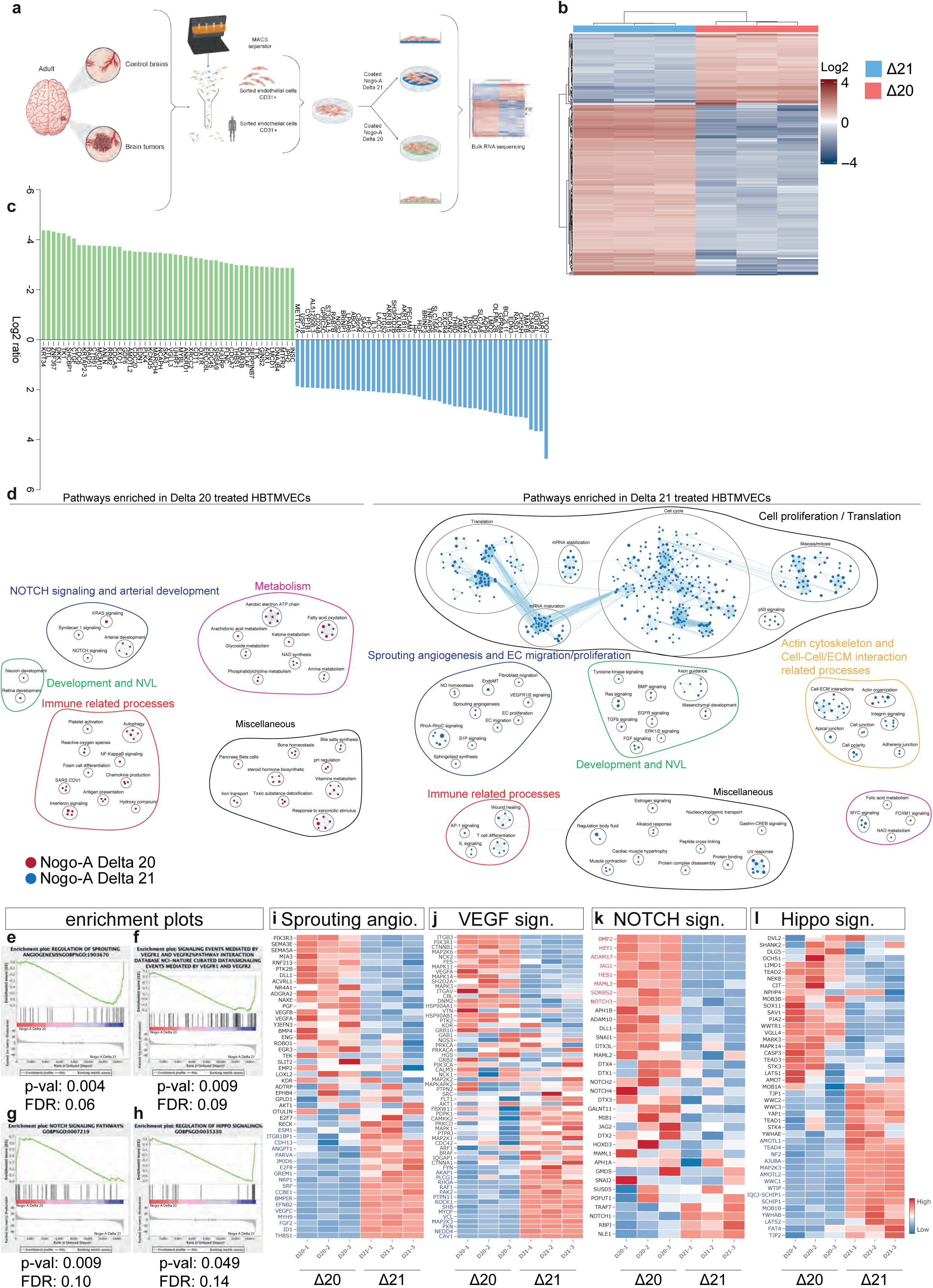
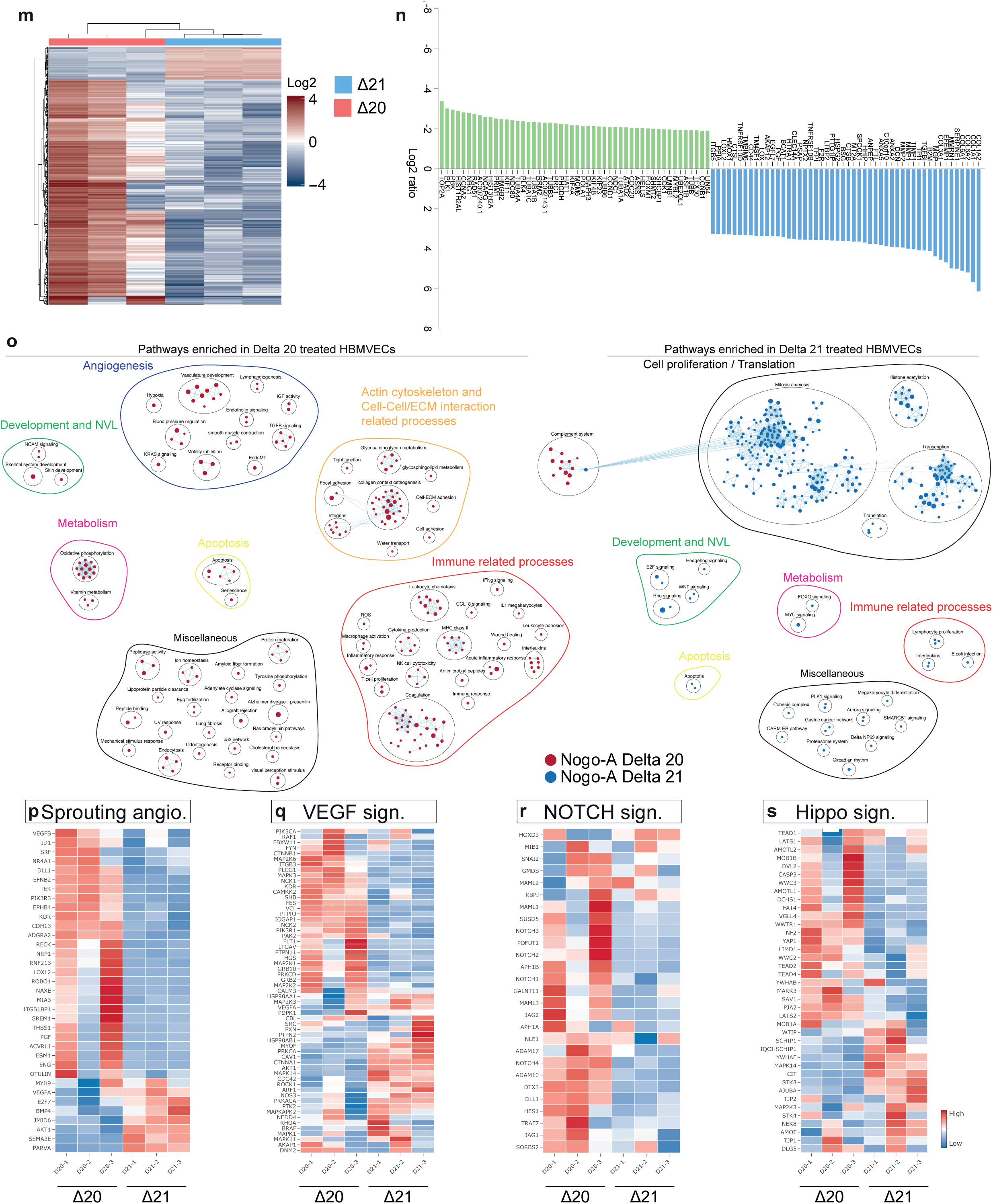
Nogo-A Delta 20 induces regulation of angiogenic pathways including Dll4-Jagged-Notch-Hey-Hes-, YAP-TAZ-CTGF-Cyr61-, and VEGF-VEGFR in HBMVECs and HBTMVECs. Transcriptome analysis via RNA sequencing of HBTMECs treated with Nogo-A Delta 20 and control Nogo-A Delta 21 peptides in three independent experiments. (**a**) Schematic representation of MACS based isolation of human adult brain (tumor) endothelial cells and subsequent treatment with coated Nogo-A Delta 20 or Nogo-A Delta 21. Cells were treated for 16hours before the RNA was extracted and processed for bulk RNA sequencing. (**b**) Heatmap and hierarchical clustering of Nogo-A Delta 20 treated HBTMVECs as compared to Nogo-A Delta 21 treated control HBTMVECs. (**c**) Top 50 significantly up (blue)-or down (green)-regulated genes detected by RNA-sequencing in HBTMVECs upon Nogo-A Delta 20 treatment as compared to Nogo-A Delta 21 treatment. Differentially regulated genes were arranged according to fold change of gene expression. Note that the Notch target gene *HEY1* was among the top up-regulated genes, while five of the top down-regulated genes were Hippo-YAP-TAZ target genes. (**d**-**h**) Gene set enrichment analysis (GSEA), cytoscape enrichment map (**d**), and enrichment plots showed a significant up-regulation of signaling pathways related to regulation of sprouting angiogenesis VEGF signaling and Hippo signaling (**e**-**f,h**) in HBTMVECs treated with Nogo-A Delta 21. Whereas pathways involved in Notch signaling (**g**) were enriched in the Nogo-A Delta 20 treatment. Pathways enriched in Nogo-A Delta 20 treated HBTMVECs are labeled in red and pathways enriched in control peptide treated HBTMVECs are labeled in blue. Pathways are indicated by colored nodes. Their size represents the number of genes they contain. Green lines indicate relationships between the pathways. Black circles group related pathways. (**i**-**l**) Heatmaps showing the expression of the genes involved in the pathways described previously namely, sprouting angiogenesis (**i**), VEGF signaling (**j**), Notch signaling (**k**), and Hippo signaling (**l**). (**m**) Heatmap and hierarchical clustering of Nogo-A Delta 20 treated HBMVECs as compared to Nogo-A Delta 21 treated control HBMVECs. (**n**) Top 50 significantly up (blue)-or down (green)-regulated genes detected by RNA-sequencing in HBMVECs upon Nogo-A Delta 20 treatment as compared to Nogo-A Delta 21 treatment. Differentially regulated genes were arranged according to fold change of gene expression. (**o**-**s**) Gene set enrichment analysis (GSEA), cytoscape enrichment map (**o**), and showed a significant up-regulation of signaling pathways related to regulation of cell proliferation, metabolism and Wnt signaling (**o**) in HBMVECs treated with Nogo-A Delta 21. Whereas pathways involved in vascular development, and cell-cell/ECM interactions (**o**) were enriched in the Nogo-A Delta 20 treatment. (**p**-**s**) Heatmaps showing the expression of the genes involved in the pathways enriched in HBTMVEC differential analysis namely, sprouting angiogenesis (**p**), VEGF signaling (**q**), Notch signaling (**r**), and Hippo signaling (**s**).

Brain tumor endothelial cell migration is a crucial process in glioma/glioblastoma vascularization^4,5,21,91^. To study the effects of Nogo-A on brain tumor vascular endothelial cell migration, we coated the underside of the inserts of transwell dishes with fibronectin mixed with different concentrations of Nogo-A Delta 20 or Nogo-A Delta 21 (Figure 4z) and 10 ng/ml recombinant VEGF-A was added in the bottom well of the chamber to stimulate endothelial cell migration^1^ (Figure 4z,ai). Nogo-A Delta 20 dose-dependently inhibited the fibronectin– and VEGF-stimulated HBTMVEC migration across the transwell insert (Figure 4c). The observed slight inhibitory effects of Nogo-A Delta 21 on HBTMVEC migration are in agreement with previous reports in neurons and postnatal mouse brain MVECs ^59^.

During brain development and in brain tumors, sprouting angiogenesis is the major process of blood vessel development ^92^. Based on our expression and functional results suggesting a restrictive role for Nogo-A in sprouting angiogenesis and ETC filopodia in developmental and tumor angiogenesis in vivo, we next investigated the functional role of Nogo-A in human angiogenic brain (tumor) endothelial cell sprouting in vitro. Therefore, we referred to an in vitro spheroid angiogenesis assay reproducing various characteristics of in vivo sprouting angiogenesis^93^. HBMVECs and HBTMVECs in the hanging drops in the sprouting spheroid assay grew vessel-like sprouts featuring multiple branches in the Nogo-A Delta 21 and PBS control groups (Figure 4ci,di,fi,gi). In contrast, Nogo-A Delta 20 significantly inhibited the number of vessel sprouts per spheroid as well as the length of the sprouts as compared to the control group in both cell types (Figure 4ei,hi,ii-li), indicating an inhibitory effect on sprouting angiogenesis.

Taken together, these results suggest that Nogo-A Delta 20 is a negative regulator of sprouting angiogenesis, endothelial spreading and migration, as well as filopodia formation in the brain and in brain tumors that can potentially be utilized to target brain tumor ECs.

### Nogo-A Delta 20 promotes NOTCH– and inhibits Hippo-YAP-TAZ and VEGFA-VEGFR2 signaling in human normal brain and brain tumor/glioblastoma ECs

To investigate the underlying mechanisms of the observed phenotypes of Nogo-A Delta 20 treatment on the brain tumor vasculature in vivo and on human brain (tumor) endothelial cells in vitro, we addressed the downstream signaling pathways induced by Nogo-A Delta 20 in human brain (tumor) endothelial cells. Therefore, we performed a time course experiment to examine the kinetics of Nogo-A-induced NOTCH-DLL4 pathway regulation, given the crucial role of this pathway as a central pattern generation for angiogenesis and EC tip-versus stalk cell discrimination^19,46,47,94,95^. In HUVECs, 1µM Nogo-A Delta 20 significantly upregulated Hey1 between 6h and 24h, peaking at 16h, as compared to controls (Supplementary Figure 6a). Thus, to address the downstream signaling pathways induced by Nogo-A Delta in human brain (tumor) ECs, we performed an unbiased transcriptome analysis by RNA sequencing of HBTMVECs and HBTMVECs treated with 1µM Nogo-A Delta 20 and 1µM Nogo-A Delta 21, respectively, for 16h (HBMVEC^Nogo-A Delta20^ and HBMVEC^Nogo-A Delta21^ as well as of HBTMVEC^Nogo-A Delta20^ and HBTMVEC^Nogo-A Delta21^, Figure 5a). RNAseq revealed 3077 and 3547 significantly regulated genes between HBTMVEC^Nogo-A Delta20^ and HBTMVEC^Nogo-A Delta21^ and between HBMVECs^Nogo-A Delta20^ and HBMVECs^Nogo-A Delta21^, respectively (Figure 5b,m). In HBTMVECs^Nogo-A Delta20^, genes involved in angiogenesis regulation, such as the Notch target gene *HEY1* figured within the top upregulated genes, whereas the Hippo-YAP-TAZ target genes *CYR61* and *CTGF* were the top downregulated genes (Figure 5c). Among the top-regulated genes in HBMVECs^Nogo-A Delta20^ figured genes implicated in endothelial cell proliferation and ECM-cell related processes, such as *COL1A2*, *COL1A1*, *COL6A2*, whereas *TOP2A* was the top downregulated gene (Figure 5n).

Next, to further address the molecular pathways regulated by Nogo-a Delta 20 in ECs, we performed gene set enrichment analyses (GSEA)^96^ between Nogo-A Delta 20 treated HBTMVECs/HBMVECs and the control Nogo-A Delta 21 treated HBTMECs/HBMVECs, respectively. In HBTMVEC^Nogo-A Delta20^, GSEA followed by pathway visualization using cytoscape^96^ revealed that upregulated genes were mainly involved in negative regulation of(sprouting) angiogenesis, while the downregulated genes were linked to positive regulation of (sprouting) angiogenesis (Figure 5d). Indeed, NOTCH signaling was significantly enriched upon Nogo-A Delta 20 treatment in HBTMVECs whereas VEGF-VEGFR signaling, Hippo-YAP-TAZ signaling and sprouting angiogenesis showed significant enrichment in Nogo-A Delta 21 HBTMVECs (Figure 5d-h).

We next examined genes driving the enrichment of positive and negative regulation of sprouting angiogenesis in brain tumor ECs. Genes involved in the positive regulation of sprouting angiogenesis including *ESM-1* (a tip cell marker) as well as the pro-angiogenic NVL regulatory genes *EFNB2*, *NRP1*, *BMPER* and the pro-angiogenic cue *FGF2*^97^ were significantly downregulated in HBTMVEC^Nogo-A Delta20^ (Figure 5i). Similarly, genes involved in the regulation of the key pro-angiogenic *VEGFA-VEGFR2* pathway such as *MAP2K3*, *CAV1*, *PXN*, and *VCL*, as well as genes involved in the pro-angiogenic Hippo-YAP-TAZ pathways including *LATS2*, *AJUBA*, *NF2* and *TEAD4* were significantly downregulated in HBTMVEC^Nogo-A Delta20^ (Figure 5j,l), indicating that Nogo-A negatively regulates those genes promoting angiogenic sprouting in brain tumor ECs. On the other hand, genes involved in the regulation of the key anti-angiogenic *JAG1*-*DLL4-NOTCH* pathway such as *HEY1*, *HES1*, *NOTCH3*, *JAG1*, and *ADAM17* were significantly upregulated in HBTMVEC^Nogo-A Delta20^ (Figure 5k), indicating that Nogo-A positively regulates these genes inhibiting angiogenic sprouting in brain tumor ECs.

Notably, we observed similar yet less strong regulatory effects of Nogo-A Delta 20 in normal brain ECs (HBMVECs) including overlapping genes for sprouting angiogenesis and *VEGFR-VEGFR2* (*CAV1 and PXN*, *Hippo-YAP-TAZ* (*AJUBA*) and *JAG1-DLL4-NOTCH* (*NOTCH3* and *JAG1*) in as compared to the brain tumor ECS (HBTMVECs) (Figure 5p-s), suggesting that Nogo-A Delta 20 exerts stronger regulatory effects on underlying mechanisms driving vascular growth and sprouting angiogenesis in the more active brain tumor ECs.

In HBMVEC^Nogo-A Delta20^, on the other hand, GSEA analysis revealed regulation of pathways involved in angiogenesis, cell-cell and cell-ECM related processes and cell proliferation (Figure 5o). Based on the observed regulatory effects of Nogo-A Delta 20 on sprouting angiogenesis, we next analyzed the expression of main regulators of the key anti– and pro-angiogenic pathways Dll4-Jagged-Notch, Hippo-YAP-TAZ and VEGF/VEGFR (Figure 6a-p, Supplementary Figure 6b,c). Nogo-A Delta 20 significantly upregulated the Notch downstream effectors *HES1* and *HEY1*, in HBTMVEC^Nogo-A Delta20^ but not in HBMVEC^Nogo-A Delta20^ (Figure 6a,e). Validation by qRT-PCR and Western blot confirmed Nogo-A Delta 20-induced upregulation of *HEY1* and *HES1* (but not of NOTCH1) at the mRNA level in HBMVECs and in HBTMVECs (Figure 6b,f), and Nogo-A Delta 20 induced upregulation of the Notch Intracellular Domain (NICD) in HBMVECs and to a lesser extent in HBTMVECs (Figure 6c,d,g,h), indicating potentially different underlying mechanisms between normal brain and brain tumor ECs. On the other hand, Nogo-A Delta 20 significantly downregulated the Hippo-YAP-TAZ pathway downstream effectors *CTGF* and *CYR61* in HBTMVEC^Nogo-A Delta20^ and *CYR61* but not *CTGF* in HBMVEC^Nogo-A Delta20^ (Figure 6i,m). Validation by qRT-PCR and Western blot confirmed Nogo-A Delta 20-induced downregulation of *CTGF* and *CYR61* at the mRNA level in HBMVECs and in HBTMVECs (Figure 6j,n), and Nogo-A Delta 20 induced downregulation of YAP-TAZ signaling via upregulation of the ratios of the phosphorylated forms of LATS1 (p-LATS1) and of YAP (p-YAP) over the total proteins, namely p-LATS1/LATS1 and p-YAP/YAP (p-YAP being degraded resulting in decreased YAP-TAZ signaling) in HBMVECs and HBTMVECs (Figure 6k,l,o,p), indicating Nogo-A Delta 20 negative regulatory effects on both normal brain and brain tumor ECs.

**Figure 7:**
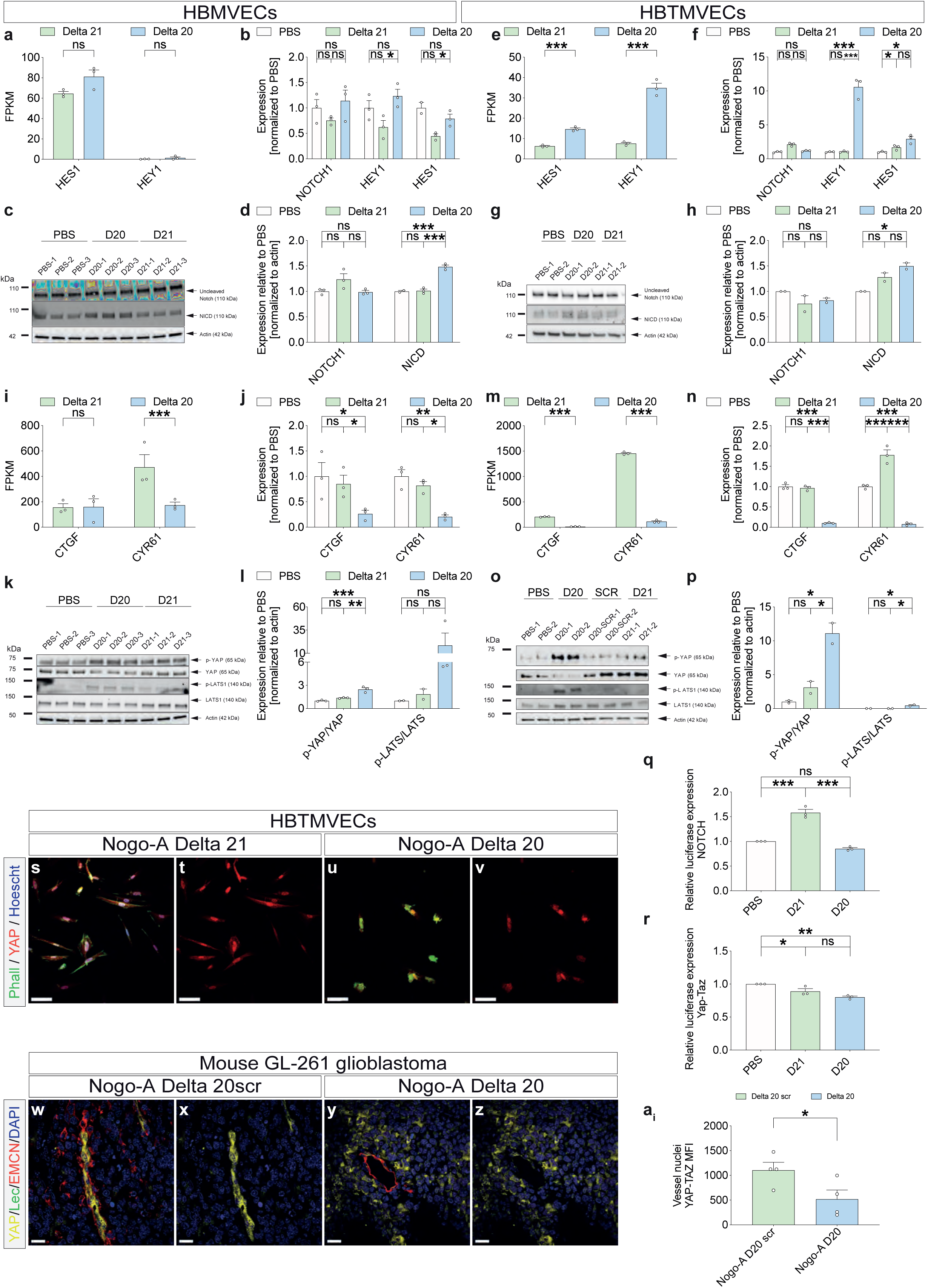
Nogo-A Delta 20 restricts HBMVEC and HBTMVEC sprouting angiogenesis by regulating the DLL4-NOTCH, the YAP-TAZ and the VEGF-VEGFR pathways. (**a**-**h**) Nogo-A Delta 20 treatment induced upregulation of Notch signaling in HB(T)MVECs. (**a**)RNAseq analysis showed that the NOTCH pathway downstream effectors *HES1* and *HEY1* gene expression (FPKM) was upregulated by Nogo-A Delta 20 treatment in HBMVECs (**a**) and significantly upregulated in HBTMVECs (**e**). qRT-PCR analysis for DLL4-NOTCH pathway genes performed on HBMVECs (**b**) and HBTMVECs (**f**) 16hrs post-stimulation with Nogo-A Delta 20, and Nogo-A Delta 21 showed that Nogo-A Delta 20 induced a significant upregulation of the Dll4-Jagged-Notch signaling pathway including: NOTCH1, HEY1 and HES1 (**b**,**f**). (**q-r**). Western blots using antibodies against uncleaved Notch1, Notch intracellular domain (NICD), on HBMVECs (**c**,**d**) and HBTMVECs (**g**,**h**) treated with Nogo-A Delta 20, Nogo-A, and Nogo-A Delta 21. Nogo-A Delta 20 increased NICD protein levels and without affecting NOTCH1 protein levels (**d**) in HBMVECs but Nogo-A Delta 20 tend to decrease uncleaved NOTCH1 (**h**) and increase NICD protein levels (**h**). (**i**-**p**) Nogo-A Delta 20 treatment induced downregulation of Hippo-YAP/TAZ signaling in HB(T)MVECs. RNAseq analysis showed that YAP/TAZ pathway downstream effector *CTGF* gene expression (FPKM) was significantly downregulated by Nogo-A Delta 20 treatment in HBMVECs but not *CYR61* (**q**) and that both gene expression was significantly downregulated in HBTMVECs (**m**). (**j-n**) qRT-PCR analysis for YAP-TAZ pathway genes performed on HBMVECs (**j**) and HBTMVECs (**n**) 16hrs post-stimulation with Nogo-A Delta 20 and Nogo-A Delta 21 showed that Nogo-A Delta 20 induced a significant downregulation of the YAP-TAZ target genes in HBMVECs and HBTMVECs, namely CTGF and CYR61. (**k**,**l**,**o**,**p**) Western blots using antibodies against YAP, phosphorylated YAP, LATS1 and phosphorylated LATS1 on HBMVEC (**k**,**l**) and HBTMVECs (**o**,**p**) treated with Nogo-A Delta 20 and Nogo-A Delta 21. In HBMVECs and HBTMVECs Nogo-A Delta 20 decreased YAP protein levels and increased phosphorylated YAP and phosphorylated LATS1 protein levels, leading to increased p-YAP/YAP and p-LATS1/LATS1 ratios (**k**,**l**,**o**,**p**). (**q**,**r**) Dual luciferase assay using reporter plasmids for Notch (**q**) and YAP-TAZ (**r**) signaling pathways. **(s**-**v**) HBTMVECs were stained for actin (Phalloidin, green), YAP (red), and the general nuclear marker Hoechst (blue). (**w-a**_i_). Coronal brain sections (40 µm) of GL-261 glioblastoma-bearing mice (28 days post-inoculation, DPI28) which were perfused with FITC Lectin (green) to visualize perfused blood vessels and stained for endomucin to visualize blood vessels including endothelial tip cells (red), YAP (yellow) and the nuclear marker DAPI (blue). YAP mean fluorescence intensity in endothelial cell DAPI was significantly reduced upon treatment with Nogo-A Delta 20 as compared to control peptide (**a**_i_, n= 4 mice per group). Data represent mean ± SEM. For statistical analysis, Wald test corrected for multiple testing using the Benjamini and Hochberg method (**a**,**e**), two-way ANOVA with Sidak’s post-hoc test (**b**,**d**,**f**,**h**,**j**,**l**,**n**,**p**), one-way ANOVA with Tukey’s post-hoc test (**q**,**r**)and two-tailed unpaired Student’s t –test (**a_i_**) were performed. \**P* < 0.05, \*\**P* < 0.01, \*\*\**P* < 0.001. Scale bars: 150 μ in **s**-**v**; and 40 µm in **w**-**z**.

Next, we further validated the alteration of YAP expression at the protein level using immunofluorescence in HBTMVECs in vitro and in the GL-261 mouse model in vivo. We observed a decrease of the expression of YAP in EC nuclei (and thus Hippo-YAP-TAZ signaling, Figure 6s-ai), further confirming a negative regulatory effect of Nogo-A Delta 20 on YAP-TAZ signaling in brain tumor ECs in vitro and in vivo. At the functional level, whereas dual-luciferase reporter assays confirmed YAP downregulation upon Nogo-A Delta 20 treatment, and surprisingly, NOTCH did not reveal significant regulation upon Nogo-A Delta 20 treatment (Figure 6q,r).

Finally, given its central role in angiogenesis and vascular growth and based on its regulation in the bulkRNAseq data, we validated the alteration of transcriptional expression of *VEGFR2* at the protein level using Western blot and immunofluorescence. We observed Nogo-A Delta 20 induced downregulation of the phosphorylated form of VEGFR2 (p-VEGFR2) in HUVEC ^Nogo-A Delta20^ (Supplementary Figure 6b,c), indicating a negative regulatory effect of Nogo-A Delta 20 on VEGFR2 and suggesting a potential interaction between the Nogo-A and VEGF-A-VEGFR2 pathways in ECs.

Taken together, these results suggest that Nogo-A Delta 20 exerts restricting regulatory effects on brain (tumor) sprouting angiogenesis via interaction with the JAG1-DLL4-NOTCH, the Hippo-YAP-TAZ and the VEGFA-VEGFR2 signaling pathways in vivo and in vitro.

### Metabolomics reveals negative regulation of brain (tumor) endothelial glucose metabolism upon Nogo-A Delta 20 treatment

In addition to be regulated by VEGF-VEGFR, Dll4-Jagged-Notch, and Hippo-YAP-TAZ signaling, sprouting angiogenesis, ETC formation, and endothelial lamellipodia– and filopodia-dynamics crucially depend on vascular endothelial metabolism during development and in disease^95,98,99^. Furthermore, endothelial cell glycolysis regulates EC rearrangement by promoting filopodia formation and by decreasing intercellular adhesion^99^. To address possible effects of Nogo-A Delta 20 on endothelial metabolism, we performed unbiased metabolic profiling using liquid chromatography tandem mass spectrometry (LC-MS/MS) ^100^ in Nogo-A Delta 20 (1µM) treated HBMVECs and HBTMVECs compared to Nogo-A Delta 21 (1µM) treated HBMVECs and HBTMVECs, respectively (Figure 7a-l). Hierarchical clustering revealed that metabolite levels of Nogo-A Delta 20 treated HB(T)MVECs clearly separated from Nogo-A Delta 21 treated HB(T)MVEC, revealing significantly regulated metabolites between the groups (Figure 7a-d,g-j).

**Figure 7:**
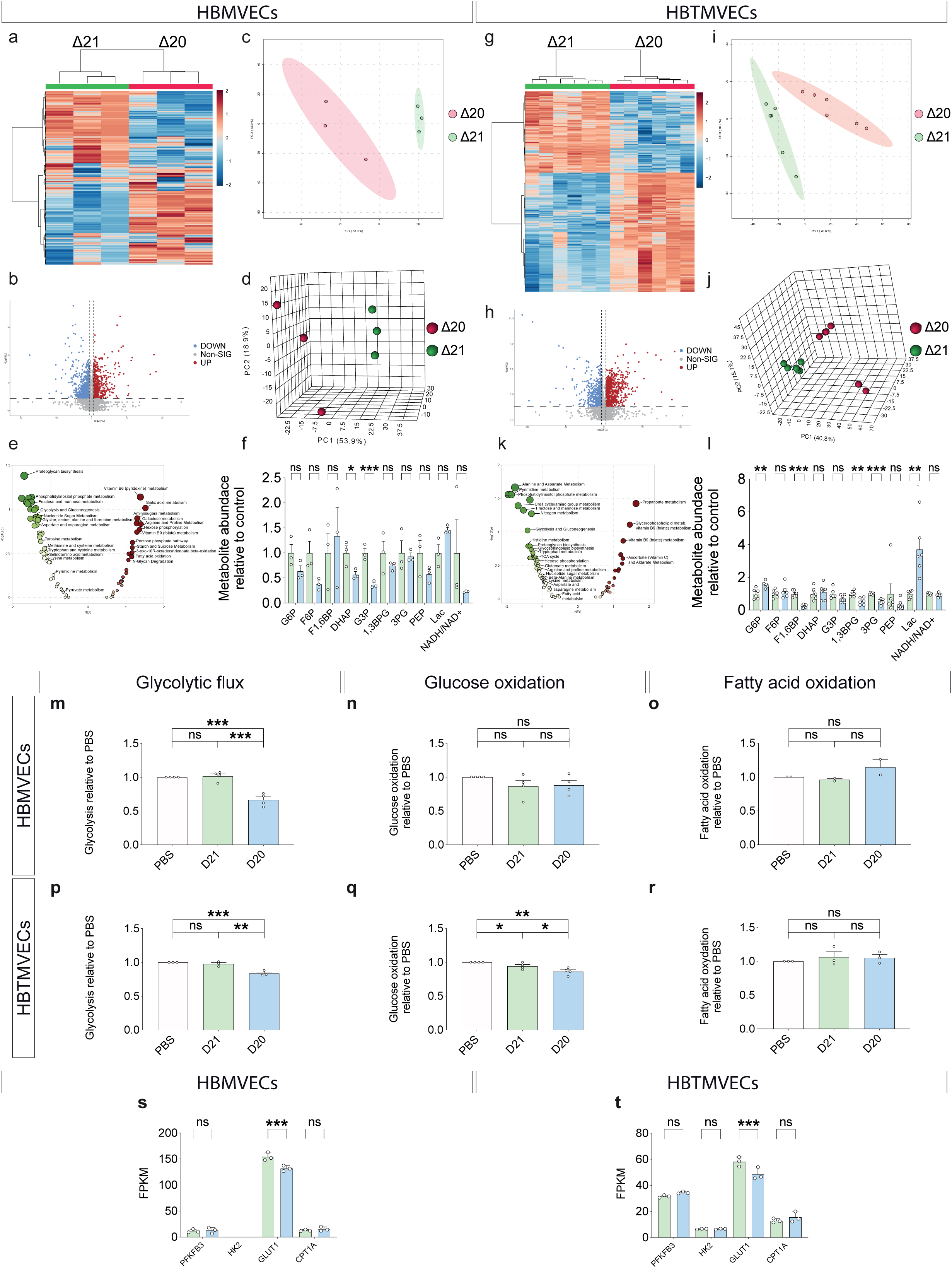
Nogo-A Delta 20 negatively regulates endothelial glucose metabolism but does not affect fatty acid oxidation in human brain (tumor) endothelial cells. (**a**-**k**) Metabolite profile comparison between Nogo-A Delta 20 treated HBMVECs/HBTMVECs and Nogo-A Delta 21 control treated HBMVECs/HBTMVECs (n=3-5). Heatmap and hierarchical clustering showing all detected differentially regulated metabolites in HBMVECs (**a**) and HBTMVECs (**f**) upon Nogo-A Delta 20 treatment. Scatter plot showing the significantly regulated metabolites upon Nogo-A treatment in HBMVECs (**b**) and HBTMVECs (**g**). 678 and 755 metabolites were significantly differentially in HBMVECs^Nogo-A Delta20^ and HBTMVECs^Nogo-A Delta20^ respectively (**b**,**g**, indicated in red) while 645 and 588 metabolites were significantly downregulated HBMVECs^Nogo-A Delta20^ and HBTMVECs^Nogo-A Delta20^ respectively (**b**,**h**, indicated in blue). Two-dimensional and three-dimensional PCA plot of log-transformed normalized concentration of all detected metabolites, each dot represents a sample, and is colored by experimental group (**c**,**d**,**i**,**j**). Gene set enrichment analysis (GSEA) indicated a significant regulation of various metabolic pathway in HBMVECs (**f**) and HBTMVECs (**l**) upon Nogo-A Delta 20 treatment. Relative abundance of glycolysis intermediates in HBMVECs (**F**) and HBTMVECs (**F**) upon Nogo-A Delta 20 treatment. **(m**-**r)** Metabolic assays in HBMVECs (**m**-**o**) and HBTMVECs (**p**-**r**), upon Nogo-A Delta 20 treatment. Nogo-A Delta 20 treatment decreased the glycolytic flux in both cell types (**m**,**p**, n=3), glucose oxidation in HBTMVECs but not in HBMVECs (**n**,**q**, n=3) as compared to the tested controls. Nogo-A Delta 20 treatment didn’t affect fatty acid oxidation in both cell types (**o**,**r**, n=2-3). (**s**-**t**) Key metabolic genes expression (FPKM) in HB(T)MVECs upon Nogo-A Delta 20 and Nogo-A Delta 21 control treatment. BulkRNAseq revealed a significant downregulation of glucose transporter GLUT1 (*SLC2A1*) in both cell types upon Nogo-A Delta 20 treatment as compared to control treatment. Key glycolytic enzymes *PFKFB3* and *HK2* gene expression as well as fatty oxidation enzyme *CPT1A* expression were not changed upon Nogo-A Delta 20 treatment. Data represent mean ± SEM. For statistical analysis, two-tailed unpaired Student’s t –test (**f**,**l**), one-way ANOVA with Tukey’s post-hoc test (**m**-**k**) and Wald test corrected for multiple testing using the Benjamini and Hochberg method (**s**,**t**) were performed. \**P* < 0.05, \*\**P* < 0.01, \*\*\**P* < 0.001.

This analysis showed 1323 and 1433 metabolites altered by the Nogo-A Delta 20 treatment in HBMVECs and HBTMVECs, respectively (Figure 7b,h). Principal component analysis (PCA) further identified specific groups of metabolites including those involved in endothelial glucose metabolism to be different in Nogo-A Delta 20 treated HBMVECs and HBTMVECs as compared to the control groups (Figure 7c,d, Supplementary Figure 7c,d).

Next, we examined the metabolic pathways regulated by Nogo-A Delta 20 in brain (tumor) ECs. GSEA^96^ revealed glycolysis to be downregulated in HBMVECs and HBTMVECs upon Nogo-A Delta 20 treatment and fatty acid metabolism pathways to be downregulated in HBTMVEC^Nogo-A Delta20^ while being upregulated in HBMVEC^Nogo-A Delta20^ (Figure 7e,k).

We next analyzed the abundance of metabolites involved in glycolysis (Figure 7f,l). Interestingly, we found that among the eight glucose metabolites detected in HBMVECs and HBTMVECs, Dihydroxyacetone Phosphate (DHAP) and Glyceraldehyde 3-phosphate (G3P) were significantly decreased in Nogo-A Delta 20 treated HBMVECs, whereas Fructose 1,6-bisphosphate (F1,6BP), 1,3-Bisphosphoglycerate (1,3BPG), 3-Phosphoglycerate (3PG) were significantly decreased in HBTMVECs (Figure 7f,l, Supplementary Figure 7f). Surprisingly, upon Nogo-A Delta 20 treatment, lactate levels were increased by 35% in HBMVECs and by 55% in HBTMVECs respectively (Figure 7f,l). Moreover, the NADH/NAD+ ratio indicating for metabolic activity^101,102^ showed a decrease (although not significant) upon Nogo-A Delta 20 in HBMVECs and no change in HBTMVECs (Figure 7f, Supplementary Figure 7f). Taken together, these results suggest that Nogo-A Delta 20 exerts its negative regulatory effects on CNS sprouting angiogenesis via negative regulation of brain (tumor) endothelial (glucose) metabolism.

### Nogo-A regulates brain (tumor) endothelial glucose-, but not fatty acid metabolism

To further examine the observed effects of Nogo-A Delta 20 on brain (tumor) endothelial metabolism, we next performed functional metabolic assays addressing endothelial glucose and fatty acid metabolism in HBTMVECs and HBMVECs. Referring to a glycolytic flux assay^103^, we observed a significant reduction of glycolysis upon Nogo-A Delta 20 treatment as compared to the control conditions in both HBTMVECs and HBMVECs (Figure 7m,p). Moreover, HBTMVEC and HBMVEC showed different response to Nogo-A Delta 20 treatment on glucose oxidation. Indeed, HBTMVECs glucose oxidation was reduced by Nogo-A Delta 20 but remained unregulated in HBMVECs (Figure 7n,q). These data indicate a negative regulatory effect of Nogo-A on HBTMVEC and HBMVEC glucose metabolism and highlight potential differences between brain tumor and healthy brain endothelial cells.

Next, we assessed whether Nogo-A Delta 20 also regulates endothelial fatty acid oxidation (FAO) known to support endothelial stalk cell proliferation^99,104^, and to protect quiescent ECs against oxidative stress^105^. HBMVECs and HBTMVECs fatty acid oxidation (Figure 7o,r) was not regulated by Nogo-A Delta 20. Expression analysis of key regulators of glucose metabolism, namely Phosphofructokinase 3 (*PFKFB3*), Hexokinase 2 (*HK2*) and Glucose transporter 1 (GLUT1) revealed that the Nogo-A Delta 20 might exerts its inhibitory effects via downregulation of GLUT1, but not PFKFB3 or HK2 (Figure 7s,t). Consistent with the absence of regulation of fatty acid oxidation, the expression of key regulator of fatty acid metabolism, Carnitine palmitoyltransferase 1A (CPT1a)^98,99^ was not regulated upon Nogo-A Delta 20 treatment.

Taken together, these data reveal that Nogo-A Delta 20 decreases brain (tumor) endothelial glucose metabolism without affecting brain (tumor) endothelial fatty acid metabolism, indicating a negative regulatory role of Nogo-A on sprouting angiogenesis via inhibition of endothelial glucose metabolism.

### The S1PR2/TSPAN3/SDC4 receptor complex mediates Nogo-A Delta 20 induced inhibitory effects on human brain tumor/glioblastoma ECs

The Nogo-A Delta 20 domain signals via the S1PR2 / TSPAN3 / SDC3/4 receptor complex^61–64,106,107^ but is expression pattern on human brain vascular endothelial cells across development and disease remains unknown.

Therefore, we addressed the gene expression of Nogo-A (RTN4 gene isoform) and its receptor complex S1PR2/TSPAN3/SDC4 in our recently constructed single-cell atlas of the human brain vasculature in fetal brain development, in the adult control brain and in brain tumors including low-grade glioma (LGG) and high-grade gliomas/glioblastoma (GBM)^28^ (Figure 8a-j). Interestingly, whereas S1PR2 and SDC4 showed only very low expression levels in both ECs and PVCs of the NVU, TSPAN3 showed high expression in ECs and PVCs across the fetal, adult and diseased brain entities (Figure 8a-j). In ECs, however, TSPAN3 showed expression in the fetal brain, downregulation in the adult brain, a slight elevation in the LGGs and a more pronounced expression increase in GBMs (Figure 8j), indicating its role in developmental and pathological (tumor) vascular growth, typical for onco-fetal proteins^41,108^. Notably, because the RTN4 genes encodes for all three Nogo protein isoforms (Nogo-A, Nogo-B, Nogo-C)^60^ and because Nogo-B is known to be highly expressed in the (brain) vasculature^109,110^ its mRNA expression pattern does not allow for conclusive assessment of Nogo-A expression in the human brain vasculature (Figure 8a-j). In HBMVECs, we found low expression of S1PR2, and SDC4, while TSPAN3 was the most expressed Nogo-A receptor of the complex. Moreover, we observed a similar expression pattern in HBTMVECs (TSPAN3>SDC4>S1PR2) (Figure 8k)

**Figure 8:**
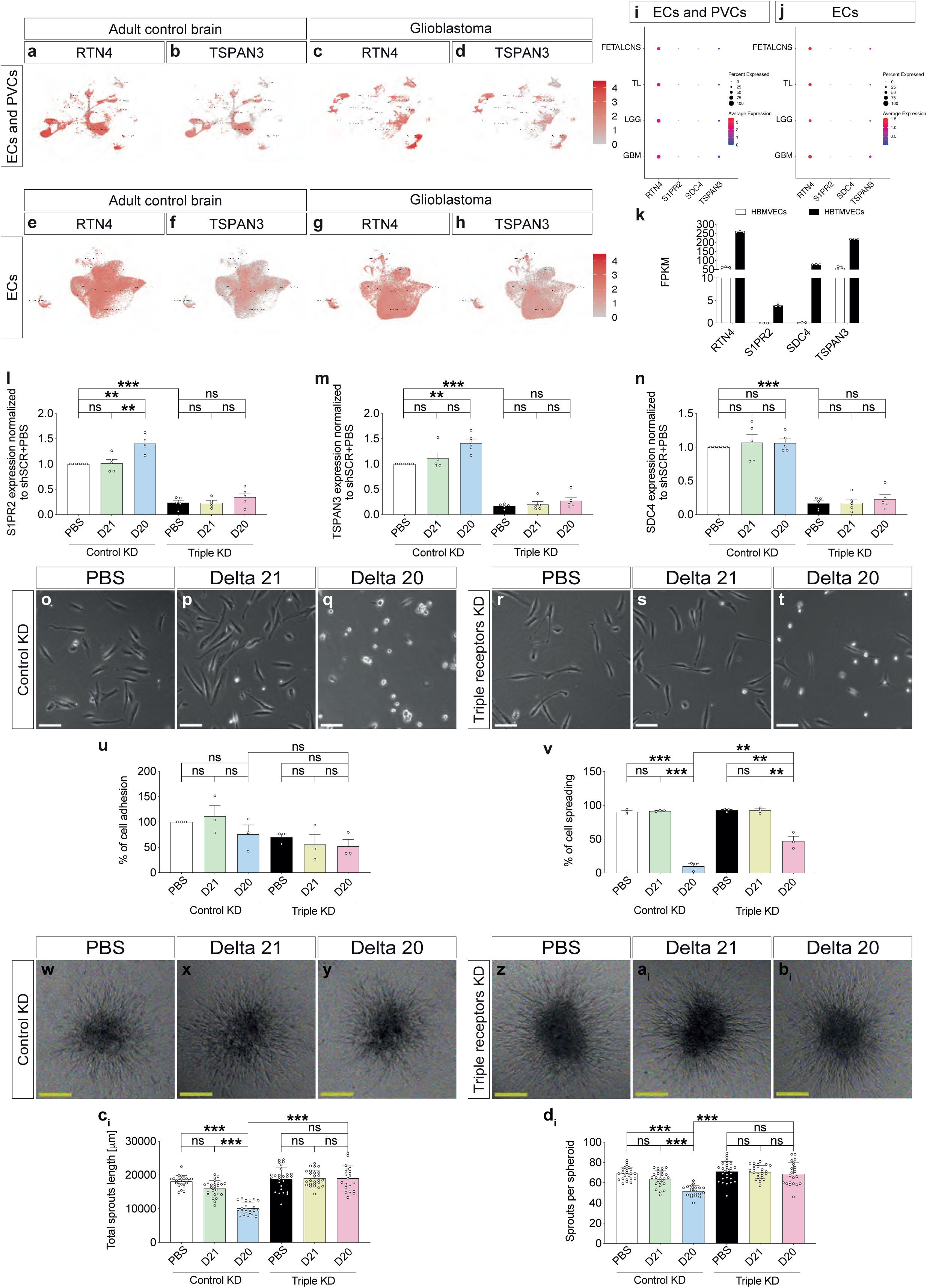
Knock-down of Nogo-A receptor complex S1PR2/TSPAN3/SDC4 reverse Nogo-A Delta 20 inhibitory effects on brain tumor endothelial cell (HBTMVEC) spreading and sprouting. (**a**-**h**), UMAPs plots, color-coded for expression of Nogo (*RTN4*) and Nogo-A receptor Tetraspanin 3 (*TSPAN3*) in perivascular cells and endothelial cells of adult normal brain (**a**,**b**) and glioblastoma (**c**,**d**) as well as in CD31+/CD45-FACS sorted endothelial cells of adult normal brain (**e**,**f**) and glioblastoma (**g**,**h**). (**i**,**j**) Dotplots of Nogo and Nogo-A receptors eCs and PVCs (**i**) and sorted eCs (**j**) from fetal brain, adult control brain, low-grade glioma and glioblastoma. (**k**) bulkRNAseq analysis showed that the Nogo and Nogo-A receptor genes were highly expressed in HBTMVECs, while the expression of Nogo-A receptors S1PR2 and SDC4 were low in HBMVECs. (**l**-**n**) qRT-PCR analysis for Nogo-A receptors *S1PR2* (**l**), *TSPAN3* (**m**) and *SDC4* (**n**)genes performed on HBTMVECs treated with short hairpin RNA (shRNA) targeting Nogo-A receptor complex (S1PR2, TSPAN3, SDC4, HBTMVEC^Triple KD^) or control shRNA (HBTMVEC^Control KD^) and stimulated for 16hrs with Nogo-A Delta 20, and Nogo-A Delta 21 confirmed Nogo-A receptors knock-down. Note that Delta 20 induced a significant upregulation of S1PR2 and TSPAN3 but not SDC4 in HBTMVEC^Control KD^. **(o**-**v)** HBTMVEC spreading (**v**) and adhesion (**u**) were decreased on dishes coated with Nogo-A Delta 20 both in HBTMVEC^Triple KD^ and HBTMVEC^Control KD^. No inhibition of HBTMVEC spreading and adhesion could be seen on dishes coated with Nogo-A Delta 21. Triple receptor KD didn’t affect the adhesion of HBTMVECs but partially rescued the spreading of HBTMVECs as indicated by significant increase in HBTMVEC^Triple KD^ cell spreading on Nogo-A Delta 20 coated dishes as compared to HBTMVEC^Control KD^ (n= 3). **(w**-**d_i_)** HBTMVEC^Control KD^ sprout formation using hanging drops composed of a collagen type I-methylcellulose matrix containing VEGF-A, basic FGF (bFGF), and EGF was decreased upon Nogo-A Delta 20 treatment (**e_i_**,**h_i_**) as compared to the PBS and Delta 21 control groups (**w**-**y**,**c_i_**,**d_i_**). HBTMVEC^Triple^ KD showed no difference in sprout number and sprout length between Nogo-A Delta 20 and control treatments (**z**-**b_i_**,**c_i_**,**d_i_,** n=1). (Data represent mean ± SEM. For statistical analysis, two-tailed unpaired Student’s t –test (**l**-**n**) and two-way ANOVA with Sidak’s multiple comparison test comparing treatment columns (**u**,**v**,**c_i_**,**d_i_**) were performed. \**P* < 0.05, \*\**P* < 0.01, \*\*\**P* < 0.001. Scale bars: 200 µm in **o**-**t** and **w**-**b_i_**.

Based on these expression studies suggesting a role for the Nogo-A receptor complex in the human brain tumor vasculature in vivo, we next investigated the functional role of S1PR2/TSPAN3/SDC4 in human brain tumor endothelial cells in vitro. To that regard, we used short hairpin RNA (shRNA) to knock down the Nogo-A Delta 20 receptor components in HBTMVECs (Figure 8l-n). In shRNA-mediated S1PR2/TSPAN3/SDC4 triple knock-down (TKD) HBTMVECs (HBTMVEC^TKD^), expression of S1PR2, TSPAN3, and SDC4 was significantly decreased at the mRNA level as revealed by qRT-PCR (Figure 8l-n).

To address the effects of the Nogo-A triple receptor complex knockdown at the functional level, we performed spreading and spheroid assays in HBTMVEC^TKD^ treated with Nogo-A Delta 20 or the control peptide Nogo-A Delta 21 (HBTMVEC^TKD – Nogo-A Delta20^ and HBTMVEC^TKD – Nogo-A Delta21^). The Nogo-A Delta 20 induced inhibitory effects on HBTMVEC cell spreading (and to a lesser extent on cell adhesion) were rescued in HBTMVEC^TKD^ (Figure 8o-v). Similarly, the Nogo-A Delta 20-induced inhibitory effects on HBTMVEC cell sprouting were reversed in HBTMVEC^TKD^ (Figure 8w-d_i_), indicating that the S1PR2/TSPAN3/SDC4 receptor complex at least partially mediates the Nogo-A Delta 20 induced inhibitory effects on HBTMVEC in vitro.

Next, to address whether the triple knockdown of the Nogo-A receptor complex components affects the downstream signaling pathways induced by Nogo-A Delta 20 in human brain (tumor) ECs, we performed an unbiased transcriptome analysis by RNA sequencing of HBTMVECs^Triple KD^ treated with 1µM Nogo-A Delta 20 and 1µM Nogo-A Delta 21, respectively, for 16h (HBTMVEC^Triple KD – Nogo-A Delta20^ and HBTMVEC^Triple KD – Nogo-A Delta21^ _)_ (Figure 9a-k). RNAseq revealed 4025 significantly differentially regulated genes between HBTMVEC^Triple KD – Nogo-A Delta20^ and HBTVMEC^Triple KD – Nogo-A Delta21^ (Figure 9a). In HBTMVECs^Triple KD – Nogo-A Delta20^, genes involved in angiogenesis, as the fatty acid transporter Fatty acid binding protein 4 (*FABP4)* figured within the top upregulated genes, whereas the Notch pathway component gene ADAM Metallopeptidase with Thrombospondin Type 1 Motif 4 (*ADAMTS4)* as well as the pro-angiogenic genes *HBF* and *COL14A1* were among the top downregulated genes (Figure 9b) in agreement with the observed roles of Nogo-A in endothelial metabolism, sprouting angiogenesis, and Notch signaling.

**Figure 9:**
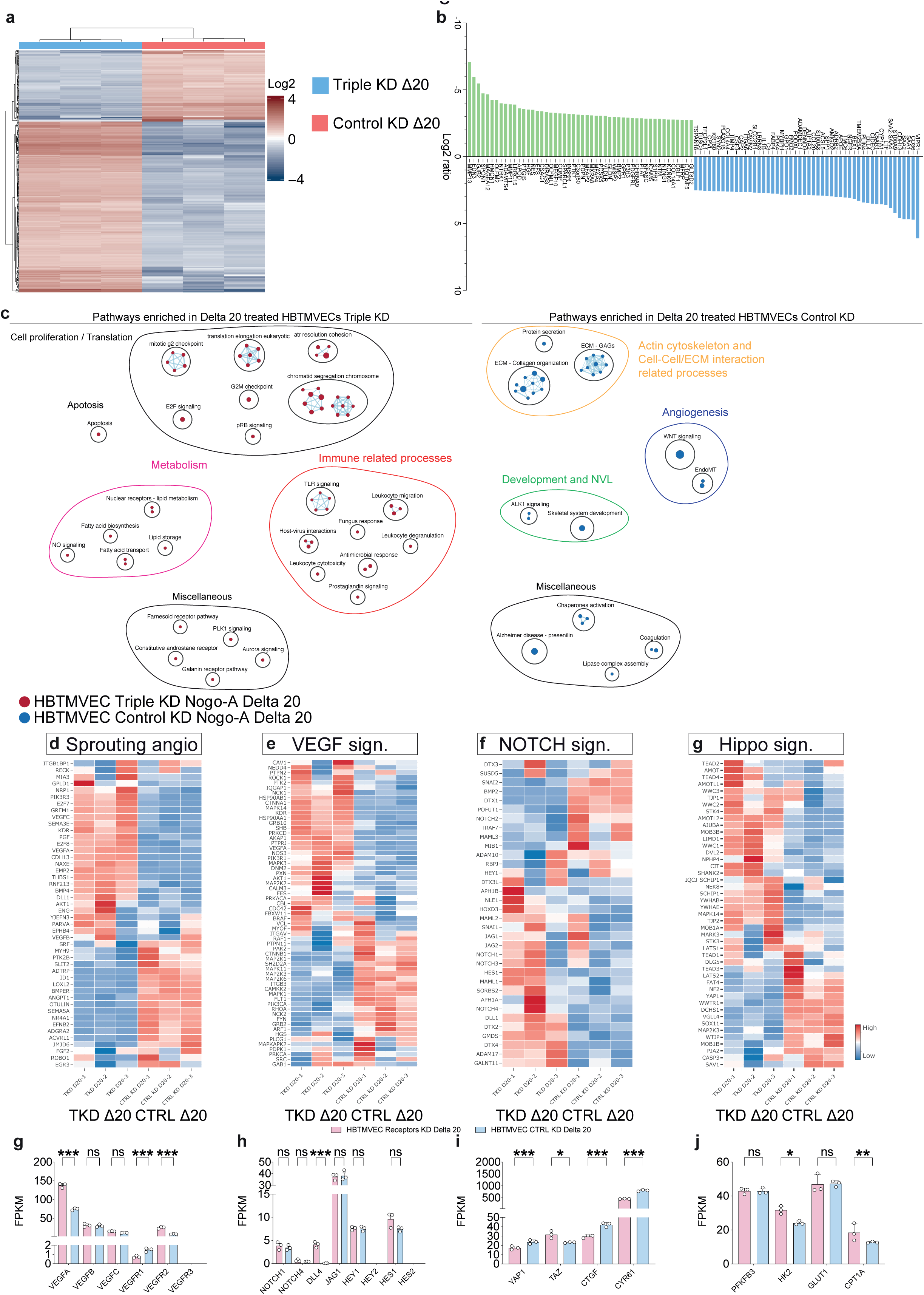
Nogo-A induces regulation of angiogenic pathways including Dll4-Jagged-Notch-Hey-Hes-, YAP-TAZ-CTGF-Cyr61-, VEGF-A-VEGFR2 and endothelial glucose metabolism in HCMECs. Transcriptome analysis via RNA sequencing of HBTMECs treated with short hairpin RNA (shRNA) against Nogo-A receptor complex S1PR2, Tetraspanin 3 (TSPAN3) and Syndecan 4 (SDC4, triple receptor knock-down, TKD) (and treated with Nogo-A Delta 20 compared to control shRNA (shControl, Control KD) and Nogo-A Delta 20 in three independent experiments. (**b**) Heatmap and hierarchical clustering of Nogo-A Delta 20 treated HBTMVECs TKD as compared to Nogo-A Delta 20 treated HBTMVECs Control KD. (**c**) Top 50 significantly up (blue)-or down (green)-regulated genes detected by RNA-sequencing in HBTMVECs TKD upon Nogo-A Delta 20 treatment as compared to Nogo-A Delta 20 treatment control HBTMVECs. Differentially regulated genes were arranged according to fold change of gene expression. Note that the Notch target gene *HEY1* was among the top up-regulated genes, while five of the top down-regulated genes were Hippo-YAP-TAZ target genes. (**c**-**g**) Gene set enrichment analysis (GSEA), followed by cytoscape enrichment map visualization (**c**), showed a significant up-regulation of signaling pathways related to cell proliferation, fatty acid metabolism and immune related processes (**c**) in HBTMVECs TKD treated with Nogo-A Delta 20. Whereas pathways involved in Wnt signaling and cell-cell/ECM interactions (**c**) were enriched in the Nogo-A Delta 20 treated shControl HCMECs. Pathways enriched in Nogo-A Delta 20 treated HBTMVECs TKD are labeled in red, and pathways enriched in control KD HBTMVECs treated with Nogo-A Delta 20 are labeled in blue. Pathways are indicated by colored nodes. Their size represents the number of genes they contain. Green lines indicate relationships between the pathways. Black circles group related pathways. (**d**-**g**) Heatmaps showing the expression of the genes involved in the pathways enriched in HBT previously namely, sprouting angiogenesis (**d**), VEGF signaling (**e**), Notch signaling (**f**), and Hippo signaling (**g**).

Next, to further address the molecular pathways regulated by Nogo-a Delta 20 in brain (tumor) ECs, we performed a pathways analysis between Nogo-A Delta 20 treated HBTMVECs^Triple KD – Nogo-A Delta20^ and the control Nogo-A Delta 21 treated HBTMVECs^Triple KD – Nogo-A Delta21^. In HBTMVEC ^Triple KD – Nogo-A Delta20^, GSEA followed by cytoscape pathway visualization^96^ revealed that upregulated genes were mainly involved in regulation of endothelial metabolism and cell proliferation, indicating rescue of these Nogo-A Delta 20 induced regulatory effects upon Nogo-A Delta 20 triple receptor knockdown (Figure 9c).

Upon examination of genes driving the enrichment of regulation of sprouting angiogenesis, and the VEGF-VEGFR, JAG1-DLL4-NOTCH, Hippo-YAP-TAZ pathways in brain tumor ECs, we observed partial reversal/rescue of various genes involved in these key angiogenic signaling axes (Figure 9d-j), again indicating the involvement of the S1PR2/TSPAN3/SDC4 triple receptor complex in Nogo-A Delta 20 mediated regulatory effects on angiogenic sprouting and its underlying molecular mechanisms in brain tumor ECs.

Together, these results support a crucial role for the S1PR2/TSPAN3/SDC4 receptor complex in mediating the inhibitory regulatory roles of Nogo-A Delta 20 on brain tumor/GBM ECs vascular endothelial cells.

### Endogenous Nogo-A expression within the NVU negatively correlates with human glial brain tumor vessel density, malignancy and progression and with patient survival in IDH mutant low-grade gliomas

To address endogenous Nogo-A expression in human glial brain tumor progression (from WHO low-to high-grade tumors^83,84^) and to investigate whether we could observe a negative regulatory effect on angiogenesis in human glioma patients, we referred to tissue microarrays (TMAs) of human gliomas comprising 103 low– and high grade tissue cores and four cores of normal brain tissue stained immunohistochemically for Nogo-A and the nuclear marker Mayer’s hemalum (Figure 10a-f). Endogenous Nogo-A expression was markedly upregulated in human glial brain tumors as compared to the adult normal brain (Figure 10a-g). Moreover, Nogo-A expression was significantly increased during glial brain tumor progression, ranging from a Nogo-A H-score of 8.13 in the adult normal brain, to 28.58 in WHO grade I glioma, and to 99.27 in WHO glioma grade IV (= glioblastoma) (Figure 10a-g). Nogo-A showed a significant upregulation from low-grade (WHO grade I and II) to high-grade glioma (WHO grade III) as well as a significant increase from WHO grade III to WHO grade IV glioma (= glioblastoma) (Data not shown). In recurrent glioblastoma, Nogo-A expression showed a slight but significant decrease as compared to primary glioblastoma (Figure 10e-g).

**Figure 10:**
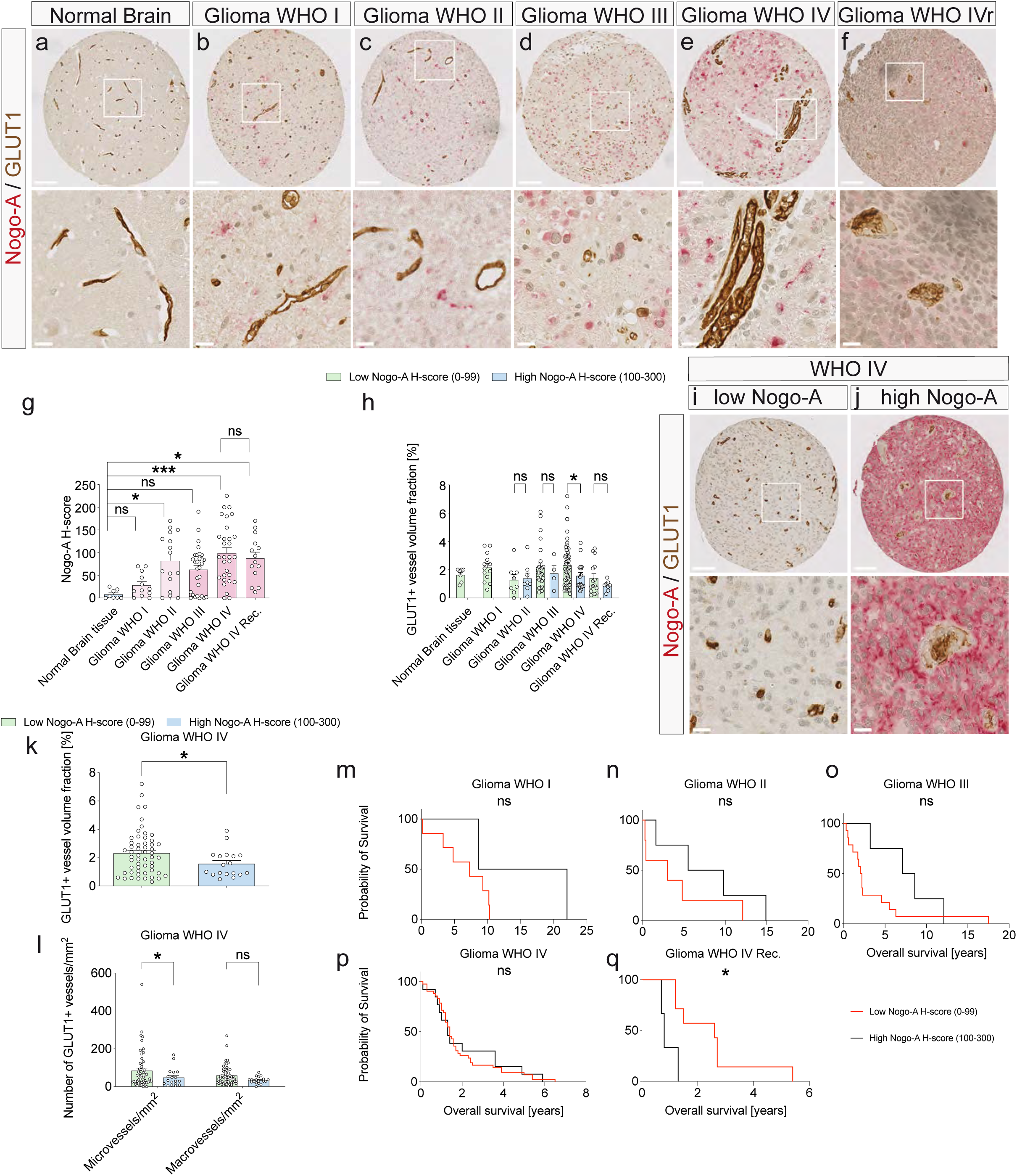
Endogenous expression of Nogo-A inversely correlates with vessel density during tumor progression of human astrocytomas. (**a**-**g**) Nogo-A (red) expression increases during tumor progression of human astrocytomas. Nogo-A expression was addressed using the Nogo-A H-score: *(percentage of cells expressing no Nogo-A * 0) + (percentage of cells expressing low Nogo-A * 1) + (percentage of cells expressing medium Nogo-A * 2) + (percentage of cells expressing high Nogo-A * 3*); Nogo-A H-score ranges from 0-300 (**g**). In the normal human brain parenchyme (n=8), Nogo-A shows weak expression (**a,g**). In low-grade astrocytomas (WHO grades I and II, n= 12 and 16), Nogo-A expression is higher than in the normal brain parenchyme (**b,c,g**), but lower as compared to high-grade astrocytomas (WHO grades III and IV, **d,e,g**, n= 29 and 30). Nogo-A expression decreases again in recurrent WHO grade IV (**f,g**, n=14). (**h**) Analysis of GLUT1 vessel volume fraction in low Nogo-A expressing (H-score 0-99) and high Nogo-A expressing (H-score 100-300) adult control brain, low-grade glioma and high-grade glioma samples. Adult control brain and WHO I glioma didn’t show high expressing samples. GLUT1 vessel volume fraction was not affected by Nogo-A expression in WHO II glioma, but tend to decrease in high Nogo-A expressing higher-grade glioma, with glioblastoma showing a significant decreased in vessel density with higher Nogo-A expression (**h**,**k**). Nogo-A expression displays a high variability in human astrocytomas WHO IV (glioblastoma), ranging from low expression (**i**) to very high expression (**j**). (**l**) GBM samples expressing high levels of endogenous Nogo-A showed a significant lower number of small caliber vessels as compared to samples with low Nogo-A expression. Nogo-A expression slightly reduced the number of large caliber vessels in glioblastoma. (**m**-**q**) Kaplan-Meier curves indicating overall survival (in years) of patients with high and low-expressing Nogo-A low grade gliomas (**m,n**), high grade gliomas (**o,p**) and recurrent glioblastomas (q, n=). High Nogo-A expression tend to be associated with increased patient survival in low grade but showed a significant negative correlation in recurrent glioblastoma patient survival. Primary high grade glioma survival wasn’t affected by endogenous Nogo-A expression. (**r**-**e_i_**) Correlation analysis between Nogo-A expression (via two *RTN4* isoforms 202 and 209) and prognosis of glioma patients from TCGA. Nogo-A expression was significantly higher in low grade glioma (**r**,**y**), but significantly lower in GBM (**w**,**d_i_**) as compared to adult control brain. The expression between IDH1 wild type (WT) and IDH1mutant LGG/GBM was not changed. Kaplan–Meier overall survival plot showed increased overall survival for patients with high Nogo-A expression in LGG (**s**,**z**) and LGG IDH WT (**u**,**b_i_**) but not in LGG IDH WT (**v**,**c_i_**) and GBM (**x**,**e_i_**) in TCGA dataset. Data represent mean ± SEM. \**P* < 0.05, \*\*\**P* < 0.001. Scale bars represent: 100 µm in **a**-**f** (upper pictures) and in **i**-**j** (upper pictures); 20 µm in **a**-**f** (lower pictures) and in **j-k** (lower pictures)

Based on the expression of Nogo-A within perivascular cells of the developmental– and tumoral NVU in close proximity to tumor blood vessels and given its negative regulators effects on human and mouse brain (tumor) blood vessel endothelial cells in vivo and in vitro, we next addressed whether Nogo-A affected on human glioma blood vessel density. Therefore, we examined the vascular volume fraction of GLUT1^+^ and CD31^+^ tumor blood vessels in low– and high-Nogo-A expressing gliomas across glial brain tumor progression. Notably, we observed a negative correlation between Nogo-A expression and blood vessel VVF, as evidenced by a significantly decreased VVF in gliomas expressing high Nogo-A (Figure 10h and Supplementary Figure 7g), an effect that was most prominent in GBMs (Figure 10h-k and Supplementary Figure 7g-j). Furthermore, based on the inhibitory effects of Nogo-A on ETCs at the forefront of growing blood vessels during brain development and in brain tumors and in light of our previous observation that Nogo-A exerts its inhibitory regulatory effects on angiogenesis and vascular network formation during postnatal brain development predominantly at the level of capillaries^58^, we next addressed the effects of Nogo-A on capillary versus non-capillary vessel density in glioblastoma. Interestingly, Nogo-A expression negatively correlated with the vessel density of GLUT-1 in glioblastoma (Figure 10h-k), an effect that was significant for microvessels (= capillaries) and showed a non-significant trend for macrovasssels (non-capillaries) (Figure 10l and Supplementary Figure 7k), thereby further enforcing Nogo-A’s predominant inhibitory effect on capillaries, also in human glioblastoma angiogenesis. Taken together, these data indicate that Nogo-A expression is reactivated in tumor cells in vicinity of tumor endothelial cells within the NVU and negatively correlates with tumor vessel density during human astrocytic tumor progression.

We next wondered whether Nogo-A expression correlated with glioma patient survival in the TMA data. Survival analysis of these 103 TMA samples revealed a trend indicating that Nogo-A expression levels were associated with clinical prognosis in lower-grade gliomas. Indeed, in lower-grade gliomas (WHO I and WHO II), the median survival time of patients with high Nogo-A expression tend to be longer than of patients with low Nogo-A expression (Figure 10m-q). However, based on the relatively low n numbers per glioma grade in the TMA and in order to further evaluate the clinical significance of Nogo-A expression in gliomas, we referred to the glioma cancer datasets in The Cancer Genome Atlas (TCGA)^111,112^ to further analyze glioma survival of 662 total glioma cases (105 normal adult brains, 509 low-grade gliomas and 153 high-grade gliomas).

Because the Nogo-A protein is encoded by the RTN4 gene which encodes also for Nogo-B and Nogo-C, we examined the two RTN4 gene isoforms 202 and 209 because they feature exon 3 (that includes the Nogo-A-specific fragment Delta 20), which is only present in the Nogo-A protein but not in the Nogo-B and Nogo-C proteins. Thus, we addressed the expression of RTN4 gene isoforms 202 and 209 in gliomas by examining the RNA-seq data in TCGA database.

In comparison to the normal adult brain, the mRNA expression of Nogo-A (RTN isoforms 202 and 209) was significantly upregulated in the LGGs and significantly downregulated in the HGGs/GBMs. Interestingly, the observed discrepancy between the Nogo-A expression in HGGs/GBMs between our TMA and TCGA data might be due to post-transcriptional modifications given the implication of splicing factors PTBP1 and PTBP2 in the regulation of RTN4 splicing in glioma cells^113^ (Figure 10r,u,c_i_k_i_).

Next, to correlate expression of Nogo-A isoform genes with low-and high-grade glioma patient survival, we took advantage of the large bulk RNA-seq dataset of 1662 glioma patients and 105 normal adult brains of the TCGA. Notably, lower-grade (but not high-grade/glioblastoma) glioma patients, who expressed high levels of the Nogo-A gene isoforms 202 and 209 showed significantly longer overall survival as compared to those expressing low levels of the Nogo-A gene isoforms 202 and 209 (Figure 10s,t,d_i_,e_i_), presumably because Nogo-A inhibits active glioma angiogenesis and thus glioma growth. However, the survival benefit for patients expressing high levels of the Nogo-A gene isoforms 202 and 209 was not observed for HGG/GBMs (Figure 10 a_i_,b_i_,l_i_,m_i_). Notably, in HGGs/glioblastomas, whereas higher Nogo-A expression did positively correlated (not reaching significance) with patient survival for the Nogo-A isoform 202 (Figure 10a_i_,b_i_), expression of the Nogo-A isoform 209 negatively correlated with patient survival (Figure 10l_i_,m_i_). We next assessed Nogo-A expression and its correlation with patient survival in the TCGA low-grade glioma– glioblastoma database, in glioblastoma by grouping LGGs and HGGs/glioblastomas by the key defining molecular feature being the IDH mutation^7,8^ (Figure 10u-y, f_i_-j_i_). Survival analysis of IDH mutant LGG samples demonstrated that Nogo-A expression was significantly associated with clinical prognosis as the median survival time of patients with high Nogo-A isoform 202 expression was significantly longer than patients with low or medium Nogo-A expression (Fig. 10v,x), whereas the survival of IDH mutant LGGs was not significantly correlated with Nogo-A expression (Fig. 10w,y). For the Nogo-A/RTN4 isoform 209, we again found that IDH WT LGG patients with tumors with high Nogo-A expression exhibited a better prognosis than patients with a low Nogo-A expression (Figure 10g_i_,i_i_). For both Nogo-A isoforms 202 and 209, IDH WT LGG patients with low Nogo-A expression patterns fared the worst (Figure 10b), supporting the finding that Nogo-A expression is a major predictor of favorable LGG patient survival provided absence of the IDH mutation. For HGGs/glioblastomas, stratification according to the IDH mutation was not possible as only very few glioblastomas harbored the IDH mutation, in agreement with the new WHO classification, in which the presence of the IDH classifies at glioma as not being a GBM.

These results are consistent with the observations derived from the TMA dataset, which revealed that LGGs with a higher Nogo-A expression exhibited a (even though non-significant) trend towards a higher patient survival than LGGs with low Nogo-A expression. Our data suggest that Nogo-A expression is increased in human gliomas, and its expression positively correlates with survival in IDH WT LGGs which might be linked to its restricting effects on tumor vascularization and progression.

## DISCUSSION

Here, using in vitro and in vivo approaches as well as human and mouse brain (tumor) tissues, we characterize the neurite growth inhibitory membrane protein Nogo-A as an onco-fetal protein and show that it negatively regulates angiogenesis/vascularization and tumor growth in human gliomas. Our results suggest that the Nogo-A specific domain Delta 20 inhibits brain (tumor) endothelial sprouting, migration, spreading, and filopodia formation via interaction with the VEGF-VEGFR, Dll4-Jagged-Notch, YAP-TAZ pathways and restricts brain (tumor) endothelial glucose metabolism via the regulation of glycolytic regulators including GLUT1. We propose that, by acting on the cytoskeleton of brain tumor endothelial (tip– and stalk) cells and their filopodia, and by regulating vascular endothelial metabolism, Nogo-A Delta 20 controls the sprouting and filopodia extension of growing CNS blood vessels in human and mouse brain tumors. Moreover, we show that Nogo-A Delta 20 promotes brain tumor vessel normalization and reduces intratumoral hypoxia (Figure 11). Strikingly, endogenous Nogo-A expression negatively correlates with human glioma vascularization while positively correlating with IDH-WT low-grade glioma patient survival. Importantly, the characterization of Nogo-A as an onco-fetal protein in the human and mouse brain tumor vasculature and its inhibitory effects identifies Nogo-A as a potential pharmaceutical candidate for human gliomas.

**Figure 11.**
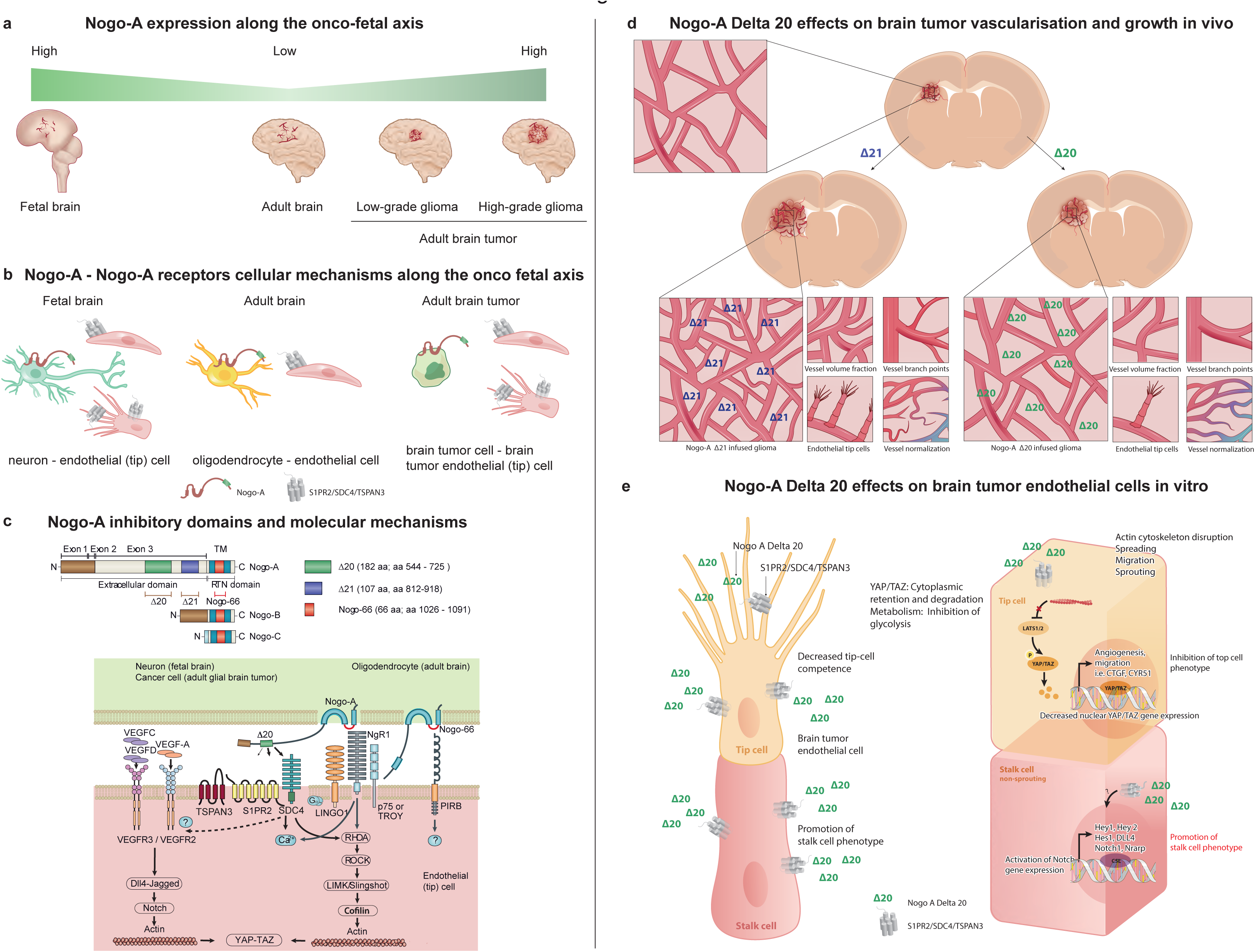
Nogo-A – S1PR2/TSPAN3/SDC4 is an onco-fetal signaling axis regulating angiogenesis during brain development and in brain tumors – working model.

### Nogo-A – a NVU/PVN derived onco-fetal NVL signal to regulate developmental brain– and brain tumor (vascular) growth in mice and humans

The bulk of evidence regarding the molecular regulation of sprouting angiogenesis during brain development and in brain tumors is based on murine studies, whereas much less knowledge exists about how vascularization and ETCs are regulated during human brain development and in human (glial) brain tumors. Yet, to achieve a comprehensive understanding of developmental and tumor angiogenesis that harbor translational potential, integrative studies referring to mouse and human models are required^4,5,114^.

In both mice and humans, angiogenesis is highly dynamic during brain development, enters an almost quiescent state in the adult healthy brain^19,29,33^, and is reactivated in a variety of angiogenesis-dependent CNS pathologies such as brain tumors, brain vascular malformations, or stroke^1,2,33^, thereby activating endothelial– and perivascular cells of the NVU^15,19,29^. In our study, we observed Nogo-A expression in perivascular cells in vicinity of angiogenic endothelial (tip) cells in the human fetal brain and in glial brain tumors. Moreover, genetic deletion of Nogo-A or S1PR2 resulted in increased length of ETC filopodia without affecting the numbers of ETC filopodia, indicating Nogo-A’s negative regulatory role on angiogenesis during mouse brain development, in agreement with. Moreover, Nogo-A restricted angiogenesis and ETCs in mouse glioma as well as a negative correlation of Nogo-A expression negatively correlates with astrocytic brain tumor vascularization, progression, and survival, in agreement with^115^, These mouse and human data suggest a crucial role of Nogo-A as a perivascular niche-derived inhibitory signal for vascularization and ETCs in both mouse and human brain development and brain tumors. Notably, the perivascular niche/neurovascular unit has been shown to activate/stimulate tumor growth/development in mouse– and zebrafish models of breast cancer^51^. Accordingly, whilst the stable microvasculature provided anti-angiogenic cues, the active microvasculature promoted breast cancer cell growth via a direct cellular endothelial-to AND tumor cell crosstalk. Strikingly, within the stable breast cancer microvasculature hrombospondin-1 derived from the endothelium induced sustained breast cancer cell quiescence, whereas in sprouting neovasculature, this suppressive cue was lost and in addition, ETC-derived active TGF-1 and β periostin promoted breast tumor growth^51^. In accordance, the stable microvasculature is characterized as a “dormant perivascular (tumor) niche”, while the sprouting neovasculature, is referred to as an “activated perivascular (tumor) niche” in which ETCs play important roles.

In light of these studies, the further exploration of Nogo-A’s inhibitory function on angiogenesis within the perivascular tip cell niche in vivo as well as of its therapeutic relevance promises to provide further insights into these exciting concepts.

Here, we found that Nogo-A is expressed in the human fetal brain and in human glial brain tumors and acts as an important negative regulator of angiogenesis and ETC filopodia while promoting vascular normalization in human and mouse glial brain tumors. The expression of Nogo-A in – in perivascular cells such as neurons, astrocytes and pericytes in vicinity of ETCs within the neurovascular unit of the human fetal brain, its downregulation in the adult healthy brain and reactivation in glial brain tumors characterizes Nogo-A as an onco-fetal protein and suggests an integral role once reactivated during brain cancer (Figure 11). The high Nogo-A expression in vicinity of CD105^+^ angiogenic blood vessel ECs in both human fetal brain and human glioblastoma supports/corroborates the presumed role of Nogo-A as a “tonic” inhibitor of active (developmental, tumor) versus stable (adult healthy) brain (tumor) angiogenesis. Accordingly, our in vivo results in Nogo-A KO (and in S1PR2 KO) as well as the in vitro results from various functional bioassays suggest/indicate that by exerting/applying negative regulatory effects on sprouting angiogenesis, cell migration, spreading, filopodia extension, and glucose metabolism of vascular brain ECs, Nogo-A Delta 20 exerts inhibitory regulatory effects on the sprouting of angiogenic blood vessel ECs into the brain parenchyma during brain development and in glial brain tumors. VEGF-A, Wnt7a, GPR124 are other neurodevelopmental regulators downregulated in the adult healthy brain and reactivated in vascular-dependent CNS pathologies including brain tumors or stroke^31,116,117^. In contrast to those angiogenic factors, however, Nogo-A is one of the few onco-fetal proteins characterized and compared in both human and mouse brain development and brain tumors, similar to our previous characterization of Nucleolin as an onco-fetal protein. The perivascular expression pattern of Nogo-A within the fetal– and tumor neurovascular unit/perivascular niche and its inhibitory roles on both endothelial and non-endothelial cell types during brain development (as well as in (glial) brain tumors identify Nogo-A as an onco-fetal protein in the human brain vasculature with inhibitory roles during brain development and in glial brain tumors.

### The onco-fetal axis including Nogo-A as novel therapeutic target in the brain tumor vasculature?

A better understanding of brain tumor biology and treatment requires a thorough characterization of the tumor microenvironment (TME) composed of the extracellular matrix as well as blood vessel endothelial– and perivascular cells^4–6,15,22,89,118,119^. Mutual interactions between the TME vascular and the immune system are highly relevant in order to achieve improvements in the treatment of brain tumors^15,5,120^.

Recent studies indicating that brain tumors can be seen as the result of aberrant/dysregulated brain development or repair. Moreover, tumors were shown to reactivate onco-fetal programs in the tumor vasculature and the perivascular niche^4–6,15,118,119^ underlining the importance of integrating cancer and neuroscience research. Interestingly, tumors escape immunity by resurrecting a developmental vascular program. Notably, during development, angiogenesis reduces immune cell infiltration by suppressing adhesion molecules on ECs to induce immune-privileged conditions and uninterrupted growth. Angiogenesis inhibitors, in consequence, overcome this state of compromised immunity / immune-compromised state and thereby improve immunotherapy^41^.

Infiltration of immune cells into tumors requires homing of immune cells, subsequent adhesion to the tumor endothelium, and extravasation to the tumor microenvironment to mediate immune responses^91^. Adhesion molecules on the tumor endothelium include intercellular adhesion molecule 1 (ICAM1), vascular cell adhesion molecule 1 (VCAM1), and CD34^91^.

Pro-angiogenic molecules like VEGF, FGF2, or EGF downregulate the expression of ICAM1, VCAM1 and CD34 on endothelial cells in non-tumor and tumor tissues leading to inhibition of leukocyte adhesion via different mechanisms^91^, whereas anti-angiogenic cues including endostatin and angiostatin are capable of reversing the angiogenesis-driven downregulation of ICAM1, VCAM1, and CD34 expression, thereby leading to increased CD8^+^ T cell infiltration into tumors^91^. These observations not only emphasize the relevance of fetal reprogramming in tumor cells^41,42^ but also highlight the mutual dependency between (anti-)angiogenesis and immunosuppression/-stimulation^41^.

Interestingly, in our transcriptomic analysis of human brain and brain tumor endothelial cells treated with Nogo-A Delta 20, we observed an increase in VCAM1 and ICAM1 (but not CD34) expression, suggesting that – together with the observed inhibitory effects of Nogo-A Delta 20 on VEGF-signaling – Nogo-A may upregulate tumor endothelial cell adhesion molecules such as VCAM1 and ICAM1 via its anti-angiogenic effects.

In summary, based on the observed expression of Nogo-A in the developing fetal and glial brain tumor vasculature and given its positive regulatory role on the expression of the adhesion molecules VCAM1 and ICAM1 (but not CD34) on glioblastoma tumor endothelial cells, a tempting possibility is that Nogo-A’s anti angiogenic role might counteract the immunosuppressive TME in glioblastomas resulting in a more immunostimulatory TME thereby contributing to improved immunotherapies for brain tumors^121^.

### Nogo-A – an NVL general mechanisms of angiogenesis restricting brain tumor vascularization and promoting brain tumor vascular normalization – combination with immunotherapy?

In various cancer types, immunotherapies have markedly improved patient outcomes^91^, but in malignant brain tumors, and particularly in primary gliomas, immunotherapy has had little impact in tackling these deadly disease.

The combination of immunotherapies with anti-angiogenic agents harbors the potential to improve therapy response rates and duration as drugs targeting VEGF, angiopoietin 2 or HGF signaling are capable of increasing the efficacy of immunotherapies. Moreover, as immunotherapies, and in particular immune-checkpoint inhibitors, might increase the efficacy of anti-angiogenic therapies and/or promote changes in the tumor vasculature, interactions between immune– and anti-angiogenic therapeutic strategies are being considered a ‘two-way street’.

Immunotherapies are most effective against tumors with an inflammatory microenvironment, but (glial) brain tumors are ‘cold’ or ‘immunologically ignorant/compromised’ tumors, given their lack of active infiltrating lymphocytes (TILs). Because the limited presence/lack of immune-cell infiltration into tumors compromises the efficacy of immunotherapies, strategies to promote immune-cell infiltration into tumors and to revert the effects of an immunosuppressive tumor microenvironment are being explored and include anti-angiogenic therapies.

As outlined above, angiogenesis and immunosuppression are closely related processes that can occur in parallel, for instance facilitating both normal tissue and tumor development and progression. To that regard, various pro-angiogenic factors and especially VEGF also have immunosuppressive functions and anti-angiogenic agents targeting these pro-angiogenic factors can stimulate an immune response, which can enhance anti-tumor immune responses. Here, we observed negative regulatory effects of Nogo-A on the pro-angiogenic VEGF-VEGFR, Hippo-YAP-TAZ and endothelial glucose metabolism signaling axes in brain tumor endothelial cells in vitro and on Hippo-YAP-TAZ signaling in the mouse glioma vasculature in vivo.

While pro-angiogenic molecules are associated with immunosuppressive effects at successive steps in the cancer-immunity cycle, including antigen presentation, T cell priming, T cell trafficking, and T cell tumor infiltration, regulators of angiogenesis affect immune cells and their interaction with tumors in multiple ways including direct effects when binding to their cognate receptors expressed by immune cells as well as indirect effects when inducing protein expression changes on endothelial cells and promoting tumor vascular normalization or reducing tumor neoangiogenesis.

Here, we observed normalization of the brain tumor vasculature upon Nogo-A Delta 20 infusion with concomitant reduction of brain tumor vessel density, sprouting, branching and ETC number in vivo, as well as a decrease of brain tumor endothelial cell sprouting, migration, spreading, lamellipodia and filopodia extension and endothelial cell proliferation in vitro (Figure 11). These are key findings because the (brain) tumor vasculature features many abnormal characteristics including increased tortuosity and a lack of an ordered structure, which is often caused, at least partially, by constant sprouting angiogenesis (without concomitant vascular remodeling and maturation), by tumor cell-mediated compression of blood vessels and by higher rates of (brain) tumor endothelial cell proliferation as compared to normal tissues. Moreover, such aberrant tumor vessels lead to disrupted immune-cell infiltration, decreased vascular perfusion, and the emergence of intratumoral hypoxic areas.

Tumor hypoxia can induce immunosuppressive effects on T cells and anti-angiogenic agents can result in vascular normalization, which transiently alleviates intratumoral hypoxia and which revers the tumor vasculature reverts into a more normal vascular network, reminiscent of that of healthy organs. Anti-angiogenic agents can induce vasculature normalization that can exert profound positive effects on immune-cell infiltration into tumors. However, anti-angiogenic agents can also lead to tumor vessel damage and tumor vessel density reduction, and these processes can intensify tumor hypoxia and reduced immune cells and penetration of therapeutics into the tumors. Based on our observations, Nogo-A Delta 20 treatment results in anti-angiogenic effects including decreased vessel density, reduced vessel diameter and reduction in ETC numbers and at the same time tumor vascular normalization and reduction of hypoxia. These properties characterize Nogo-A Delta 20 as potentially interesting anti-angiogenic and vessel normalizing agent that could be tested respecting an optimal window of tumor vascular normalization, and potentially in combination with other agents. To that regard, determining the optimal therapeutic dose of Nogo-A Delta 20 (as well as its mode of delivery, e.g. intratumoral post-resection or intravenous will be crucial, as vascular normalization upon low dose-treatment with anti-VEGFR2 antibodies was found to be associated with increased T cell infiltration in mouse models of breast cancer and presumably in glial brain tumors. indicating that the dose of anti-angiogenic agents might affect vessel normalization and in consequence the immune response.

Current strategies to promote vascularization in CNS and non-CNS tumors include blockade of angiopoietin-2 leading to vessel normalization characterized by decreased vessel diameters and increased pericyte coverage as well as to reduced sprouting angiogenesis and redistribution of junctional proteins including platelet endothelial cell adhesion molecule (PECAM1) and vascular endothelial cadherin (VE-cadherin), features reminiscent of vessels in healthy tissues. Interestingly, antibody-mediated neutralization of angiopoietin 2 in a xenograft GL261 glioma model leads to enhanced TIE2 activation resulting in vessel normalization thereby facilitating the delivery of chemotherapeutic agents into tumor tissues. Here, we observed concomitant inhibition of sprouting angiogenesis and vessel density with vascular normalization characterized by a trend for decreased vessel diameter while pericyte coverage remained unchanged. These observations together with the finding that Nogo-A signaling restricts VEGF-VEGFR signaling are especially interesting given that the combined blockade of VEGF and angiopoietin 2 is the most effective way to promote vessel normalization. As VEGF blockade alone results in transient normalization of vessels and angiopoietin 2 blockade result in a more robust and lasting normalization with heightened pericyte association and enhanced cell–cell contacts, only the dual blockade inhibits angiogenesis by reducing vessel density and promoting vascular normalization thereby improving immune effector cell extravasation and the efficacy of ICIs; and resulting in reprogramming of the tumor vasculature.

As Nogo-A Delta 20 infusion promotes vascular normalization while inhibiting sprouting angiogenesis and vessel density, it might constitute an ideal therapeutic candidate to support immunotherapies. As discussed previously, certain modulators of angiogenesis, especially VEGF, have a broad range of diverse effects on the immune system that are mainly immunosuppressive. Because blocking angiogenic molecules using a strategy based on a single therapeutic approach is likely insufficient to generate a complete or robust immune response against cancer especially in patients with advanced-stage disease. Therefore, exploring combinations of anti-angiogenic agents such as Nogo-A Delta 20 and anti-VEGF antibodies with various immunotherapies that synergize and increase adaptive immune responses.

### Nogo-A Delta 20 restricts mouse and human glioma vascularization via a Hippo-Yap-Taz-NOTCH signaling axis

Our in vitro results suggest that the effects of Nogo-A on developmental brain and brain tumor angiogenesis are mediated by negative regulatory effects on brain (tumor) endothelial sprouting, –migration, –spreading, –glucose metabolism –and tip cell filopodia formation. Our in vivo findings showing Nogo-A expression in vicinity of brain endothelial tip-stalk– and phalanx cells and revealing restricting effects on brain tumor ETCs and developing brain ETC filopodia further support this/our working model (Figure 11). Whereas our in vivo GL-261, in vivo P8 postnatal brain as well as our in vitro spheroid and nanopillar array data mainly indicate a negative regulatory role for Nogo-A Delta 20 in endothelial tip cell (filopodia) formation, we observed an inhibitory effect of Nogo-A Delta 20 on endothelial cell proliferation, suggesting a possible regulatory effect on stalk cell proliferation in vivo. However, the relative importance of Nogo-A for endothelial tip-, stalk-, and phalanx-cell types and cellular mechanisms in vivo and in vitro remains obscure. For instance, differential expression patterns of Nogo-A in brain development, and in brain tumors versus the “healthy” surrounding brain certainly influence the competitiveness of brain ECs to acquire the tip cell position^50,122^ and vascular sprout formation. Nogo-A’s in vivo role on endothelial stalk cell proliferation using Ki-67 markers in embryonic and postnatal brain angiogenesis and in brain tumor angiogenesis mouse models should be addressed. Furthermore, Nogo-A’s putative role on tip versus stalk cell specification using in vitro genetic mosaic sprouting angiogenesis assays^50^ and *in silico* computational simulation^122,123^ should be assessed. These experiments could also clarify the molecular mechanism that mediate our interesting observation of a the observed increased ETC filopodia length and decreased straightness upon Nogo-A KO (and S1PR2 KO) during postnatal mouse brain development in vivo. Investigating Nogo-A’s role on migrating tip cells, proliferating stalk cells and quiescent phalanx cells as well as tip vs. stalk cell specification in vivo in both brain development and brain tumors therefore promises to further our understanding of these interesting observations.

Actin cytoskeleton-based endothelial tip– and stalk cell filopodia– and lamellipodia formation is crucial during vascular sprout formation and guidance, as well as endothelial cell migration^19,29,46,47,94,124^, and various NVL molecules including Nogo-A have crucial regulatory effects on endothelial tip– and stalk cell filopodia/lamellipodia during brain development and in brain tumors. The genetic ablation of Nogo-A and of the Nogo-A receptor complex component S1PR2 KO in postnatal mouse brain development led to increased length and disoriented ETC filopodia in vivo and Nogo-A Delta 20 gain-of-function lead to decreased number of ETCs in brain tumor angiogenesis in vivo and to retraction of actin cytoskeleton-based lamellipodia and filopodia protrusions of brain (tumor) ECs in vitro. Notably, recent evidence suggests that ROCK decreases EC filopodia-like protrusion length in vitro, which might explain the above-mentioned phenotypes given the Nogo-A induced Rho-A-ROCK activation in neuronal, non-neuronal as well as brain endothelial cells.

### Nogo-A as negative regulator of glucose-, but not fatty acid brain (tumor) endothelial metabolism – effects on endothelial tip– and stalk cells?

In both development and in tumor, endothelial metabolism acts as crucial regulator of sprouting angiogenesis ^90,95,98,99,104,125^, and while endothelial tip cells predominantly rely on glycolysis, endothelial stalk cells also rely on fatty acid metabolism to support endothelial cell proliferation^90,95,98,99,104,125^. We found here that Nogo-A negatively regulates brain (tumor) endothelial glycolysis-but not fatty acid metabolism in vitro, suggesting that Nogo-A predominantly affects endothelial tip cells. These findings are, on the one hand, supported by the negative regulatory effects of Nogo-A on ETCs and their filopodia during postnatal mouse brain development and in mouse glial brain tumors in vivo as well as the inhibitory effect of Nogo-A on the actin cytoskeleton in vitro. Moreover, while the formation of filopodia depends on the actin cytoskeleton, glycolytic production of ATP promotes filopodia formation in vivo, in vitro, and in silico^122^. On the other hand, however, based on the observed expression of Nogo-A in vicinity of both endothelial tip– and stalk cells in vivo as well as in light of the negative regulatory effect on sprouting angiogenesis (an endothelial tip cell function) as well as on endothelial proliferation (an endothelial stalk cell function) in vitro, the precise roles of Nogo-A and its receptor complex S1PR2/TSPAN3/SDC4 on both tip– and stalk cells in– and outside the CNS, as well as in the human fetal brain, in the mouse embryonic or postnatal brain^29,43^ and in the postnatal retina^126^, as well as in mouse and human glial brain tumors need to be further examined in vivo.

As evidenced by our bulk RNA sequencing data, signaling axes regulated downstream of Nogo-A were linked to angiogenesis and endothelial proliferation. The observed negative regulatory effects of Nogo-A on brain (tumor) endothelial glucose metabolism are in line with the crucial role of glycolysis in angiogenesis and vascular biology in (brain) development and tumor in as well as outside the CNS^90,98,99,127,128^. Nevertheless, we cannot rule out the possibility that Nogo-A also regulates other metabolic pathways that contribute to brain (tumor) angiogenesis and brain (tumor) EC biology, underlining the need for future studies aimed to determine the precise in vivo and in vitro roles of Nogo-A on brain (tumor) EC metabolism and angiogenesis.

Notably, upon Nogo-A Delta 20 addition, we observed a downregulation of the VEGF-A-VEGFR2 signaling pathway at the mRNA level as well as to a decrease of p-VEGFR-2 and VEGFR-2 at the protein level, indicating an inhibitory effect of Nogo-A on the canonical VEGF-A-VEGFR2 signaling axis in brain and brain tumor endothelial cells, in line with what was observed in the peri-infarct area after stroke. Interestingly, VEGF-A mRNA was upregulated upon Nogo-A Delta treatment in HBTMVECs and downregulated in HBMVECs, and these seemingly contradictory findings may be due to the complexity of the VEGF – VEGFR signaling system also involving compensatory mechanism^129^, due to translational modifications^130,131^, and also due to the capacity of brain tumor (but not normal brain) endothelial cells to counteract anti-angiogenic stimuli with pro-angiogenic cues. Given the crucial role for the VEGF-VEGFR signaling system for CNS angiogenesis^92^ and ETCs^1,21,33,132,133^ during brain development and in brain tumors, and given that Nogo-A inhibition reduced VEGF-A-VEGFR2 driven BBB permeability after stroke, examining the precise molecular interactions between the VEGF-VEGFR and the Nogo-A – Nogo-A receptor complex signaling pathways in brain tumor vascularization in vivo harbors the potential to further characterize Nogo-A as a therapeutic agent for malignant gliomas. The very same holds true for further investigations regarding the molecular interactions of Nogo-A – Nogo-A receptor signaling with endothelial metabolism, given the crucial role for vascular endothelial (glucose and fatty acid) metabolism for vessel sprouting in development and disease^90,95,98^ as well as of tumor metabolism in gliomas^57^.

In summary, whether Nogo-A acts as an onco-fetal protein as well as its role on angiogenesis and endothelial cell function in human gliomas remains unknown. Here, using a variety of in vivo and in vitro assays, we characterize Nogo-A as an onco-fetal protein in human glioma vascularization that negatively regulates/restricts sprouting angiogenesis and endothelial metabolism via its S1PR2/TSPAN3/SDC4 receptor complex in human and mouse glioma via molecular crosstalk with the key angiogenic pathways VEGF-VEGFR, Dll4-Jagged-NOTCH, and Hippo-YAP-TAZ, and find that its endogenous expression negatively correlates with human glial brain tumor vascular density and is associated with better survival of IDH-WT low-grade glioma patient.

## ACKNOWLEDGMENTS

We thank our colleagues Giovanna Longo for help with fetal brain sample preparation for immunofluorescence and confocal imaging, Niklaus Krayenbühl and Oliver Bozinov for help with the human adult tissue asservation, Rudolf Steiner for help with the in vivo/ex vivo CAM angiogenesis assay, Jiazhuo He for help with in vitro spheroid assays, Hang Zhong for help with the p-VEGFR2/VEGFR2 immunofluorescence image processing and analysis, Hubert Rehrauer for help with bulkRNAseq analysis, Sebastian Streb and Endre Laczko for help with metabolomics analysis, and Nancy Chu Ji for help with the illustrations.

The author(s) disclosed receipt of the following financial support for the research, authorship, and/or publication of this article: T.W. was supported by the OPO Foundation, the Swiss Cancer Research foundation (KFS-3880-02-2016-R, KFS-4758-02-2019-R), the Stiftung zur Krebsbekämpfung, the Kurt und Senta Herrmann Foundation, Forschungskredit of the University of Zurich, the Zurich Cancer League, the Theodor und Ida Herzog Egli Foundation, the Novartis Foundation for Medical-Biological Research, and the HOPE Foundation. K.D.B. was supported by grants of the Swiss National Science Foundation (31003A-176056) and a starting grant of the European Research council (716140). V.V. was supported by the Swiss National Science Foundation (SNF 310030B_133122/1), SNF NCCR “Molecular Systems Engineering”, the Commission of the European Communities/European Research Council (ERC) Advanced Grant (231157), FIRST and SCOPEM facility from ETHZ. D.V. was supported by Intramural Funding (2019).

## AUTHOR CONTRIBUTIONS

T.W. had the idea for the study, T.W. conceived the study, designed the experiments, wrote the manuscript with the help of M.S, analyzed the data with M.S and M.G., designed the figures and made the figures with M.S. and the help of M.G.. T.W., L.R., K.S., M.B., P.K., G.Z, T.V. acquired the tissue. T.W. and K.F: performed the in vivo/ex vivo CAM angiogenesis assay and T.W., K.F. and A.W. analyzed the data. J.Y. and R.V.D. performed TCGA analyses. D.G. and M.W. performed TMA analyses. M.B. and J.V.B performed the mouse-in-mouse glioma experiments, with the help of T.W., K.F. and MS. M.S. analyzed the mouse-in-mouse glioma experiments with the help of M.G. J.L.J-M. and O.N. performed human-in-mouse glioma experiments. M.G. and J.H. performed the in vitro angiogenesis and signaling assays and M.S. helped with the analysis. T.W. and K.F. developed the initial isolation experiments. T.W., S.L., K.D.B., P.P.M. acquired funding. T.W. and M.S. edited the final version of the manuscript. P.P.M., K.B., J.E.F., M.L.S., P.D., P.C., V.T., G.Z., T.V., I.R., G.B. gave critical inputs to the manuscript. T.W. supervised all the research. All authors read and approved the final manuscript.

## METHODS

### Human fetal, adult brain and brain tumor tissue

Samples of fetal brain were obtained from a 22-week-old post-mortem fetus, derived from spontaneous abortions and received by the Department of Pathological Anatomy, University of Bari School of Medicine. Permission to collect fetal tissue was obtained from the mother at the end of the abortion procedure. The sampling and handling of the specimens conformed to the ethical rules of the Department of Emergency and Organ Transplantation, Division of Pathology, University of Bari School of Medicine, and approval was gained from the local Ethics Committee of the National Health System in compliance with the principles stated in the Declaration of Helsinki. The fetuses did not reveal macroscopic structural abnormalities at autopsy and/or microscopic malformations of the central nervous system after conventional histological analysis with H&E or toluidine blue staining. The fetal age was estimated based on the crown-rump length and/or pregnancy records (counting from the last menstrual period). From each fetus, samples of the dorso-lateral wall of the telencephalic vesicles (n=6; future cerebral hemispheres) were dissected along the coronal plane in slices about 0.5-cm thick, fixed for 2–3 hours at 4°C by immersion in 2% paraformaldehyde (PFA) plus 0.2% glutaraldehyde in phosphate-buffered saline solution (PBS, pH 7.6), washed in PBS and stored in PBS plus 0.02% PFA at 4°C. The parahippocampal cortex, used as normal adult brain samples obtained after selective amygdalohippocampectomy from patients with chronic pharmaco-resistant mesial temporal lobe epilepsy and glioblastoma samples were also cut in 0.5-cm thick slices and submitted to the same histological procedure applied to fetal slices.

**Mice.** We used wild-type (WT), Nogo-A^−/−^ and S1PR2^−/−^ C57BL/6, as well as athymic nude mice. All animal experiments were approved by the Cantonal Veterinary Department of Zurich (license number 132/2014).

### In vivo mouse tumor cell culture

GL-261 tumor cells syngeneic for C57BL/6 mice were cultured in Dulbecco’s modified Eagle’s medium (DMEM; Gibco) supplemented with 10% fetal bovine serum (FBS, Gibco) and 1% penicillin/streptomycin (Invitrogen).

Human GBM cell lines G30-LRP was cultured as neurosphere in complete Neurobasal medium (ScienCell) supplemented with of human recombinant EGF (20 μg/ml, Peprotech), FGF-B (20 μg/ml, Peprotech), 2% B-27 supplement (Thermo Fisher Scientific), 0.1% GlutaMax (Thermo Fisher Scientific) and 0.1% penicillin/streptomycin (Invitrogen).

### Establishing mice tumors and in vivo imaging

#### Syngeneic tumors

Six-to ten-week-old female C57BL/6 (Charles River laboratories) were anaesthetized and immobilized on a stereotactic frame. 2×10^4^ GL-261 tumor cells in 4 μl phosphate-buffered saline (PBS) were implanted 4 mm below dura, 1.5mm lateral and 1 mm frontal from bregma in the right hemisphere.

#### Xenogeneic tumors

8-week-old female athymic nude mice (Charles River laboratories) were injected with 5×10^4^ luciferase expressing G30-LRP human tumor cells as described above for syngeneic tumors

#### In vivo imaging

Mice were injected with luciferin intraperitoneally and imaged for luciferase activity once weekly.

### Intratumoral Nogo-A Delta 20 delivery in syngeneic mice tumors

At day 14 post-implantation (DPI) of the luciferase expressing GL-261 glioma cells, the tumor-bearing animals were evenly distributed among experimental groups based on their ROI-photon flux. Osmotic pumps (model 2002, 0.5 µl/h; Alzet) were filled with Nogo-A Delta 20 (100µM in PBS) or Nogo-A Delta 21 (100µM in PBS) and primed at 37°C in PBS. Implantation of osmotic minipumps was performed as previously described (<u>vom Berg et al., 2012</u>). Briefly, the burr hole of the glioma injection was located, the bone wax and periosteal bone was removed, and the infusion cannula was inserted through the burr hole into the putative center of the tumor. Pumps were removed on day 28 post-implantation after mice euthanasia.

### Intratumoral Nogo-A Delta 20 delivery in xenogeneic mice tumors

At DPI8 of the luciferase expressing G30-LRP glioma cells, the tumor-bearing animals were randomly distributed among experimental groups Osmotic pumps (model 2002, 0.5 µl/h; Alzet) were filled with Nogo-A Delta 20 (150µM in PBS) or Nogo-A Delta 21 (150µM in PBS) and primed at 37°C in PBS. Implantation of osmotic minipumps was performed as described above. Pumps were removed at DPI22 after mice euthanasia.

### Immunofluorescence staining and analysis of postnatal brain and brain tumor angiogenesis

After perfusion with 0.1M PBS followed by 4% PFA in 0.1M PBS, P8 brains were dissected, fixed overnight in 4% PFA, frozen in Tissue Tek and 40-µm coronal free-floating sections were processed as described previously^30^. The sections were incubated for 72h at 4°C in biotinylated IB4 (Sigma, 20 µg ml^−1^) diluted in CaCl_2_-containing buffer, 0.05% Triton X-100 and 2% normal goat serum (NGS) in 0.1M PBS. After incubation with primary antibodies, the sections were washed and incubated with streptavidin-Alexa Fluor-594 conjugate (Jackson Laboratories, 1:200) diluted in CaCl_2_-containing buffer and mounted with MOWIOL.

Similarly after perfusion with FITC-lectin (Sigma), followed by perfusion with 0.1M PBS and 4% PFA in 0.1M PBS, adult glioma-bearing brains were removed, postfixed, frozen in Tissue Tek and 40-µm coronal sections were processed with a protocol adapted from^30^. Briefly sections underwent antigen retrieval by incubating the sections in 50 mM NH_4_Cl in 0.1M PB for 30 min and subsequent microwave heating in CaCl_2_-containing buffer (0.1mM CaCl_2_ 0.1mM MgCl_2_ 0.1mM MnCl TritonX-100 0.3%, diluted in 0.1M PBS, pH=6.8, slightly modified according to ^46^). After permeabilization in 0.1M tris-buffered saline (TBS) 0.3% Triton X-100 for 10 min at room temperature, the sections were incubated overnight at 4°C in primary antibodies diluted in CaCl_2_-containing buffer, 0.05% Triton X-100, 0.25% BSA IgG free (Jackson) and 0.25% Top Block (Lubioscience) in 0.1M PBS using the following primary antibodies:

rat anti-Endomucin (1:100, eBioscience), goat anti-ESM-1 (1:200, R&D systems), rabbit anti-Laminin (1:200, Sigma), rabbit anti-CA9 (1:500, Abcam), rabbit anti-GLUT1 (1:250, Millipore), rabbit anti-PDGFRβ (1:100, Abcam), goat anti-CD31 (1:500, AF3628), rabbit anti-Vimentin (1:400, Abcam), and YAP/TAZ (1:200, Cell Signaling). Sections were then incubated with secondary antibodies (goat anti-rat Alexa Fluor 568, goat anti-rabbit Alexa Fluor 568/633, donkey anti-goat Alexa Fluor 633;1:500, ThermoFisher)

Images were acquired using a Leica SP2-/SP5-, an OLYMPUS or a Zeiss confocal microscope and IMARIS software was used for 3D reconstruction of confocal z-stacks.

### Stereological analysis of mice brain tumor vasculature

Stereological analyses of vessel volume fraction, vessel length, vessel diameter, and vascular branching were performed as described previously^29^. Briefly a 140 µm x 140 µm grid ^134^ consisting of 8 µm spaced squares was superimposed on digital images of brain tumor sections perfused with FITC-Lectin, stained for Endomucin and DAPI. The relative density of blood vessels in the tissue (= vessel volume fraction) was calculated on each section by dividing the number of points falling on blood vessel structures (perfused or not perfused) by the total number of points falling on the sampling area using Stereoinvestigator. Vessel length and vessel diameter were calculated by counting the number of blood vessel structures and derived as previously described^29^.

### Isolation and culture of human brain (tumor) –derived microvascular endothelial cells (HB(T)MVECs)

The net weight of the adult control brain tissue or glioma tissue was measured followed by dissociation with collagenase/dispase (Roche) and rotation with gentleMACS C-tubes (Miltenyi biotech). Erythrocytes were lysed by resuspending and incubating the cells in ACK buffer. Cells were then counted and 20 μl of FcR blocking reagent and 20 μl of CD146 microbeads (were added per 10 cells. After 15 min rotating incubation at 4°C, the cells were passed through a 40 μ aggregates. Cells were then passed through an LS column placed in a magnetic field of a MACS separator. The CD146-positive cells remaining in the column were collected and cultured on collagen type I-coated dishes in in endothelial basal medium (EBM-2, Lonza) supplemented with endothelial growth factors EGM-2 SingleQuots (Lonza).

### Spreading assay

Spreading of HB(T)MVEC cells was determined as described previously for mouse MVECs^30^. Analysis of spreading of HBMVECs or HBTMVECs on Nogo-A Delta 20 and Nogo-A Delta 21 were performed by coating 6-well petri dishes (Greiner BioOne, well area: 1 cm^2^) with Nogo-A Delta 20 or Nogo-A Delta 21 at 1uM concentration or with PBS only. Analysis of the percentage of cell spreading was performed using an inverted microscope (OLYMPUS camedia C-7070) by determining the ratio between spread HB(T)MVECs and the total number of adhered HB(T)MVECs in three randomly chosen high power fields in each well. The percentage of cell adhesion was determined by analyzing the ratio between the number of adhered HB(T)MVECs on a given concentration of coated Nogo-A Delta 20/21 and the number of adhered HB(T)MVECs on the PBS control dish.

### Transmigration assay

The undersides of the 24-well plate inserts (FluoroBlock, BD Falcon) were coated for 1 h at 37°C with 0.5 ml PBS containing 10 µg ml^−1^ fibronectin (recombinant human fibronectin, Biopur AG) and different concentrations of Nogo-A Delta 20 and Nogo-A Delta 21, respectively. HBTMVECs were starved in migration medium (DMEM + 0.25% BSA) 4 h before detachment with accutase (PAA Laboratories). Cells were resuspended in migration medium (1 Mio ml^−1^) and labeled with calcein AM (Molecular Probes, 4 µg ml^−1^), followed by washing with HBSS and resuspending in migration medium (62,500 cells ml^−1^). The coated inserts were washed, and the bottom chamber was prepared by adding migration medium with or without mouse VEGF-A (PeproTech Inc., 10 ng ml^−1^). The inserts were then put in an empty 24-well plate (BD Falcon) and labeled HBTMVECs (25,000 per insert) were added.

The loaded inserts were transferred to the prepared cluster plate and incubated at 37°C, 5% CO_2_ for 4 h. A cell titration plate was prepared by adding different cell numbers (25,000/12,500/6,250/3,125/1,563/782 cells per well) in 1.2 ml migration medium and incubated. The fluorescence was measured after 4 h by a cytofluorometer (Millipore Cytofluor 2350) and the number of migrated cells was calculated based on the titration curve.

### Spheroid angiogenesis assay

A three-dimensional (3D) in vitro sprouting angiogenesis assay was performed as described previously^98,108^. Briefly, HB(T)MVECs were incubated overnight in hanging drops in EGM-2 medium containing 20% methylcellulose (Sigma) to form spheroids. Spheroids were then embedded in collagen gel containing 4 µM of Nogo-A Delta 20 or Nogo-A Delta 21 and cultured for 24 hours (at 37°C, 5% CO2) to induce sprouting in EGM-2 medium also supplemented with 4 µM of Nogo-A peptides. Spheroids were fixed with 4% PFA at room temperature for 15 min and images of spheroids were captured with a Leica DMi1 (objectives: 20x and 40x). Analysis of the number of sprouts and sprouts length was done using Image J.

### Nanopillar arrays

Nanopillars made of polymer (photoresistant SU8 nanopillars, Micro Chem) were fabricated using nanosphere lithography combined with a molding process, as previously described ^135,136^. Briefly, the silicon nanopillar arrays were fabricated by nanosphere lithography and plasma etching. Polydimethylsiloxane (PDMS) was used to replicate the inverse structure and SU8 polymeric nanopillars were obtained by filling SU8 liquid into PDMS molds, followed by hardening through UV exposure. The spring constant k of nanopillars was calibrated by atomic force microscopy. From the resulting force curve, k was calculated resulting in a value of 78.76 nN mm^−1^. To enable HB(T)MVEC culture on nanopillars, arrays were placed on petri dishes.

### HB(T)MVEC traction force generation and time-lapse video microscopy on nanopillar substrates

For time-lapse experiments, Vybrant Dil (Invitrogen, 1:200)-labeled HB(T)MVECs (10^4^ cells ml^−1^) were added onto the nanopillar structures. The imaging process was started 1h after seeding of the MVECs onto the nanopillars. During image processing, HB(T)MVECs were kept at constant conditions (37°C/10% CO2). The fluorescent images were acquired by using the lasers 488 nm (for nanopillar structures) and 546 nm (for labeled HB(T)MVECs) respectively.

To study acute effects of Nogo-A Delta 20 on HB(T)MVEC retraction, the nanopillar surface was coated with 25 μg ml^−1^ human fibronectin for 30 min and washed with PBS. Subsequently, DiI-labeled HB(T)MVECs were cultured for 60 min on the nanopillars and were then treated with soluble Nogo-A Delta 20 and Nogo-A Delta 21 at 1 μM, respectively. Single HB(T)MVECs were then imaged for a total time of 1.5 h, with a scanning ratio of 30 s per frame. The nanopillar displacements were recorded as sequence images when the MVECs applied forces on the nanopillar surface. For each condition, at least three individual HB(T)MVECs were analyzed.

The displacements (x) of the nanopillar tips induced by HB(T)MVECs were processed using the program Diatrack 3.03 (Powerful Particle Tracking, Semasopht). Finally, corresponding traction forces (F) were calculated using the formula from Hooke’s law: F=k*x (k = spring constant of the nanopillars = 78 nN/µm).

### Cell immunofluorescence

Cells were cultured on an 8-well chamber slides (Sigma-Aldrich), cells were fixed with 4% PFA (Sigma-Aldrich) for 15 min at room temperature (RT). The cell membrane was then permeabilized with 0.1% Triton X-100 (X100, Sigma-Aldrich) in PBS (10 min) followed by 1% BSA/PBS (85040C, Sigma-Aldrich) blocking step (30 min, RT).

HUVECs were stained with rabbit anti-VEGFR2 (1:100, Santa Cruz cat. sc-55486), rabbit anti-phospho-VEGFR2 (1:100, abcam cat. ab127894) in 1% BSA/PBS at 4°C overnight. Cells were the incubated with secondary antibodies goat anti-rabbit Alexa 568 and goat anti-mouse Alexa 488 (1:500, Thermo Fisher Scientific cat. A-11011 and A-11001), or TRITC-labeled phalloidin (1:100 Sigma) in 1% BSA/PBS for 1.5 hours at RT. Nuclei were counterstained with DAPI staining (1:20,000, Thermo Fisher Scientific) in PBS for 5 min.

### Bulk RNA-sequencing and analysis

RNA extraction and bulk RNAseq was performed as previously described^108^. Total RNA was extracted using the RNeasy RNA isolation kit (Qiagen, Hilden, Germany) including a DNase treatment to digest residual genomic DN. RNA sequencing of endothelial cells was performed by the Functional Genomics Center Zurich. Libraries preparation was performed following Illumina TruSeq stranded mRNA protocol. RNA and final libraries quality were addressed using an Agilent 4200 TapeStation System. The libraries were pooled equimolarly and sequenced in an Illumina NovaSeq sequencer (single-end 100 bp) with a depth of around 20 Mio reads per sample. For mapping and trimming of FASTQ format sequences was performed using Trimmomatic v0.3.3, and sequence quality control was assessed using FastQC. Alignment to the Ensembl Homo_sapiens GRCh38.p10 reference genome (Release_91-2018-02-26) was performed using the STAR aligner. Gene expression values were computed with the function featureCounts from the R package Rsubread. Differential expression was computed using the generalized linear model implemented in the Bioconductor package DESeq2. Statistical analysis included Wald test followed by correction for multiple testing using the Benjamini–Hochberg method (False Discovery Rate, FDR).

Pathway analysis was performed on the ranked differential expression gene list (rank formula:“*-LOG10(pvalue)*SIGN(logFC)*”) using the Gene Set Enrichment Analysis (GSEA) software from the Broad Institute (software.broadinstitute.org/GSEA) (version 4.0.1)^137,138^.

“Human_GOBP_AllPathways_no_GO_iea_November_01_2021.gmt” from [http://baderlab.org/GeneSets] was used to identify enriched pathways in GSEA analysis (inclusion criteria are pathways with a minimum size of 15 and a maximum size of 300 genes). The resulting pathways were filtered based on passing the threshold of FDR<0.05 and pvalue<0.05. Significantly regulated pathways were plotted using Cytoscape (Version 3.7.0) and EnrichmentMap (version 3.3)^139^. Related pathways were grouped into themes, labeled by AutoAnnotate (version 1.3) and manually curated.

### Immunoblotting

Cells were lysed in RIPA buffer and mechanical disruption through a 1ml insulin syringe (BD). Proteins were resolved by 10% SDS-polyacrylamide gel electrophoresis (SDS-PAGE) at 150 V and transferred to a nitrocellulose membrane at 100 V for 1.5 hours. After blocking with TBST 5% skim milk, the membrane was probed with rabbit anti-YAP/TAZ ( 1:100, Cell Signaling), rabbit anti-Phospho-YAP (1:1,000, Cell Signaling), rabbit anti-LATS1 (1:500, Cell Signaling), rabbit anti-pLATS1 (1:500, Cell Signaling), rabbit anti-LATS2 (1:500, Cell Signaling), rabbit anti-NOTCH (1:500, abcam), rabbit anti-NICD (1:400, abcam), rabbit anti-VEGFR2 (1:1,000; Cell signaling) and rabbit anti-phospho-VEGFR2 (1:1,000; Cell signaling) at 4°C overnight. A secondary For the generation of HUVEC lysates for VEGFR2 immunoblotting, HUVECs were starved overnight in serum-free medium. The next day, HUVECs were stimulated with soluble Nogo-A Delta 20 or Nogo-A Delta 21 at a final concentration of 1 µM as well as with or without soluble VEGF-A at a concentration of 50µg/ml for 5, 10, and 30 min prior to the generation of MVEC lysates.

Bands were visualized by chemiluminescence using ECL (Thermo Fisher, cat. 32132). Densitometric analysis was performed with ImageJ (NIH freeware). Data were normalized to actin, and values of control cells were set to 1.

### Quantitative real-time PCR

Total RNA was prepared using the RNeasy RNA isolation kit (Qiagen, Hilden, Germany) including a DNase treatment to digest residual genomic DNA. mRNA was reverse-transcribed using iScript cDNA synthesis kit (Bio-Rad). Real-time qPCR analysis was performed using a SYBR Green-based master mix (ThermoFisher Scientific). Relative quantification was calculated using the comparative threshold cycle (ΔΔ^CT^) method. cDNA levels were normalized to *S18* (reference genes), and a control sample (calibrator set to 1) was used to calculate the relative values.

### LC-MS/MS analysis polar metabolites

Sample preparation for LC-MS/MS analysis were modified from Paglia et al., 2014^100^ as described previously (Schwab et al., JCI Insight). HB(T)MVECs were washed twice with PBS and fixed with 80% methanol. HB(T)MVECs were detached using a cell scraper and lyzed mechanically using a dounce homogenizer. Extracts were centrifuged for 20min at 10,000g and 4°C. Samples were stored at –20°C until LC-MS/MS analysis.

50 µl methanol extract was dried under a N2, reconstituted in 20 µl water and diluted with 80 µl injection buffer (90% acetonitrile, 8.8% methanol, 50 mM NH4-acetate). Samples were vortexed and centrifuged (10,000g, 4°C, 15 min). 50 µl of the supernatant was transferred to a glass vial (Total Recovery Vials, Waters, Milford, MA, USA) for LC-MS/MS injection.

Metabolites were separated on a nanoAcquity UPLC (Waters, Milford, MA, USA) equipped with a BEH Amide capillary column (150 µm × 130 mm, 1.7 µm particle size, Waters, Milford, MA, USA). Solvent system consists of buffer A (5 mM NH4-acetate in water) and buffer B (5 mM NH_4_-acetate, in 95% acetonitrile). Linear gradient applied from 10% A to 90% A over 10 min, with flow rate ramped down from 3 µl/min to 2 µl/min. Washing for 1 min at 50% A with a flow rate of 2 µl/min, followed by 5 min re-equilibration with flow rate ramped to 3 µl/min. Injection volume was 1 µl. The UPLC was coupled to Synapt G2-Si mass spectrometer (Waters, Milford, MA, USA) by a nanoESI source. MS1 and MS2 data was acquired using negative polarization and MSE over a mass range of 50 to 1,200 m/z at resolution of >20,000.

Data were aligned and searched against databases with the Progenesis QI software (Waters, Milford, MA, USA). Polar metabolites and lipids were searched against the KEGG database with a precursor mass tolerance of 20 ppm and fragment mass tolerance of 50 ppm. Quality controls were run on pooled samples and reference compound mixtures to verify technical accuracy and stability. Reference compound included key metabolites from glycolysis, oxidative phosphorylation, TCA cycle, pentose phosphate, as well as amino acids and nucleotides. Those metabolites were manually curated in Skyline.

MetaboAnalyst 5.0 was used The peak intensity (peaks(mz/rt)) table was uploaded as a data matrix file to MetaboAnalyst 5.0^140^ for reading, processing the raw data and for statistical analysis. Statistical analysis performed include univariant analysis (fold change analysis and t-test) to plot the volcano plots. Features that passed the threshold of fold change –1.5<x<1.5 and p-value < 0.05 were counted as significantly changed, red dots indicate upregulated, while blue dots indicate down regulated. Multivariant analysis performed include unsupervised principal component analysis (PCA) to plot 2D and 3D PCA plots was done using prcomp package. Clustered heatmaps were plotted using the PlotHeatMap function implemented from the R pheatmap package (distance measure using euclidean, and clustering algorithm using ward.D).

### Glycolytic Flux

HB(T)MVECs were incubated for 6 hours in EGM-2 containing 0.4 μCi/ml [5-3H]-D-glucose (PerkinElmer). Supernatant was transferred into glass vials containing perchloric acid and sealed with rubber stoppers. ^3^H_2_O was captured in hanging wells containing filter paper soaked with H_2_O over a period of 48 hours at 37°C. Thereafter, the filter paper was transferred in scintillation cocktail for radioactivity measurement by liquid scintillation counting.

### Fatty acid oxidation

HB(T)MVECs were incubated in FBS-free EGM-2 medium supplemented with 50 fatty-acid free BSA, 50 μ carnitine, 100 μ unlabeled palmitic acid, and 2 μCi/ml [9,10-acid. Again, supernatant was transferred into glass vials and sealed with rubber stoppers. Radioactivity was measured as in the glycolytic flux assay.

### Statistical analysis

Statistical significance was determined using unpaired two-tailed Student’s t-test, one-way or two-way ANOVAs (GraphPad Prism8). Differences were considered significant with a P value less than 0.05. Quantified data are presented as mean ± SEM.

## SUPPLEMENTARY FIGURE LEGENDS

**Supplementary Figure S1.**
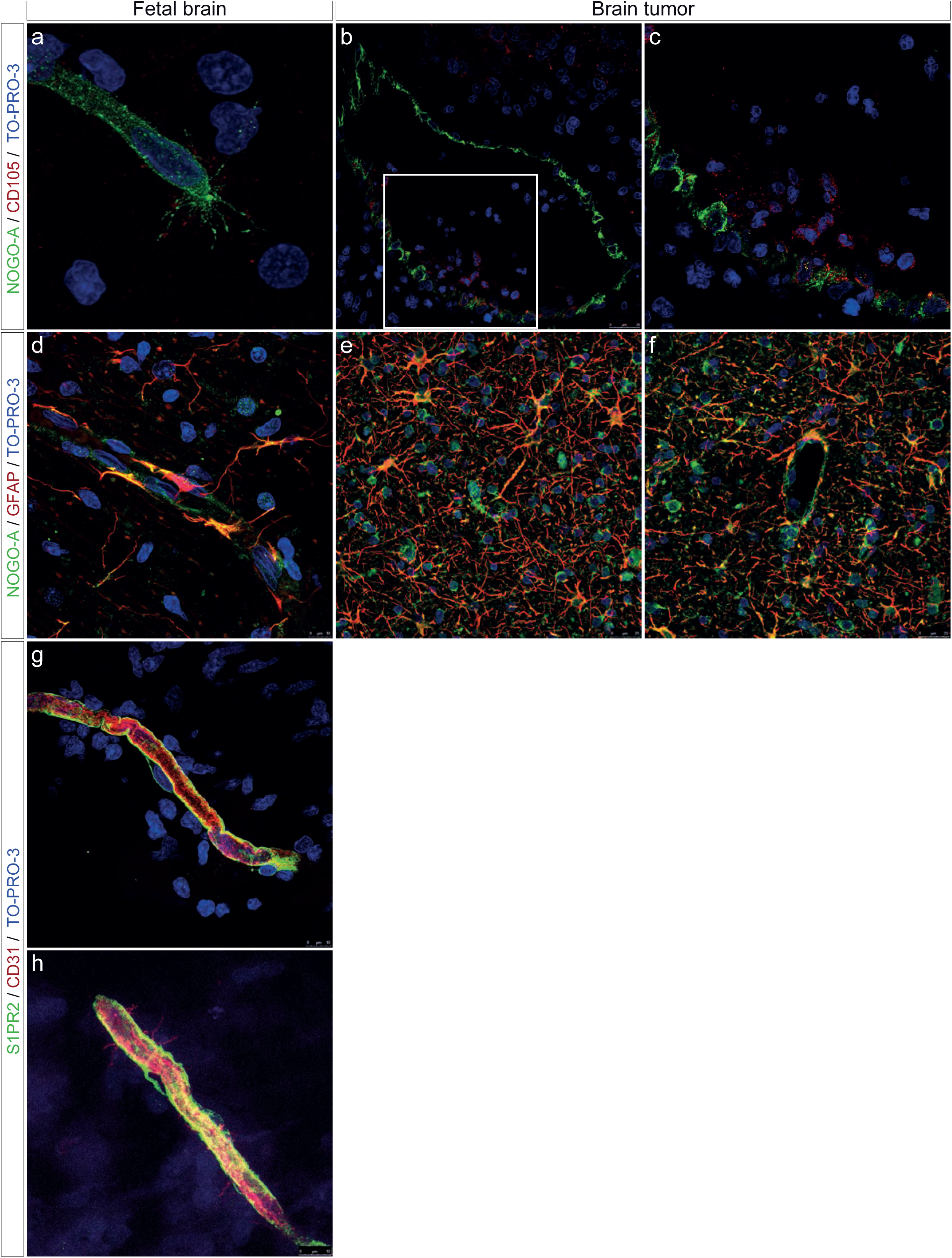
Nogo-A is expressed in the vicinity of endothelial (tip) cells and in perivascular astrocytes during human brain development and in glial brain tumors in vivo. Coronal sections (20 µm) of human fetal (GW 122) and adult human glial brain tumors (GBMs) were stained for Nogo-A (red), the vascular endothelial cell markers CD31 (-endothelial (tip) cell, red) and CD105 (endoglin, endothelial (tip) cells, green), the astrocytic marker GFAP (green), Nogo-A receptor S1PR2 (green) and TO-PRO-3 nuclear counterstaining (blue). **(a-c)** Nogo-A (red) is expressed in the vicinity of the endothelial tip cell in human fetal brain (**a**) and of human brain tumors endothelial cells (**b-c**). **(d-f)** Nogo-A (red) is highly expressed in GFAP^+^ neural precursors cells (green) in the fetal brain (**d**) and in tumoral astrocytes in glioblastoma (**e-f**). **(g-h)** S1PR2 (green) is highly expressed in CD31^+^ blood vessel endothelial (tip) cells (red) in the human fetal (**g**,**h**). Scale bars: 10 μm in **a**,**d**,**g**,**h**, and 25 µm in **b**-**c**,**e**-**f**.

**Supplementary Figure S2.**
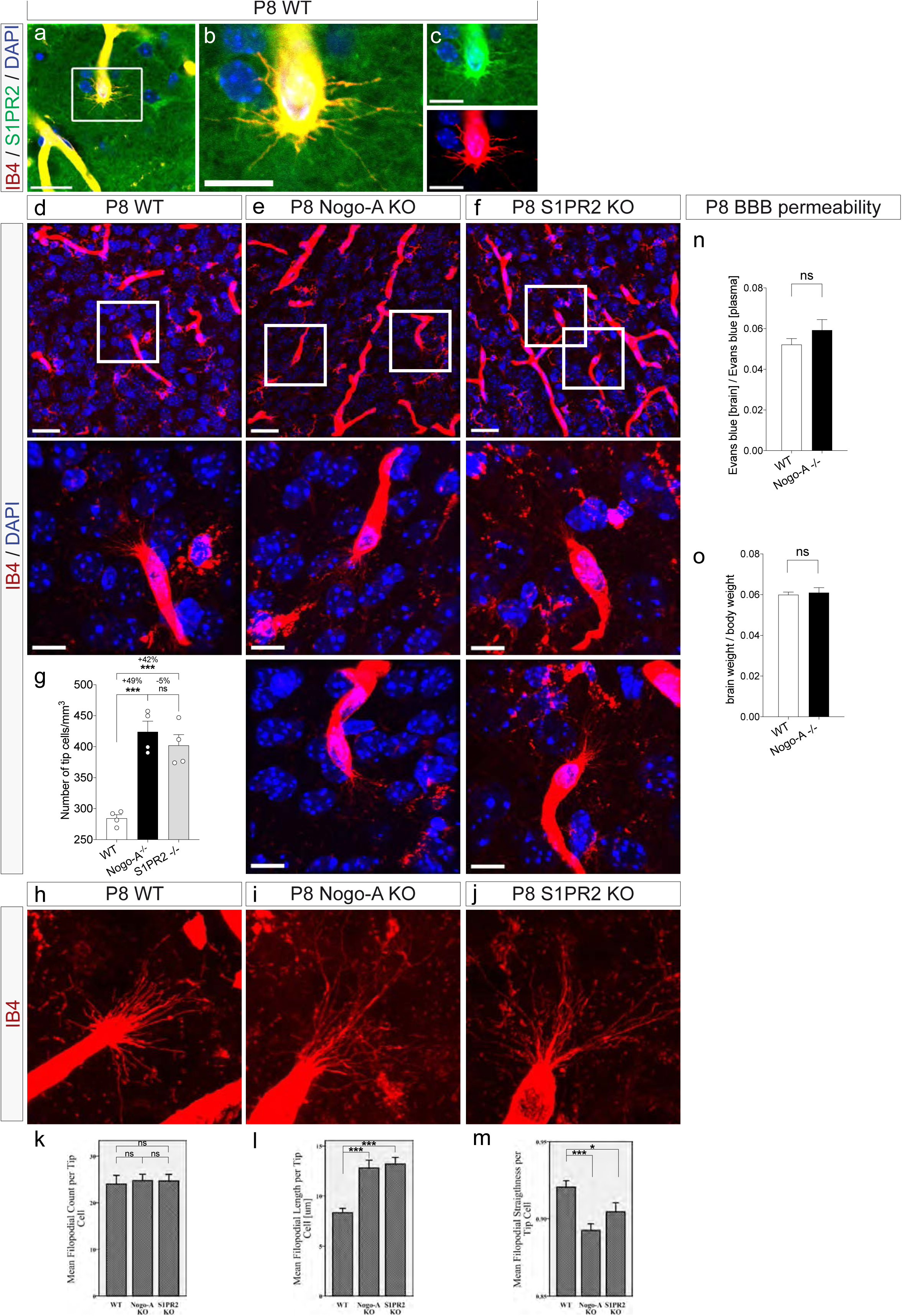
S1PR2 is expressed by endothelial (tip) cells during mouse brain development and Nogo-A/S1PR2 regulate tip cell filopodia structure but not BBB integrity. (**a-m**) Coronal sections of postnatal day 8 (P8) mouse brains were stained for the endothelial marker IB4 (red), S1PR2 (green) and cell nuclei (DAPI, blue). S1PR2 (green) is expressed on IB4-labeled endothelial tip cells and its filopodia (**a**-**c**). (**d**-**g**) The number of IB4^+^ endothelial tip cells was significantly increased in the cortices of Nogo-A KO mice (**e**) and of S1PR2 KO mice (**e**) as compared to WT mice (**d**,**g**). Boxed areas are enlarged below. (**h**-**m**) Number of IB4^+^ endothelial cell filopodia was not affected by genetic deletion of Nogo-A or its receptor S1PR2 (**k**) but Nogo-A KO and S1PR2 KO showed increased filopodia length (**l**) and decreased filopodia straightness (**m**, n=20). (**n**-**o**) P8 WT, Nogo-A KO and S1PR2 KO mice were perfused with Evans blue prior euthanasia. Brains were extracted, weighted and Evans blue was measured in brain tissue and in blood (plasma). Evans blue diffusion in the brain through the blood brain barrier (BBB) was not affected by genetic deletion of Nogo-A (**n**), indicating that Nogo-A doesn’t regulate BBB permeability. Nogo-A KO doesn’t affect brain size, a potential confounding factor in this analysis. Data represent mean ± SEM. For statistical analysis, two-tailed unpaired Student’s t –test (**g**,**k**-**o**) were performed. \**P* < 0.05, \*\*\**P* < 0.001. Scale bars represent: 30 µm (**d**-**f**) 10 µm (**a**-**c**, and insets in **d**-**f**).

**Supplementary Figure S3:**
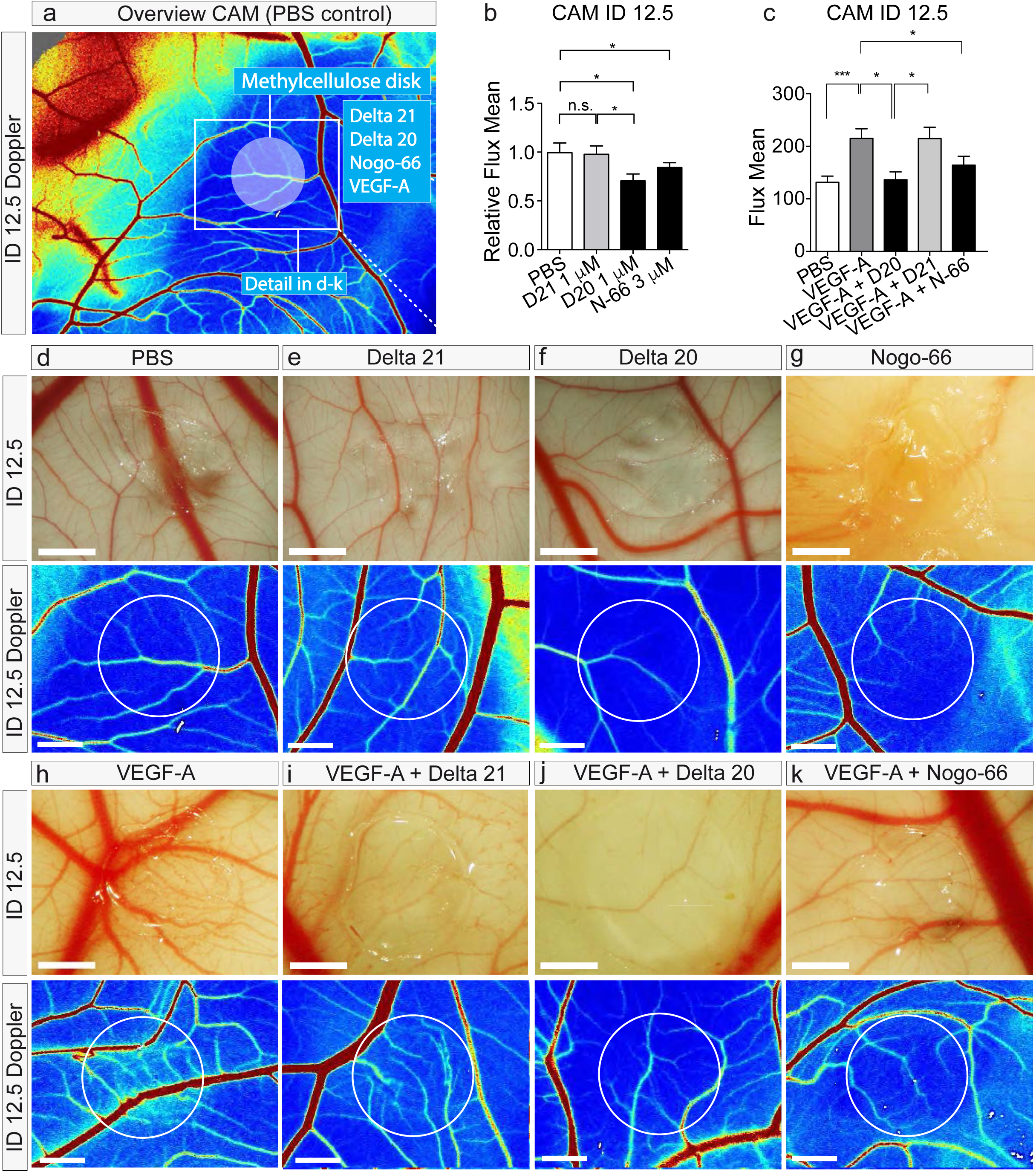
Nogo-A Delta 20 and Nogo-66 exert local inhibitory effects on in vivo CAM angiogenesis and counteract the pro-angiogenic effects of VEGF-A. (**a**) Overview of control CAM blood vessels treated with PBS at ID12.5. Methylcellulose disks (MC disks) soaked with proteins were applied as indicated by the semi-opaque circle. Red colors indicate high blood flow values, blue colors indicate low blood flow values. (**b**) Quantification of Doppler measurements relative to PBS control. Inhibition of blood vessel density and significant reduction of local blood flow (blue) in Nogo-A-Delta 20-treated CAMs at ID12.5 as compared to Nogo-A Delta 21-treated or PBS-treated CAMs. A strong induction of vascular density and flow was seen after addition of VEGF-A (n ≥ 3). (**c**) Quantification of Doppler measurements. The strong induction of local vascular density and local blood flow seen after addition of 1 μg VEGF-A could be counterbalanced by concomitant application of Nogo-A Delta 20 but not Nogo-A Delta 21 (n ≥ 3). (**d**-**k**) Representative images of different conditions displayed as incident light microscopy– and Doppler images, with VEGF-A stimulation (**h**-**k**) and without VEGF-A stimulation (**d**-**g**). A reduction of local blood vessel density and local blood flow values could be observed in the Nogo-A Delta 20 (1 μM, **f**) and Nogo-66 (3 μM, **g**) treated CAMs as compared to Nogo-A Delta 21 (1 μ **e**) and PBS (**d**) control treatments. Note the dark blue color (corresponding to a reduction in local blood vessel density– and flow) between larger vessels in the Nogo-A Delta 20 (**f**, lower panel) and Nogo-66-treated CAMs (**g**, lower panel), where usually a dense network of capillaries is seen in controls (**d**,**e**, lower panel). Application of 1 μg VEGF-A had a strong pro-angiogenic effect that was clearly visible in incident light microscopy-as well as in Doppler images (**h**). Nogo-A Delta 20 (**j**) and Nogo-66 (**k**) were capable of attenuating the strong pro-angiogenic effects of VEGF-A. Nogo-A Delta 21 had no effect on the pro-angiogenic effects of VEGF-A (**i**). Data represent mean ± SEM. For statistical analysis, two-tailed unpaired Student’s t –test (**b**,**c**) were performed. \**P* < 0.05, \*\*\**P* < 0.001. Scale bars represent: 2 mm (**d**-**k**).

**Supplementary Figure S4:**
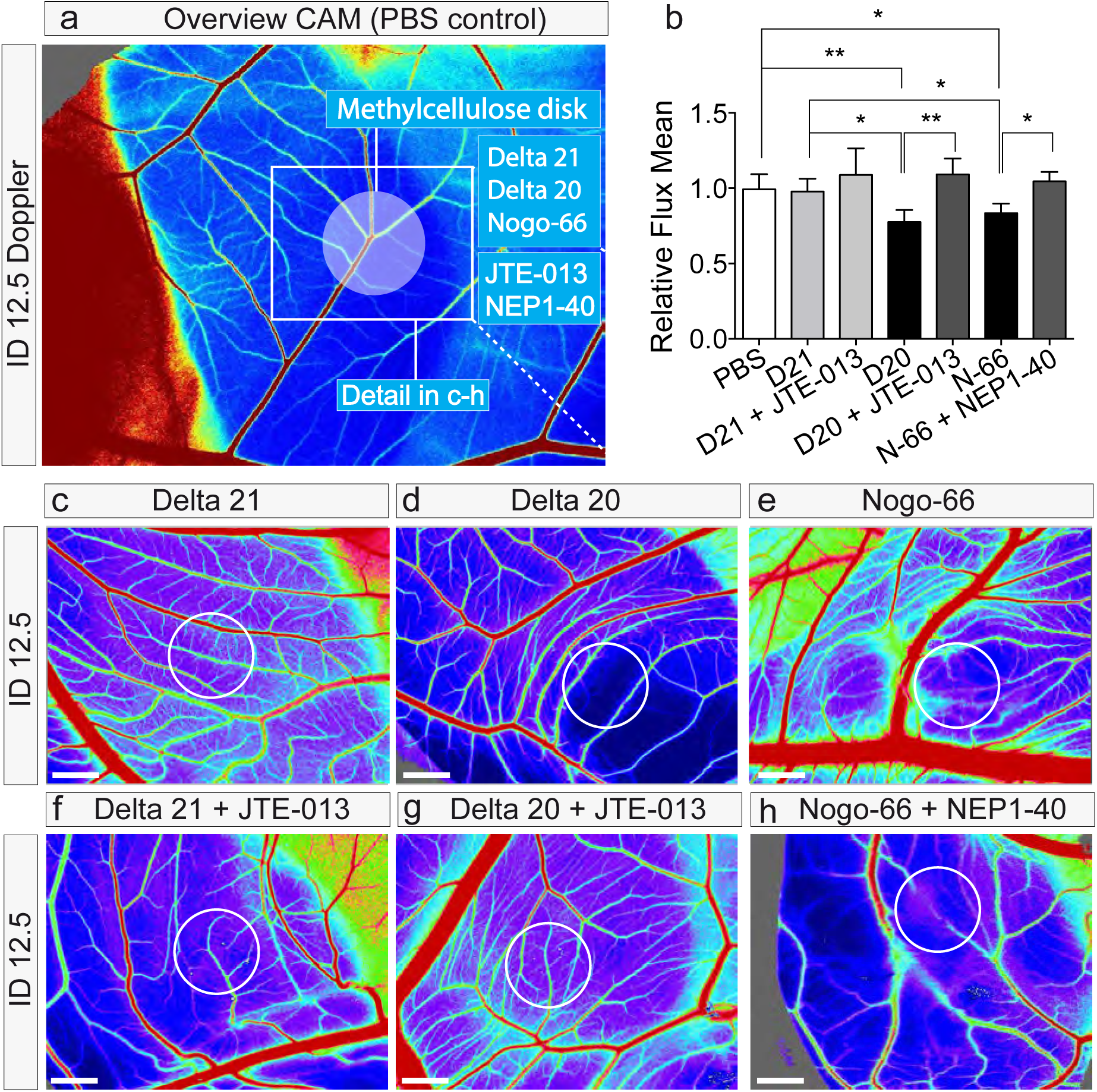
Blocking Nogo-A receptor components (S1PR2, NgR1) partially prevents the Nogo-A-induced local inhibitory effects on in vivo CAM angiogenesis. (**a**) Overview of control CAM blood vessels treated with PBS at ID12.5. Methylcellulose disks (MC disk) soaked with proteins and pharmacological blockers were applied as indicated by the semi-opaque circle. Red colors indicate high blood flow values, blue colors indicate low blood flow values. (**b**) Quantification of Doppler measurements relative to PBS control. The inhibition of blood vessel density and blood flow in Nogo-A-Delta 20-treated CAMs at ID12.5 was prevented by addition of the S1PR2-specific inhibitor JTE-013 (red, 0.4 µg per MC disk). Similarly, the inhibitory effect of Nogo-66 was partially counterbalanced by addition of NEP1-40 (red, 4.6 μg per MC disk), a competitive antagonist of NgR1. (**c**-**h**) Representative images of different drug treatments displayed as incident light and Doppler images, with (**f**-**h**) and without (**c**-**e**) addition of Nogo-A receptor blockers. The inhibitory effects of Nogo-A Delta 20 (**d**) and of Nogo-66 (**e**) could partially be prevented by daily application of the S1PR2-blocker JTE-013 (**g**) and of the competitive antagonist of NgR1, NEP1-40 (**h**). JTE-013 had no effect on blood vessel density and flow values in combination with the control peptide Nogo-A Delta 21 (**c**,**f**). Data represent mean ± SEM. For statistical analysis, two-tailed unpaired Student’s t –test (**b**,**c**) were performed. \**P* < 0.05, \*\**P* < 0.01. Scale bars represent: 2 mm (d-k).

**Supplementary Figure S5:**
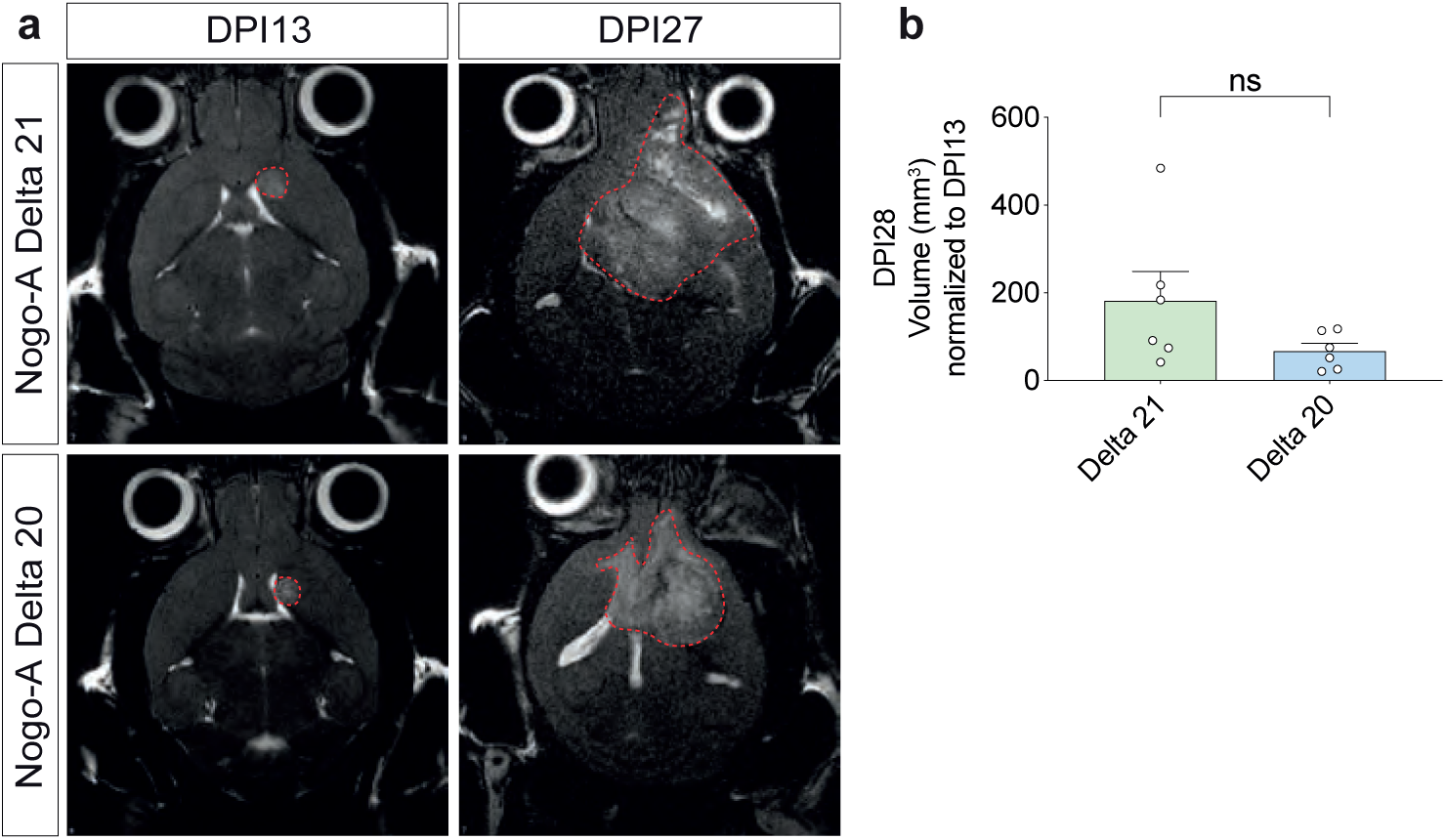
Local infusion of Nogo-A Delta 20 inhibits tumor growth as revealed by MRI. (**a**,**b**) T2 weighted (T2w) MRI imaging of mice one day before Nogo-A Delta 21 and Nogo-A Delta 20 treatments as well as at the endpoint of the experiment (**a_i_**, red dashed lines surround the tumors). Tumor volume at DPI28 (normalized to volume at DPI14) tend to decrease in Nogo-A Delta 20 treated mice as compared to Nogo-A Delta 21 control (**b**).

**Supplementary Figure S6.**
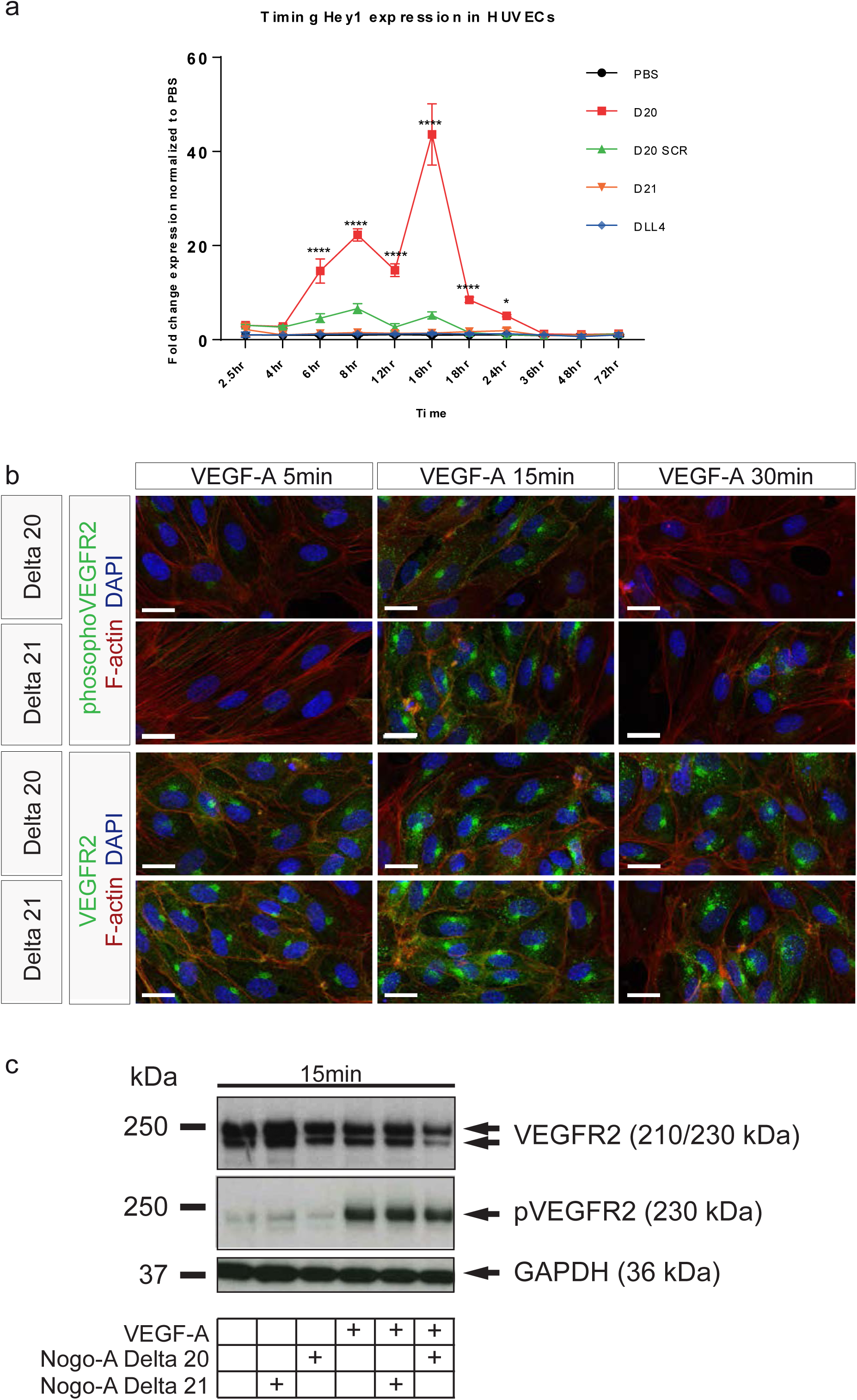
Nogo-A Delta 20 signaling interacts with the VEGF-VEGFR2 pathway. (**a**) HUVECs were treated with VEGF-A for 5min, 15min, or 30min and Nogo-A Delta 20 or Nogo-A Delta 21. Phosphorylation of VEGR2 at Tyr1175 and total VEGFR2 was visualized with an immunofluorescent antibody staining. VEGFR2 phosphorylation peaked after 15min and was considerably lower in the Nogo-A Delta 20-treated conditions as compared to Nogo-A Delta 21 conditions. While in the Nogo-A Delta 21-treated conditions there was residual phosphoVEGFR2 after 30min, no phosphoVEGFR2 was detectable in the Nogo-A Delta 20 conditions. (**b**) HUVECs were treated with or without VEGF-A for 15min and Nogo-A Delta 20 or Nogo-A Delta 21. Phosphorylation of VEGR2 at Tyr1175 and total VEGFR2 was visualized with western blot. Treatment with Nogo-A Delta 20 or Nogo-A Delta 21 alone did not induce VEGFR2 phosphorylation. Addition of VEGF-A strongly induced phosphorylation of VEGFR2 at Tyr1175. Addition of Nogo-A Delta 20 reduced the amount of phosphoVEGFR2 protein as well as the total amount of VEGFR2 protein. Scale bars represent: 20 µm.

**Supplementary Figure S7:** Endogenous expression of Nogo-A inversely correlates with vessel density during tumor progression of human astrocytomas. (**a**-**f**) Nogo-A (red) expression increases during tumor progression of human astrocytomas. (**g**) Analysis of CD31^+^ vessel volume fraction in low Nogo-A expressing (H-score 0-99) and high Nogo-A expressing (H-score 100-300) adult control brain, low-grade glioma and high-grade glioma samples. Adult control brain didn’t show high expressing samples. CD31^+^ vessel volume fraction was significantly higher in high Nogo-A expressing WHO II glioma, but tend to decrease in WHO II glioma and higher-grade glioma, (**g**,**j**). Nogo-A expression displays a high variability in human astrocytomas WHO IV (glioblastoma), ranging from low expression (**h**) to very high expression (**i**). (**l**) The number of CD31^+^ small caliber and large caliber vessels was not affected by Nogo-A expression in glioblastoma. Data represent mean ± SEM. \**P* < 0.05, \*\*\**P* < 0.001. Scale bars represent: 100 µm in **a**-**f** (upper pictures) and in **i**-**j** (upper pictures); 20 µm in **a**-**f** (lower pictures) and in **j-k** (lower pictures)

